# Sequential *in vitro* enzymatic N-glycoprotein modification reveals site-specific rates of glycoenzyme processing

**DOI:** 10.1101/2022.07.27.501726

**Authors:** Trevor M. Adams, Peng Zhao, Digantkumar Chapla, Kelley W. Moremen, Lance Wells

## Abstract

N-glycosylation is an essential eukaryotic post-translational modification that affects various glycoprotein properties, including folding, solubility, protein-protein interactions, and half-life. N-glycans are processed in the secretory pathway to form varied ensembles of structures, and diversity at a single site on a glycoprotein is termed ‘microheterogeneity’. To understand the factors that influence glycan microheterogeneity, we hypothesized that local steric and electrostatic factors surrounding each site influences glycan availability to enzymatic modification. We tested this hypothesis by expression of a panel of reporter N-linked glycoproteins in *MGAT1-* null HEK293 cells to produce immature Man_5_GlcNAc_2_ glycoforms (38 glycan sites total). These glycoproteins were then sequentially modified *in vitro* from high-mannose to hybrid and on to biantennary, core fucosylated, complex structures by a panel of N-glycosylation enzymes and each reaction time-course was quantified by LC-MS/MS. Substantial differences in rates of *in vitro* enzymatic modification were observed between glycan sites on the same protein and differences in modification rates varied depending on the glycoenzyme being evaluated. By comparison, proteolytic digestion of the reporters prior to N-glycan processing eliminated differences in *in vitro* enzymatic modification. Comparison of *in vitro* rates of enzymatic modification with the glycan structures found on the mature reporters expressed in wild type cells correlate well with the enzymatic bottlenecks found *in vitro*. These data suggest that higher-order local structures surrounding each glycosylation site contribute to the efficiency of modification both *in vitro* and *in vivo* to establish the spectrum of site-specific microheterogeneity found on N-linked glycoproteins.

## Introduction

Glycans are important modulators of protein properties and functions across all clades of life (1, 2). N-glycosylation is a conserved and essential co- and post-translational modification in higher eukaryotes (3) and plays an important role in protein homeostasis (4). N-glycans are co- and/or post-translationally attached *en bloc* by oligosaccharyltransferase (5) to the conserved motif N-X-S/T(C), also known as a “sequon”, where ‘X’ can be any amino acid except proline (6, 7). The initial oligosaccharide is mannose-rich and it is trimmed by a series of glycoside hydrolases in the endoplasmic reticulum (ER), eventually exposing the core structure of N-glycans, which is made up of a chitobiose core (GlcNAc-GlcNAc) with branching mannoses, Man_5_GlcNAc_2_. N-glycans are generally categorized as belonging to one of three classes based on the extent of their processing: high-mannose, hybrid, or complex. High-mannose glycans are the least processed and most closely resemble the initial oligosaccharide that is transferred onto proteins, while complex glycans are the most processed and can take a variety of forms. This can include branching, extensions, and core fucosylation (8). However, the efficiency of glycan maturation at a given acceptor site on glycoproteins can often be incomplete, most notably because of steric or electrostatic factors that impact enzyme-substrate recognition (8–10). This often results in heterogeneous ensembles of glycan structures on glycoprotein acceptor (11), and even on individual glycosites on the same glycoprotein (12). This phenomenon is termed ‘microheterogeneity’ and is a hallmark of protein glycosylation.

An important branching point in N-glycan processing is the addition of a branching β-2-linked GlcNAc to the α3 mannose of the Man_5_GlcNAc_2_ structure by the GT-A family glycosyltransferase MGAT1 (13, 14). This step marks a class switch from high-mannose (e.g. Man_5_GlcNAc_2_) to hybrid (e.g. GlcNAcMan_5_GlcNAc_2_) glycans, as this branching GlcNAc can be further elaborated by other glycosyltransferases into a variety of structures. The activity of MGAT1 is also necessary for the subsequent trimming of the two terminal mannoses from the α6 mannose by the glycoside hydrolase MAN2A1 (11), which results in a GlcNAcMan_3_GlcNAc_2_ structure. This GlcNAcMan_3_GlcNAc_2_structure acts as the substrate for MGAT2 (12), which marks the point at which hybrid N-glycans transition to complex N-glycans, with two branching GlcNAc moieties that serve as a base for highly elaborated bi-, tri-, or tetra-antennary structures.

There is much interest in understanding the underlying criteria that define N-glycan microheterogeneity. N-glycans have been shown to be important modulators in antibody-receptor interactions, both with respect to glycosylation of the antibody (15–17) and their receptors (18, 19). In particular, the contribution of glycosylation to the properties of therapeutics is of particular interest in the development and manufacturing of biologics (4, 20) and biosimilars (21). Glycosylation of these therapeutics is known to impact their stability and pharmacokinetics (4). Additionally, N-glycans are a vital component of viral glycoprotein properties and are known to impact host immune surveillance (22, 23) and host receptor interactions involved in viral entry (24).

There have been several approaches to studying N-glycan microheterogeneity. Early studies demonstrated that N-glycan microheterogeneity is reproducible on a site-by-site basis (25). NMR studies of N-glycan structures suggest that glycan interactions with the protein backbone can alter glycan conformations in ways that can impact N-glycan processing (26–28). There is evidence that changing nearby amino acids can alter N-glycan heterogeneity (9, 29). Some recent studies have taken a systems approach, often by monitoring *in vivo* processing of N-glycans in cell culture systems (9, 10). The involvement of the peptide-glycan interactions in affecting glycan conformation, and thus potentially N-glycan processing, has also been supported by molecular dynamics simulations using yeast protein disulfide isomerase (PDI) as a model reporter glycoprotein (30, 31). A recent study by Mathew et al. studied early N-glycan processing steps through molecular dynamics simulations and *in vitro* processing of N-glycans on the yeast protein disulfide isomerase, a similar approach as this study (32). They found that the shape of the surrounding protein environment and subsequent glycan conformation can influence the rate at which individual sites are processed. They studied early mannose trimming and class switching from high-mannose to hybrid glycans using three mannosidases and the GlcNAc transferase MGAT1 on the processing of PDI. Our work here expands on this by using enzymes involved in later N-glycan processing using not only the model yeast protein PDI but also multiple N-linked glycoproteins of interest to human health.

In this study, we report extensive site-specific *in vitro* N-glycan processing data for five multiply N-linked glycosylated proteins, with 38 different sites of N-glycosylation in total. By enriching all sites of all glycoproteins with a common Man_5_GlcNAc_2_ substrate and then monitoring N-glycan processing through time-course reactions, we were able to identify key bottlenecks that prevent specific sites on glycoproteins from being converted from high-mannose to complex N-glycans. These bottlenecks appear to persist *in vivo* upon microheterogeneity analysis of each site of the reporter proteins when expressed in wild type cells. Additionally, we found that removing the tertiary structure of the protein abolished all site-specificity of N-glycan processing, highlighting the importance of protein tertiary structure in defining N-glycan microheterogeneity.

## Results

### Expression of reporter proteins in WT-HEK293F cells

In order to probe individual steps of N-glycan processing, we first established a set of reporter proteins to be used as case studies (**Table 1**, **Fig**. S1). These proteins were selected based on their various applications in biology, virology, and use as therapeutics as well as their diversity in displayed glycans and the availability of quality crystal structures. CD16a (Fc γ receptor IIIa) is an IgG receptor that is known to have differential affinities to antibodies depending on its glycan presentation which impacts downstream signaling (19, 29, 33). Protein disulfide isomerase (PDI) is a resident ER glycoprotein that has been used as a model protein for studying N-glycan processing due to its ease of expression, analysis, and well-defined site-specific glycan heterogeneity (9, 10, 30–32). Etanercept is a bioengineered therapeutic fusion protein of a TNFα receptor and an IgG1 Fc domain commonly used to help treat auto-immune disorders (34). Erythropoietin is a therapeutic glycoprotein that stimulates red-blood cell growth, and its glycosylation is known to impact its pharmacokinetics (35–37). SARS-CoV-2 spike glycoprotein is a highly glycosylated trimer that is responsible for the viral entry of the associated coronavirus SARS-CoV-2 via binding to the human receptor ACE2 (24, 38).

**Table 1:**
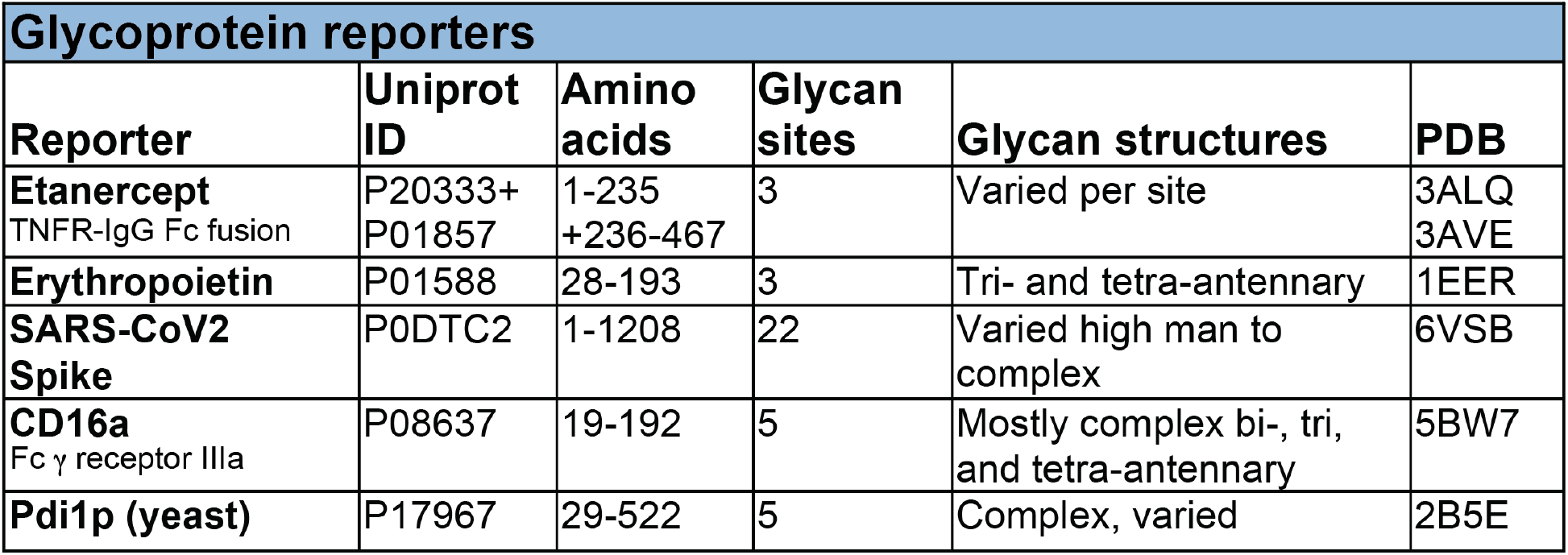
Reporter proteins used as models for studying N-glycan processing

The reporter proteins were first transiently expressed in high yields in wild type HEK293F cells then harvested from supernatant and purified with Ni-NTA chromatography (**Figs. S1b-f)**. Glycopeptide analysis using LC-MS/MS was performed on these purified proteins in order to determine their glycan occupancy and diversity when expressed in a “wild type” background (**Fig. 1**). With liquid chromatography and tandem mass spectrometry techniques, we were able to obtain a detailed characterization of the N-glycan profile at each site on the reporter proteins, some of which contained dozens of different glycan moieties with a variety of terminal structures including sialylation, as well as core fucosylation (**Fig. 2a-c**).

**Figure 1:**
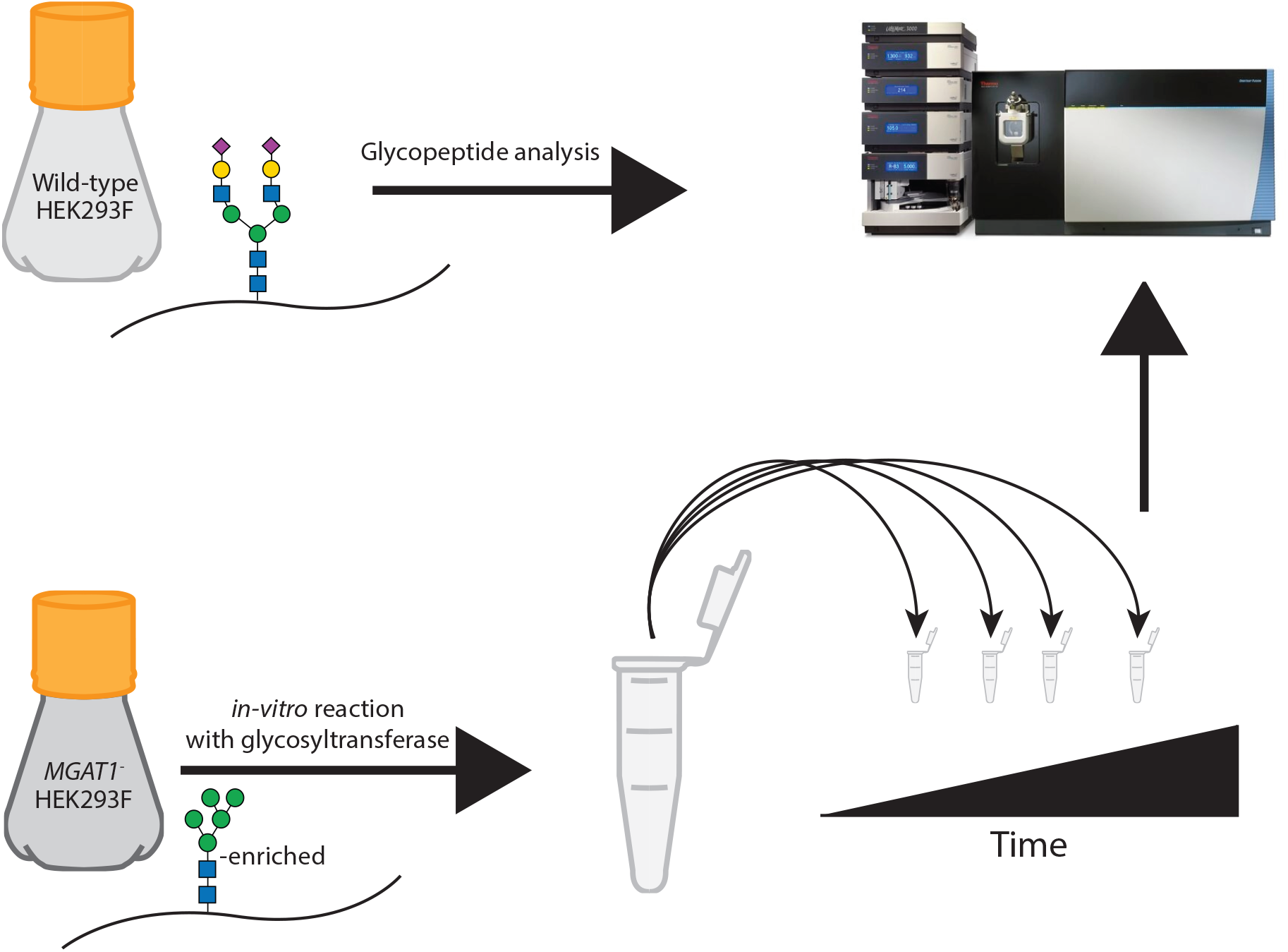
Graphical representation of approach. Reporter proteins were expressed in HEK293F WT and MGAT1-cells, analyzed via LC-MS/MS, and then processed by purified glycosyltransferases and hydrolases *in vitro*.

**Figure 2:**
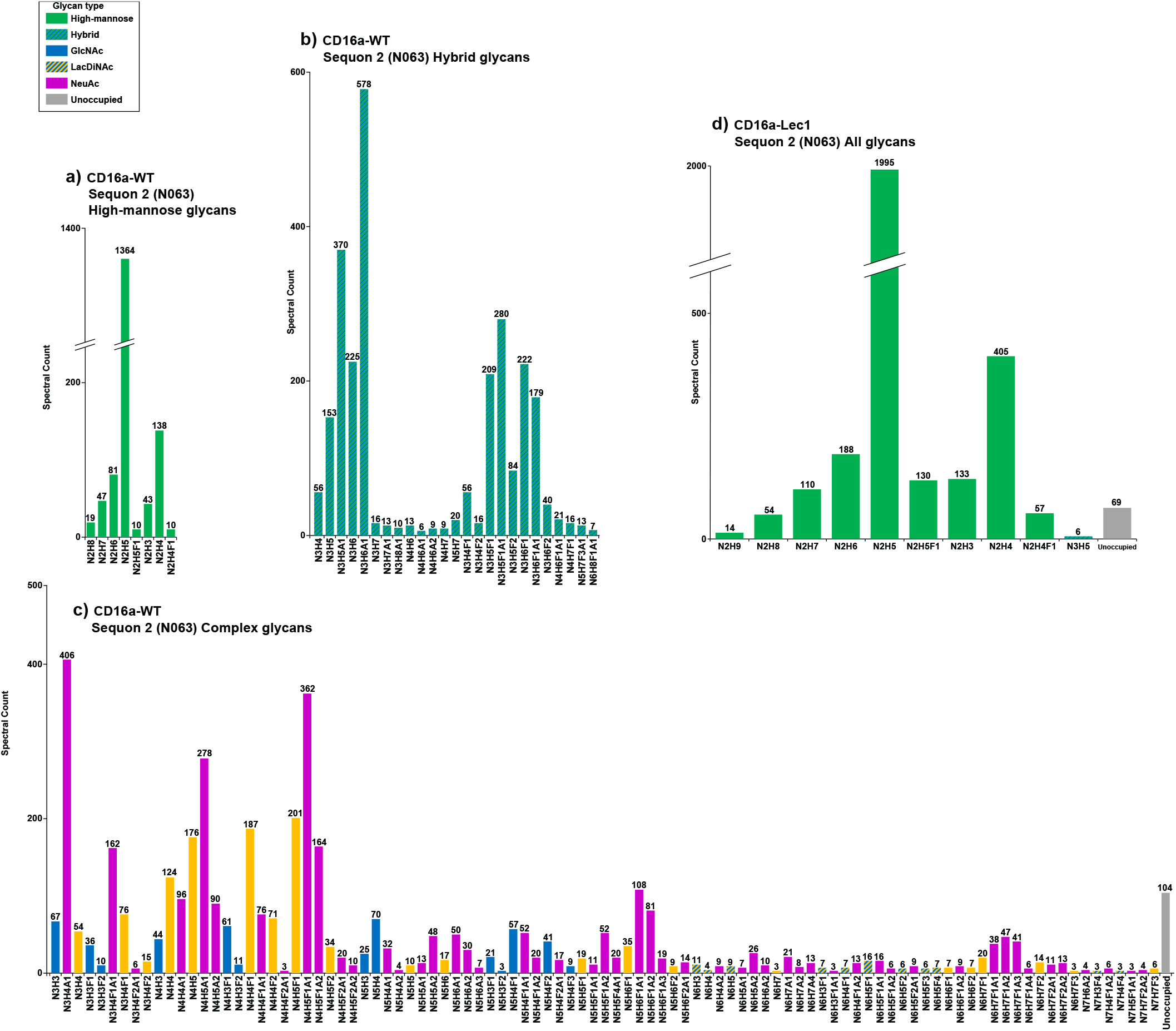
Microheterogeneity at a single site on CD16a (Site 2, N063). Glycopeptide analysis with individual glycan types quantified via spectral count when expressed in a wild-type HEK293F cells or Lec1-HEK293F (MGAT1-) cells, divided by glycan class. *a*, WT-expressed CD16a high-mannose N-glycans. *b*, WT-expressed CD16a hybrid N-glycans. *c*, WT-expressed CD16a complex N-glycans. *d*, Lec1-expressed CD16a N-glycans (all). Colored bars denote glycan terminal features.

All classes of glycans were observed at most sites (**Fig. 3**, S5-S42), with SARS-CoV-2 spike glycoprotein pictured separately due to its large number of sites (**Fig**. S2a). Of particular interest are the sites on reporter glycoproteins that greatly differ from other sites on the same protein: sequons 2 and 4 on CD16a (**Fig. 3a**) and sequon 4 on PDI (**Fig. 3b**) are predominantly less-processed high-mannose and hybrid structures, while the other sites of N-glycosylation on the same proteins are mostly highly processed complex structures. This is in contrast to etanercept (**Fig. 3c**) and erythropoietin (**Fig. 3d**), which have more homogenous N-glycan presentations. The SARS-CoV-2 spike glycoprotein had a diversity of N-glycan presentations on its 22 sites, with most sites enriched with complex N-glycans and certain sites mostly presenting high-mannose N-glycans (**Fig**. S2). Interestingly, we noted that Man5GlcNAc2 was always the most abundant high mannose structure on 36 of our 38 sites with two exceptions being N0234 and N0717 of SARS-CoV-2 that both contain less than 15% complex structures (**Figs**. S2, S27, S35).

**Figure 3:**
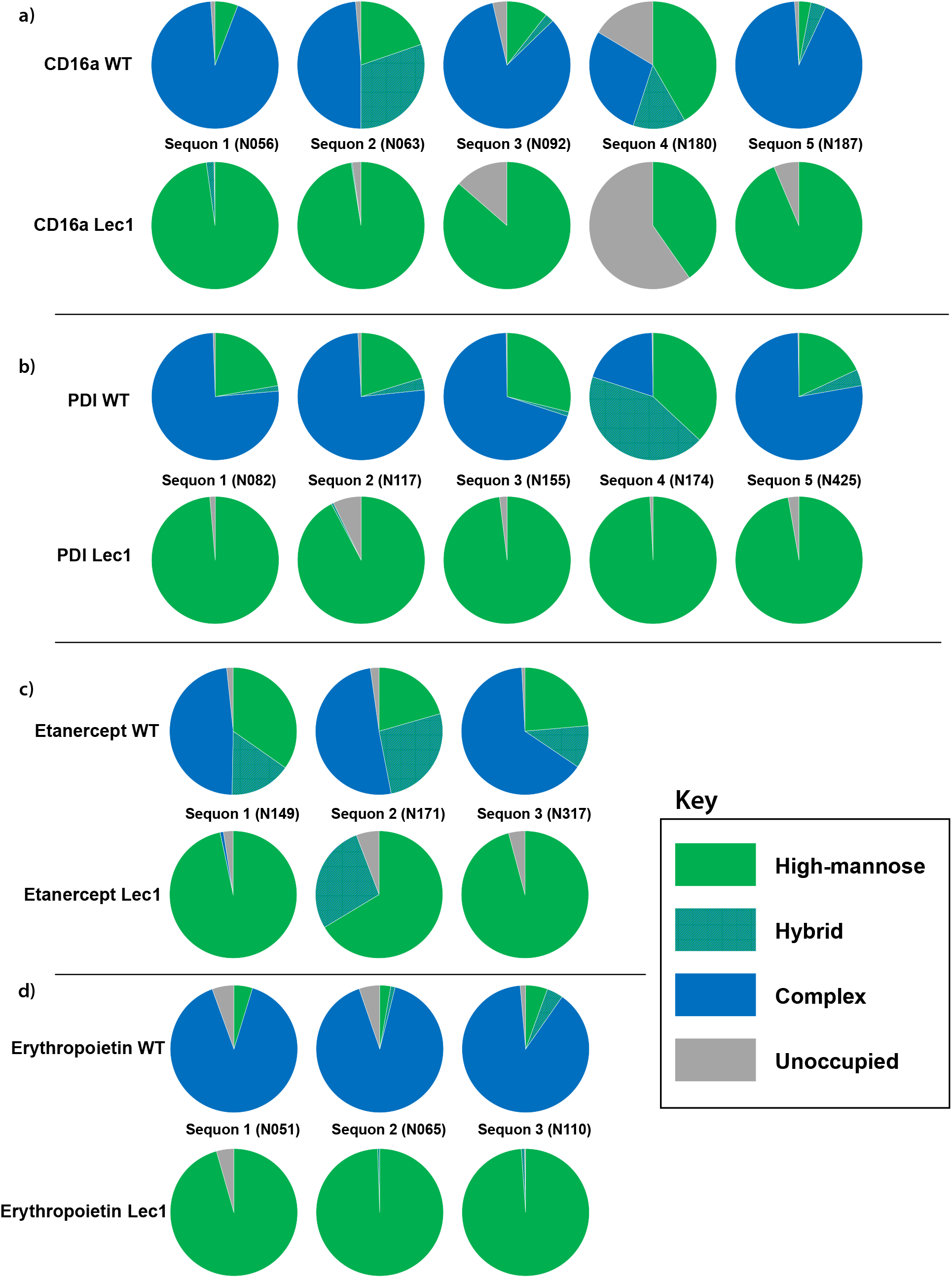
Site occupancy of reporter proteins expressed in WT-HEK293F and Lec1-HEK293F cells. Relative proportion of glycan classes at each site on reporter proteins when expressed in a wild-type or Lec1 (*MGAT1-*) background. *a*, CD16a. *b*, PDI. *c*, Etanercept. *d*, Erythropoietin. Relative populations were ascertained with glycopeptide analysis and quantified with spectral counts

### Expression of reporter proteins in Lec1-HEK293F cells

Next, these same proteins were transiently overexpressed in HEK293S GnTI- (MGAT1 null) cells. The activity of MGAT1 is necessary for the formation of both hybrid and complex N-glycans, as the addition of GlcNAc to the non-reducing end α3-linked mannose is needed for further elaboration and capping by downstream enzymes. Knockout of MGAT1 substantially reduces the diversity of glycans at all sites of N-glycosylation and causes a significant enrichment of Man_5_GlcNAc_2_ structures N-glycans on expressed glycoproteins, as shown on sequon 2 of CD16a (**Fig. 2d**) as well as the other reporter sites (**Figs**. S5-S42). This is useful because it allows for the *in vitro* processing of all N-glycans on a glycoprotein to begin from a common substrate. This enrichment was successful for most sites on all reporter proteins (**Figs. 3**, S2b). However, there is an exception at Sequon 2 of etanercept, which contained a significant population of apparent hybrid N-glycans by an unknown processing event that we are currently exploring.

### Conversion of high-mannose glycans to hybrid glycans

In order to probe the effects of tertiary structure on N-glycan processing, we first monitored the conversion of Man_5_GlcNAc_2_ N-glycans to GlcNAcMan_5_GlcNAc_2_ N-glycans via the addition of GlcNAc by the glycosyltransferase MGAT1 on intact reporter proteins expressed and purified from MGAT1 deficient cells. We did this through a series of time-course reactions using purified protein and MGAT1 in the presence of the nucleotide sugar donor UDP-GlcNAc followed by analysis and quantitation via LC-MS/MS (**Fig. 4**). The ratio of enzymes to molarity of reporter N-glycan sites was kept constant so that we could compare the inter-protein as well as intra-protein rates of N-glycan processing. We observed site-specific rates of GlcNAc addition across the range of our 38 sites of N-glycosylation. Generally, these rates corresponded well to the distributions of N-glycans that were found on the respective sites when the reporters were expressed in wild type HEK293F cells (**Figs. 3**, S5-42). For example, sequons 2 and 4 on CD16a are both modified relatively slowly by MGAT1 (**Fig. 4a**) and are also enriched with high-mannose structures when expressed in wild type HEK293 cells (**Fig. 3a**). This also holds for sequon 4 of PDI (**Fig. 3b**), which is the only site on PDI that is modified slowly by MGAT1 (**Fig. 4b**). This pattern is also demonstrated on SARS-CoV-2 spike glycoprotein: we observed a broad distribution of MGAT1 activity rates across its 22 sites of N-glycosylation (**Figs. 4e**, S3), and the fastest and slowest sites are enriched in complex and high-mannose N-glycans, respectively (**Fig. 4f**). This is best represented by sites N0234 and N0717 on Spike glycoprotein, which had slow transfer rates and primarily were occupied by high-mannose N-glycans when expressed in a wild type background (**Figs. 4f**, S2).

**Figure 4:**
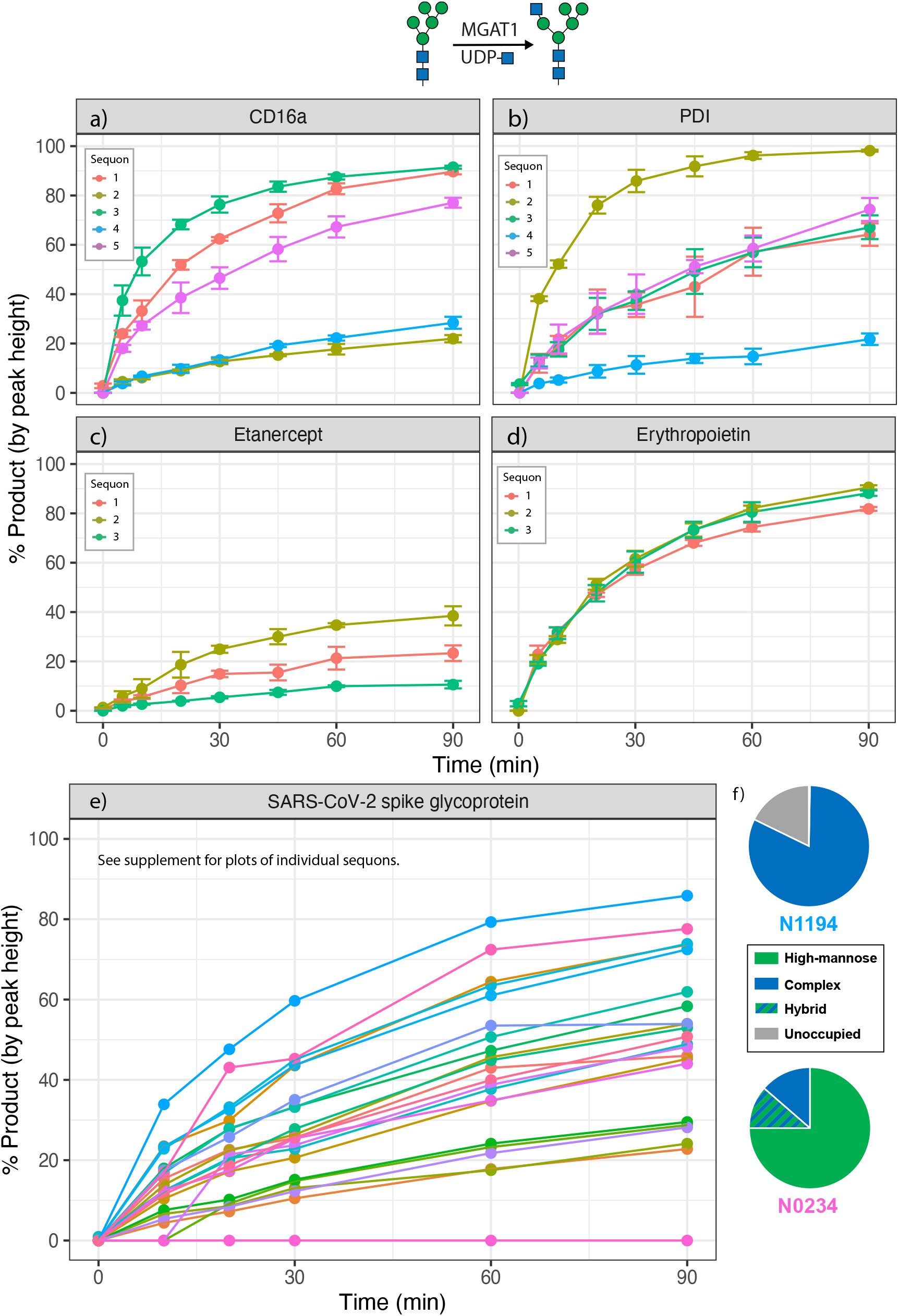
Site-specific monitoring of MGAT1 activity. Time-course reaction of GlcNAc addition to reporter proteins with recombinant MGAT1. *a*, CD16a. *b*, Protein disulfide isomerase. *c*, Etanercept. *d*, Erythropoietin. *e*, SARS-CoV-2 spike glycoprotein. *f*, wild-type glycopeptide profiles for the sites on SARS-CoV-2 spike glycoprotein with the fastest (N1194) and slowest (N0234) rates of MGAT1 activity. Error bars and legend omitted for SARS-CoV-2 spike glycoprotein due to large number of sites. Reaction progress calculated as proportion of the sum of monoisotopic peak heights of product (Man_5_GlcNAc_3_) vs. the sum of product and reactant (Man_5_GlcNAc_2_) peak heights. Experiments performed in triplicate, error bars represent standard error of the mean.

In contrast to the demonstration of site-specificity for PDI and CD16a, all sites on etanercept have relatively high levels of high-mannose glycans (**Fig. 3c**), and all are processed slowly by MGAT1 (**Fig. 4c**). Additionally, erythropoietin’s three sites of N-glycosylation are all processed efficiently by MGAT1 (**Fig. 4d**), and when expressed in wild type HEK293 cells mostly produce complex N-glycans (**Fig. 3d**).

### Conversion of hybrid glycans to complex glycans

The conversion of hybrid N-glycans such as GlcNAcMan_5_GlcNAc_2_ to complex N-glycans requires the activity of two enzymes: the glycoside hydrolase MAN2A1 and the GlcNAc transferase MGAT2. In order to probe the site-specific rates of MAN2A1 activity, we first reacted the reporters expressed in the MGAT1 null cell line with an excess of MGAT1 to enrich GlcNAcMan_5_GlcNAc_2_ structures. After ensuring >80% conversion to product at each site, we then examined conversion of the GlcNAcMan_5_GlcNAc_2_ product to GlcNAcMan_3_GlcNAc_2_ following digestion with MAN2A1 (**Fig. 5**). Similar patterns of site-specific rates were seen as with MGAT1, with sequons 2 and 4 of CD16a (**Fig. 5a**) and sequon 4 of PDI (**Fig. 5b**) all having much lower levels of activity compared to the other sites on the same protein. Again, all sites on etanercept (**Fig. 5c**) were processed much more slowly than those on erythropoietin (**Fig. 5d**). Notably, no cleavage products were observed at sequon 5 (N0149) on SARS-CoV-2 spike glycoprotein (**Fig. S3**). Inspection of the GlcNAcMan_4_GlcNAc_2_ intermediate in MAN2A1 processing revealed that at this site, only one mannose was able to be removed (**Fig. S4**). Despite this *in vitro* observation, when expressed in a wild type background this site produces an abundance of complex-type N-glycans (**Figs**. S2, S25). This is in contrast to sites N0234 and N0717 on the spike glycoprotein, which exhibit both slow transfer rates and an enrichment of high-mannose N-glycans (**Figs**. S2, S3)

**Figure 5:**
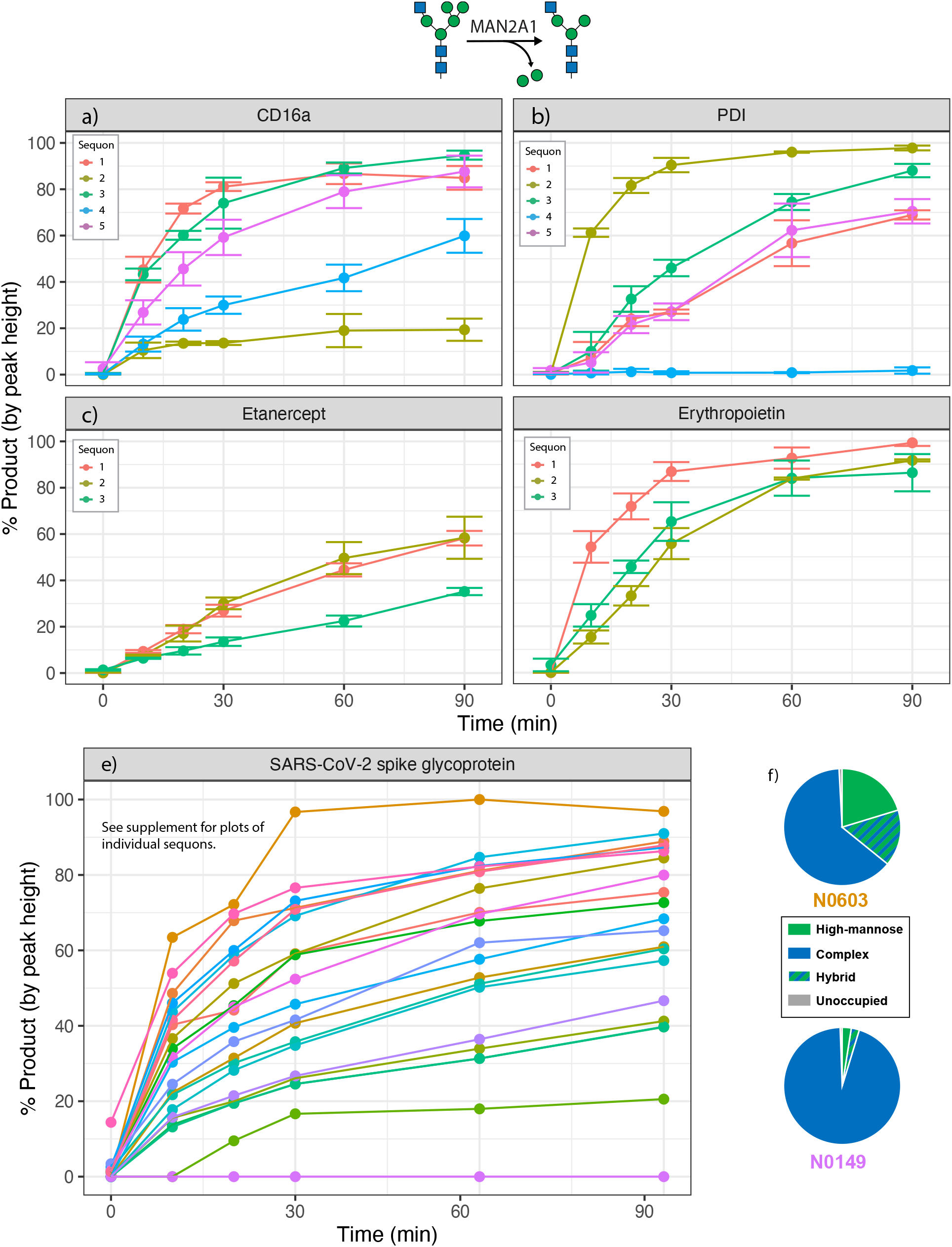
Site-specific monitoring of MAN2A1 activity. Time-course reaction of GlcNAc addition to reporter proteins with recombinant MGAT1. *a*, CD16a. *b*, Protein disulfide isomerase. *c*, Etanercept. *d*, Erythropoietin. *e*, SARS-CoV-2 spike glycoprotein. *f*, wild-type glycopeptide profiles for the sites on SARS-CoV-2 spike glycoprotein with the fastest (N0603) and slowest (N0149) rates of MGAT1 activity. Legend omitted for SARS-CoV-2 spike glycoprotein due to large number of sites. Reaction progress calculated as proportion of the sum of monoisotopic peak heights of product (Man_3_GlcNAc_3_) vs. the sum of product and reactant (Man_5_GlcNAc_3_) peak heights. Experiments performed in triplicate, error bars represent standard error of the mean.

The next step in the formation of complex glycans is the addition of a β-2-linked GlcNAc to the α6-mannose of the GlcNAcMan_3_GlcNAc_2_ moiety. We reacted the reporter proteins expressed in the MGAT1 null cells with an excess of MGAT1, MAN2A1, and the UDP-GlcNAc donor in order to enrich the GlcNAcMan_3_GlcNAc_2_ substrate, then performed another set of time-course reactions with MGAT2. Sites which could not be efficiently converted to GlcNAcMan_3_GlcNAc_2_ structures by MGAT1 and MAN2A1 treatment (e.g. sequon 4 on PDI (**Fig. 5b**)) were excluded from further analyses. Similar patterns of modification were observed in the MGAT2 reactions as were seen in the MGAT1 and MAN2A1 experiments, with lower levels of activity observed at sequons 2 and 4 on CD16a compared to other sites on the same protein (**Fig. 6a**). Additionally, all sites on etanercept (**Fig. 6c**) were processed more slowly than those on erythropoietin (**Fig. 6d**). A broad range of processing rates was observed on SARS-CoV-2 spike glycoprotein (**Fig. 6e**). Sites that were modified fastest in our *in vitro* modification studies were also enriched in complex N-glycans when expressed in wild type HEK293 cells and the sites that were the slowest for *in vitro* modification corresponded to sites that were relatively enriched with high-mannose N-glycans when generated in wild type cells (**Figs. 6f**, S3).

**Figure 6:**
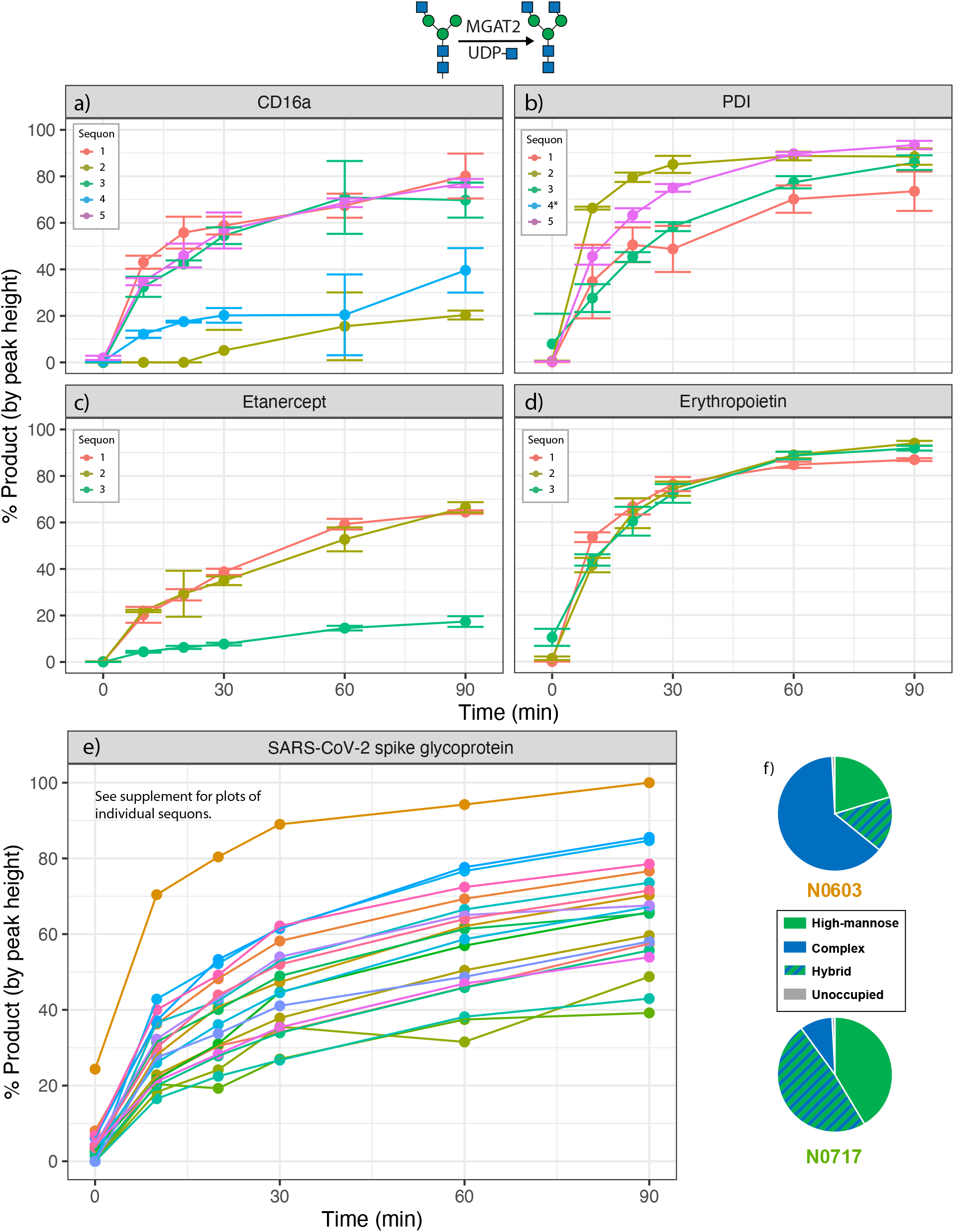
Site-specific monitoring of MGAT2 activity. Time-course reaction of GlcNAc addition to reporter proteins with recombinant MGAT1. *a*, CD16a. *b*, Protein disulfide isomerase. *c*, Etanercept. *d*, Erythropoietin. *e*, SARS-CoV-2 spike glycoprotein. f, wild-type glycopeptide profiles for the sites on SARS-CoV-2 spike glycoprotein with the fastest (N0603) and slowest (N0717) rates of MGAT1 activity. Error bars and legend omitted for SARS-CoV-2 spike glycoprotein due to large number of sites. Asterisks on site legend indicate that not enough substrate was generated from previous N-glycan processing steps to monitor reaction progress. Reaction progress calculated as proportion of the sum of monoisotopic peak heights of product (Man_3_GlcNAc_4_) vs. the sum of product and reactant (Man_3_GlcNAc_3_) peak heights. Experiments performed in triplicate, error bars represent standard error of the mean.

### Core fucosylation of N-glycans by FUT8

Following the above experiments, we wanted to see if similar patterns of site-specific N-glycan processing rates would apply to core fucosylation. Core fucosylation is the attachment of an α1,6-linked fucose to the GlcNAc that is directly attached to the asparagine at the core of N-linked glycans, a reaction which is catalyzed by the fucosyltransferase FUT8. This reaction is generally specific to complex N-glycans (39), and thus we generated GlcNAc_2_Man_3_GlcNAc_2_ glycans on our collection of reporter proteins by reacting with an excess of MGAT1, MAN2A1, and MGAT2 in the presence of the UDP-GlcNAc sugar donor. We then examined the rates of modification of the respective glycans with FUT8 (**Fig. 7**). Interestingly, at many sites we found substantial core fucosylation prior to *in vitro* processing despite the reporter proteins being expressed in an MGAT1-null cell line and thus lacking complex N-glycans (**Figs. 7a-b, d-e**, S5-42). Otherwise, we observed a diversity of fucosylation rates among the different sites. Sequon 2 on CD16a (**Fig. 7a**) and sequon 1 and 5 on PDI were markedly slow (**Fig. 7b**), as well as sequon 3 on etanercept (**Fig. 7c**). All sites on erythropoietin were fucosylated rapidly, which possibly reflects their high initial levels of fucosylation even before the FUT8 reaction (**Fig. 7d**). Most sites on SARS-CoV-2 spike glycoprotein were efficiently fucosylated (**Fig. 7e**), with a few exceptions (N0122, N0801, N1074 N1098) (**Fig. S3**).

**Figure 7:**
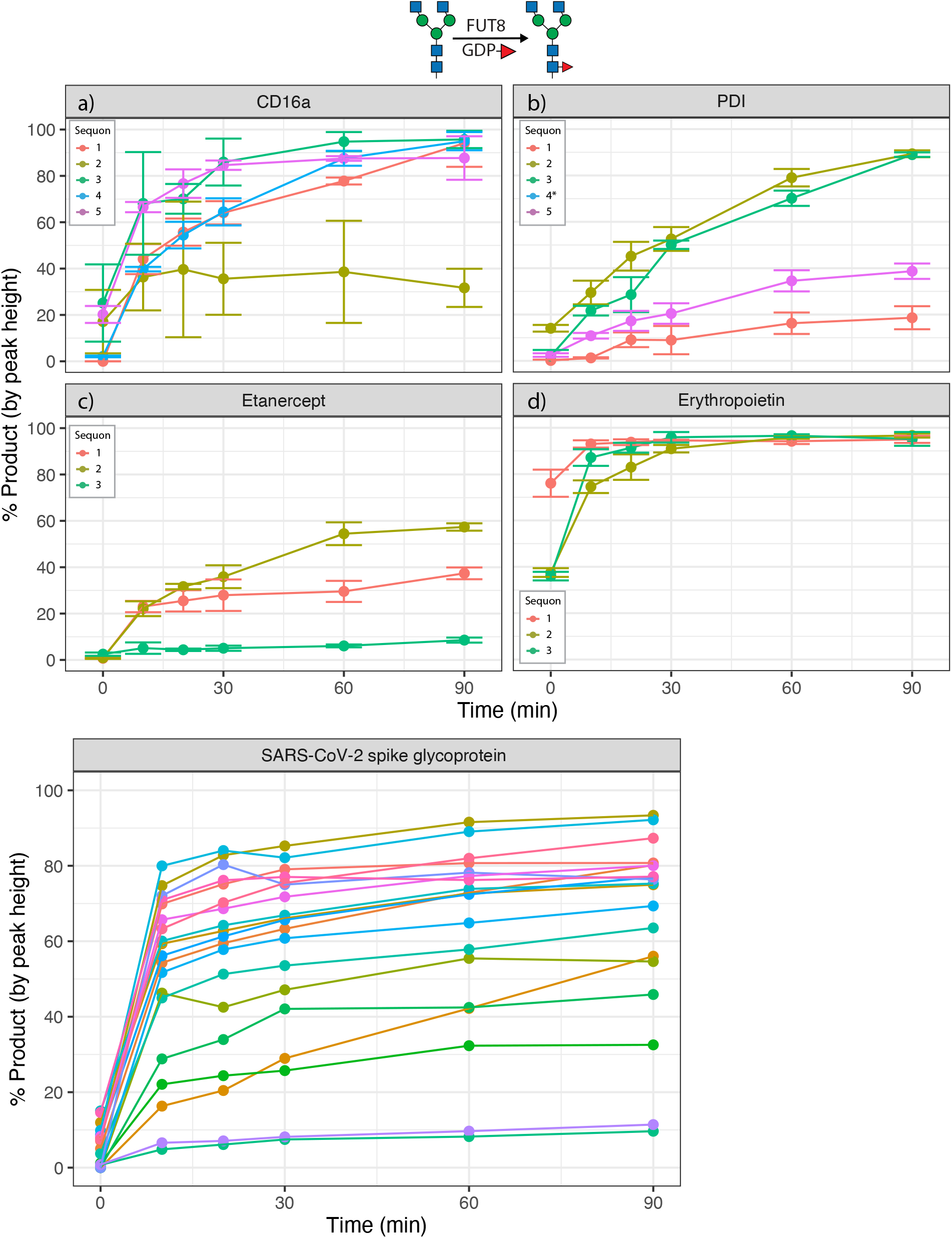
Site-specific monitoring of FUT8 activity. Time-course reaction of Fucose addition to reporter proteins with recombinant FUT8. *a*, CD16a. *b*, Protein disulfide isomerase. *c*, Etanercept. *d*, Erythropoietin. *e*, SARS-CoV-2 spike glycoprotein. Error bars and legend omitted for SARS-CoV-2 spike glycoprotein due to large number of sites. Asterisks on site legend indicate that not enough substrate was generated from previous N-glycan processing steps to monitor reaction progress. Reaction progress calculated as proportion of the sum of monoisotopic peak heights of product (Man_3_GlcNAc_4_) vs. the sum of product and reactant (Man_3_GlcNAc_4_Fuc_1_) peak heights. Experiments performed in triplicate, error bars represent standard error of the mean.

### Impact of tertiary structure on site-specificity

In order to determine whether these site-specific differences in glycosyltransferase rates were due to tertiary structure, we repeated the transfer of GlcNAc onto PDI-Man_5_GlcNAc_2_ glycans using MGAT1, but first digested the protein with trypsin to reduce the fully-folded protein substrate into glycopeptides. Without the reporter protein tertiary structure, all site-specificity of GlcNAc transfer rate was lost (**Fig. 8a** compared to **Fig. 3B**). Since FUT8 activity requires access to the core GlcNAc linked to the Asn residue of the peptide backbone, we were curious to see if cleavage to glycopeptides would eliminate the site-specificity observed on intact protein. Similar to MGAT1, all site-specific modification by FUT8 was lost following cleavage of CD16a to glycopeptides, with all sites (including the slow site 2 (Fig. 7a)) exhibiting similar rates of modification (**Fig. 8b**).

**Figure 8:**
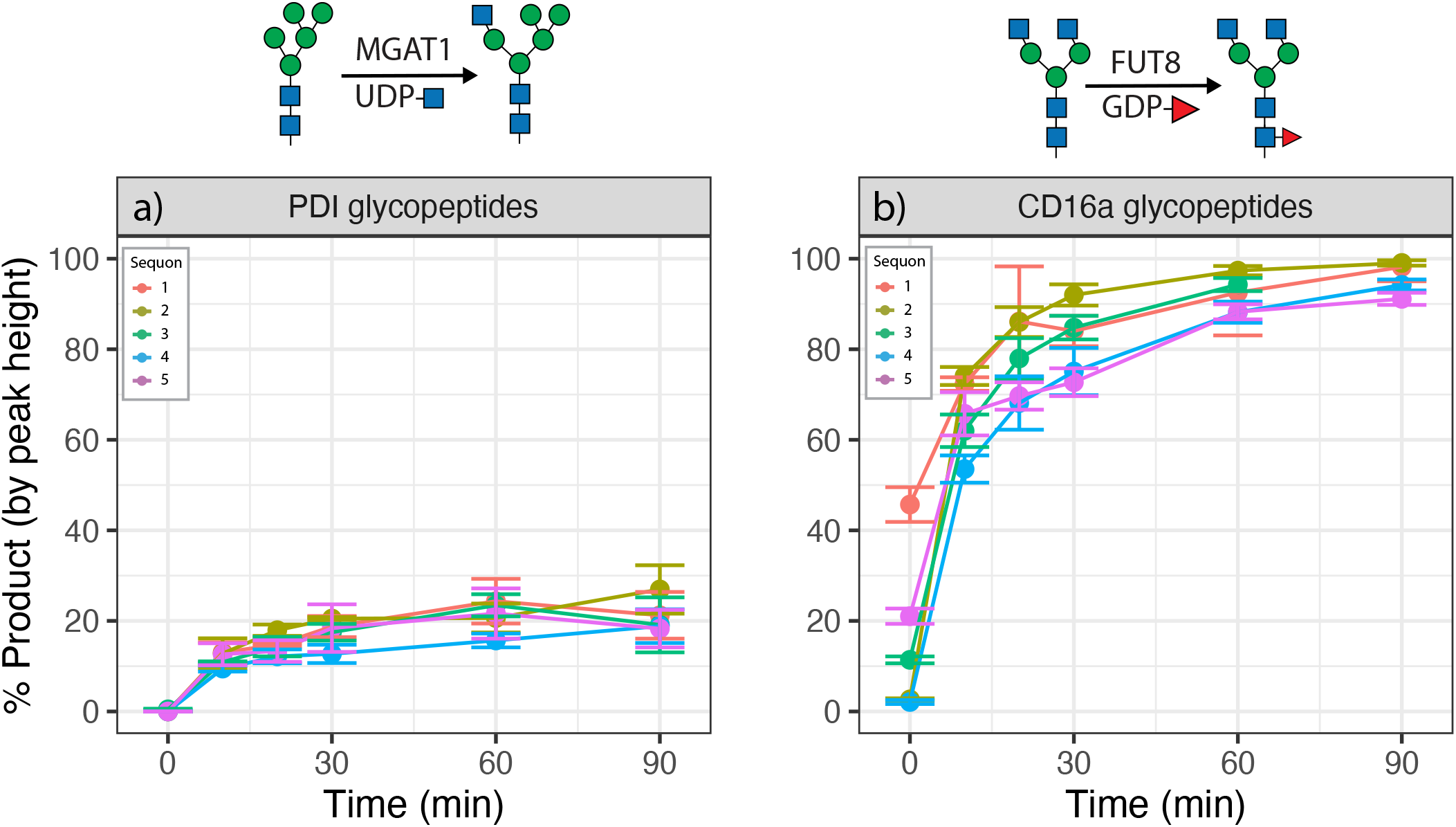
Tertiary structure is responsible for site-specificity of N-glycan processing rates. Time-course reaction of N-glycan processing on proteins digested into glycopeptides with proteases. *a*, Reaction of MGAT1 with PDI glycopeptides. *b*, Reaction of FUT8 with CD16a glycopeptides. Reaction progress calculated as proportion of the sum of monoisotopic peak heights of product vs. the sum of product and reactant peak heights. Experiments performed in triplicate, error bars represent standard error of the mean.

## Discussion

While N-glycans are a crucial component in the production of membrane bound and secreted glycoproteins, the determinants that define the diversity of N-glycan structures at any given site are not well understood. Factors that have been suggested to influence N-glycan microheterogeneity include the availability of glycosyltransferases and glycoside hydrolases (40, 41), the availability of nucleotide sugar donors (42), secretory pathway trafficking (10, 30, 41), and the accessibility of the acceptor site (9, 32, 43). The impact of enzyme availability has mostly been probed through genetic engineering approaches (40). However, availability of enzymes and sugar nucleotides cannot sufficiently explain site specific differences on the same polypeptide nor can protein trafficking in the secretory pathway. Thus, we hypothesize that it is the impact of acceptor site accessibility that allows for site specific differences on the same protein. This could occur by multiple mechanisms including the substrate glycan interacting with the substrate protein backbone at a specific site and thus hampering engagement of the glycoenzyme with the substrate glycan. An example of this is site 4 on PDI that was eloquently demonstrated by Aebi and colleagues (9). Another possibility is local secondary and tertiary structure of the substrate protein at individual sites of modification and the glycoenzyme active site resulting in steric or electrostatic clashes that prohibit optimal binding for catalysis. Interestingly, our analysis revealed that proteolytically digesting substrate proteins before transfer reactions abolished site specific rate differences (**Fig. 8**) that strongly support higher-order structure of the substrate playing an important role in transfer rates and in agreement with proposed mechanisms for microheterogeneity.

Molecular dynamics studies and Markov state modeling by Mathew et al. demonstrated that the relative amount of time a glycan spends extended away from the protein and exposed to solvent correlates with site-specificity of glycan-processing rates on the yeast model protein disulfide isomerase (32). Additionally, they monitored *in vitro* N-glycan processing rates with ER mannosidases as well as MGAT1 and MAN2A1, and the results with overlapping enzymes in our present study agree well. Site 4 of PDI was identified as a “slow” site (**Figs. 4b, 5b**), and their Michaelis-Menten analysis of PDI processing kinetics are also consistent with our observations (**Figs. 4b**, 5b). Additionally, their studies on the kinetics of earlier glycan-processing steps involving ER mannosidase I and Golgi mannosidase IB show that this site-specificity is conserved in earlier steps of N-glycan processing. However, their approach to reduce tertiary structure through reduction and alkylation prior to *in vitro* modification led to differing results compared to our glycopeptide experiments (**Fig. 8)**, and they observed differences in modification rate at different sites. This may be due to some secondary structures of the protein not being completely disrupted without the use of protease digestion to cleave the model protein utilized, or perhaps the denatured protein can still influence site kinetics.

Generally, these results indicate that the tertiary structure specific to an acceptor site can be an important factor in defining the types of N-glycans seen at a given site. In particular, the efficiency (or lack thereof) of MGAT1 and MAN2A1 appears to be highly predictive of high-mannose-type glycans at a sequon. *In vivo* it is likely that MGAT1 is rate-limiting as the most common high-mannose structure at most sites when expressed in a wild type background is the MGAT1 substrate Man_5_GlcNAc_2_ (**Figs**. S5-42), and its activity is required for downstream processing by enzymes like MAN2A1. The rate limiting role of MAN2A1 was also observed in our *in vitro* studies, particularly at sites that were also poorly modified with MGAT1. This may be partly due to lower activity of the recombinant enzyme employed in our *in vitro* studies; four times as much MAN2A1 had to be used in assays compared to the glycosyltransferases (1:250 enzyme:substrate molar ratio for MAN2A1 vs. 1:1,000 for glycosyltransferases). MAN2A1 processing has previously been identified as a potential bottleneck in N-glycan processing (32). Potential steric barriers to MAN2A1 action are suggested by the structure of MAN2A1:substrate complex that demonstrates a significant portion of the total N-linked glycan must fit into the active site of the enzyme for efficient binding and catalysis (44). If a site is not processed totally by MAN2A1, it may form hybrid structures, but cannot form complex structures due to the necessity of mannose trimming on the α6 branch of the tri-mannosyl N-glycan core. By contrast, the active site structure of MGAT1 involved acceptor recognition has not yet been determined, but likely also presents significant steric barriers for access to some poorly modified sites.

While these studies provide a sound starting point for determining what structural features may be important in determining N-glycan destiny, much work remains. We purposely chose reporter proteins and processing enzymes with experimentally determined structures (**Table 1**), (38, 45– 48). We are currently utilizing MD simulations of glycosylated reporter proteins and site-specific docking of specific glycan modified reporters with glycoenzymes to determine site-specific glycans interacting with the reporter protein as well as clashes between the reporter sites and the glycoenzymes. This will guide future work involving mutagenesis studies to influence the rate at which glycosyltransferases and glycosyl hydrolases are able to modify acceptor sites. There is evidence that this approach can indeed alter the distribution of N-glycans at a specific site, as evidenced through modification of a tyrosine residue proximal to sequon 4 on protein disulfide isomerase (9). Taking a systematic approach that involves site-specific rate monitoring coupled with modeling and mutagenesis should result in common rules that not only will allow prediction of microheterogeneity but will allow us to tune it.

## Experimental procedures

### Expression and purification of glycoprotein reporters and glycosylation enzymes for in vitro modification

Expression constructs encoding the reporter proteins were generated with either NH_2_-terminal fusion tags (CD16a (low affinity immunoglobulin gamma Fc region receptor III-A, FCGR3A), UniProt P08637, residues 19-193; Erythropoietin (EPO), UniProt P01588, residues 28-193; Etanercept (TNF receptor-IgG1 fusion), GenBank AKX26891, residues 1-467) or C-terminal fusion tags (yeast PDI1 (protein disulfide-isomerase), UniProt P17967, residues 1-494; SARS-CoV-2 Spike glycoprotein, UniProt P0DTC2, residues 1-1208). The constructs employing N-terminal fusion sequences employed the pGEn2 expression vector while the PDI1 construct was generated in the PGEc2 vector as previously described (49).For the pGEn2 constructs, the fusion protein coding region was comprised of a 25-amino acid signal sequence, an His_8_ tag, AviTag, the “superfolder” GFP coding region, the 7-amino acid recognition sequence of the tobacco etch virus (TEV) protease followed by the catalytic domain region for reporter proteins (49). Constructs encoding MGAT1, MAN2A1, MGAT2 and FUT8 employed the pGEn2 vector and were expressed and purified as previously described (49). For the PDI1 construct the pGEc2 vector was employed and encoded the segment of *Saccromyces cerevisiae* PDI1 indicated followed by an SGSG tetrapeptide, the 7 amino acid TEV recognition sequence, the “superfolder” GFP coding region, and an His_8_ tag (49). For SARS-CoV-2 Spike, the construct contained an additional COOH-terminal trimerization sequence and His6 tag as previously described (50). The recombinant reporter proteins were expressed as a soluble secreted proteins by transient transfection of suspension culture HEK293F cells (FreeStyle™ 293-F cells, Thermo Fisher Scientific, Waltham MA) for wild type glycosylated structures and in HEK293S (GnTI-) cells (ATCC) to generate Man_5_GlcNAc_2_-Asn glycan structures (49, 51). Cultures were maintained at 0.5–3.0×10^6^ cells/ml in a humidified CO_2_ platform shaker incubator at 37°C with 50% humidity. Transient transfection was performed using expression medium comprised of a 9:1 ratio of Freestyle™293 expression medium (Thermo Fisher Scientific, Waltham MA) and EX-Cell expression medium including Glutmax (Sigma-Aldrich). Transfection was initiated by the addition of plasmid DNA and polyethyleneimine as transfection reagent (linear 25-kDa polyethyleneimine, Polysciences, Inc., Warrington, PA). Twenty-four hours post-transfection the cell cultures were diluted with an equal volume of fresh media supplemented with valproic acid (2.2 mM final concentration) and protein production was continued for an additional 5 days at 37°C^3^. The cell cultures were harvested, clarified by sequential centrifugation at 1200 rpm for 10 minutes and 3500 rpm for 15 minutes at 4°C, and passed through a 0.8 µM filter (Millipore, Billerica, MA). The protein preparation was adjusted to contain 20 mM HEPES, 20 mM imidazole, 300 mM NaCl, pH 7.5, and subjected to Ni-NTA Superflow (Qiagen, Valencia, CA) chromatography using a column preequilibrated with 20 mM HEPES, 300 mM NaCl, 20 mM imidazole, pH 7.5 (Buffer I). Following loading of the sample the column was washed with 3 column volumes of Buffer I followed by 3 column volumes of Buffer I containing 50 mM imidazole, and eluted with Buffer I containing 300 mM imidazole at pH 7.0. The protein was concentrated to approximately 3 mg/ml using an ultrafiltration pressure cell (Millipore, Billerica, MA) with a 10-kDa molecular mass cutoff membrane and buffer exchanged with 20 mM HEPES, 100 mM NaCl, pH 7.0, 0.05% Sodium azide and 10% glycerol.

#### In vitro N-Glycan processing

For the time-course reactions, purified reporter proteins generated in *HEK293S (GnTI-)* cells were used. Reactions were performed at 37°C in 1.5-mL Eppendorf tubes in a reaction volume of 150 μL, with 20 mM HEPES (VWR) pH 7.5 and 300 mM NaCl (Fisher). For glycosyltransferases, the corresponding nucleotide sugar, UDP-GlcNAc (Sigma) for MGAT1 and MGAT2 and GDP-Fucose (CarboSynth) for FUT8, was kept in excess at 1 mM. For MGAT1 and MGAT2 reactions, 1 mM MnCl_2_ (Sigma) was supplemented. The concentration of total N-glycans for each reaction was kept at 5 μM: for example, for a reporter protein with 5 sites of N-glycosylation, the concentration of the protein would be 1.25 μM. For MGAT1, MGAT2, and FUT8 reactions, a 1:1,000 enzyme-to-glycan ratio was used with the concentration of respective enzyme at 5 nM; for MAN2A1 reactions, a 1:250 enzyme-to-glycan ratio was used with the concentration of MAN2A1 at 20nM. Prior to adding enzymes, time-course reaction vessels were equilibrated at 37°C for 15 minutes. At each time point, 20 μL of samples were taken and reactions were deactivated by heating at 95°C for 5 minutes. The samples were then digested by proteases and processed for LC-MS/MS analysis.

#### Enzymatic digestion of PDI1, etanercept, EPO, CD16a, and SARS-CoV-2 spike from wild type and HEK293S (GnTI-) cells

All proteins were reduced by incubating with 10 mM of dithiothreitol (Sigma) at 56 °C and alkylated by 27.5 mM of iodoacetamide (Sigma) at room temperature in dark. For the intact glycopeptide analysis, aliquots of PDI1 proteins were digested respectively using trypsin (Promega), a combination of trypsin and Glu-C (Promega), or a combination of trypsin and AspN (Promega); aliquots of etanercept proteins were digested respectively using trypsin (Promega), or AspN (Promega); aliquots of EPO proteins were digested respectively using a combination of trypsin and Glu-C (Promega), or Glu-C (Promega); aliquots of CD16a proteins were digested respectively using chymotrypsin (Athens Research and Technology), AspN (Promega), or a combination of chymotrypsin (Athens Research and Technology) and Glu-C (Promega); aliquots of S proteins were digested respectively using alpha lytic protease (New England BioLabs), chymotrypsin (Athens Research and Technology), a combination of trypsin and Glu-C (Promega), or a combination of Glu-C and AspN (Promega). For the analysis of deglycosylated glycopeptides, aliquots of PDI1 proteins were digested respectively using trypsin (Promega), or a combination of trypsin and Glu-C (Promega); aliquots of etanercept proteins were digested respectively using trypsin (Promega), or AspN (Promega); aliquots of EPO proteins were digested respectively using a combination of trypsin and Glu-C (Promega), or Glu-C (Promega); aliquots of CD16a proteins were digested respectively using chymotrypsin (Athens Research and Technology), or AspN (Promega); aliquots of S proteins were digested respectively using chymotrypsin (Athens Research and Technology), a combination of trypsin and Glu-C (Promega), or AspN (Promega). Following digestion, the proteins were deglycosylated by Endo-H (Promega) followed by PNGaseF (Promega) treatment in the presence of 18O water (Cambridge Isotope Laboratories).

#### LC-MS/MS analysis of glycopeptides of PDI1, etanercept, EPO, CD16a, and SARS-CoV-2 spike from wild type and HEK293S (GnTI-) cells

The resulting peptides from respective enzymatic digestion of each protein were separated on an Acclaim PepMap RSLC C18 column (75 µm x 15 cm) and eluted into the nano-electrospray ion source of an Orbitrap Fusion™ Lumos™ Tribrid™ or an Orbitrap Eclipse™ Tribrid™ mass spectrometer (Thermo Fisher Scientific) at a flow rate of 200 nL/min. The elution gradient for PDI1, etanercept, EPO, and CD16a proteins consists of 1-40% acetonitrile in 0.1% formic acid over 220 minutes followed by 10 minutes of 80% acetonitrile in 0.1% formic acid. The elution gradient for S protein consists of 1-40% acetonitrile in 0.1% formic acid over 370 minutes followed by 10 minutes of 80% acetonitrile in 0.1% formic acid. The spray voltage was set to 2.2 kV and the temperature of the heated capillary was set to 275 °C. For the intact glycopeptide analysis, full MS scans were acquired from m/z 200 to 2000 at 60k resolution, and MS/MS scans following higher-energy collisional dissociation (HCD) with stepped collision energy (15%, 25%, 35%) were collected in the orbitrap at 15k resolution. For the deglycosylated glycopeptide analysis, full MS scans were acquired from m/z 200 to 2000 at 60k resolution, and MS/MS scans following collision-induced dissociation (CID) at 38% collision energy were collected in the ion trap.

For time-course reactions, a shorter LC gradient was used, and digests of the same reporters were combined prior to analysis for higher throughput. The elution gradient used for PDI, etanercept, EPO, and CD16a proteins was 1-80% acetonitrile in 0.1% formic acid over 60 minutes followed by 5 minutes of 80% acetonitrile in 0.1% formic acid. The peptides were eluted into the source of an Orbitrap Fusion™ Tribrid™ mass spectrometer (Thermo Fisher Scientific). The spray voltage was set to 2.25 kV and the temperature of the heated capillary was set to 280°C. Full MS scans were acquired from m/z 300 to 2000 at 60k resolution, and MS/MS scans following collision-induced dissociation (CID) at 38% collision energy were collected in the ion trap. The elution gradient used for SARS-CoV-2 spike glycoprotein was 1-80% acetonitrile in 0.1% formic acid over 300 minutes followed by 10 minutes of 80% acetonitrile in 0.1% formic acid. The peptides were eluted into the source of an Orbitrap Eclipse™ Tribrid™ mass spectrometer (Thermo Fisher Scientific). The spray voltage was set to 2.25 kV and the temperature of the heated capillary was set to 275°C. Full MS scans were acquired from m/z 300 to 1900 at 60k resolution, and MS/MS scans following collision-induced dissociation (CID) at 38% collision energy were collected in the ion trap.

#### MS data analysis

For the intact glycopeptide analysis, the raw spectra were analyzed using pGlyco3 (52) for database searches with mass tolerance set as 20 ppm for both precursors and fragments. The database search output was filtered to reach a 1% false discovery rate for glycans and 10% for peptides. The filtered result was further validated by manual examination of the raw spectra. For isobaric glycan compositions, fragments in the MS/MS spectra were evaluated to provide the most probable topologies. Quantitation was performed by calculating spectral counts for each glycan composition at each site. Any N-linked glycan compositions identified by only one spectra were removed from quantitation. For the deglycosylated glycopeptide analysis, the spectra were analyzed using SEQUEST (Proteome Discoverer 1.4 and 2.5, Thermo Fisher Scientific) with mass tolerance set as 20 ppm for precursors and 0.5 Da for fragments. The search output from Proteome Discoverer 1.4 was filtered using ProteoIQ (v2.7, Premier Biosoft) to reach a 1% false discovery rate at protein level and 10% at peptide level. The search output from Proteome Discoverer 2.5 was filtered within the program to reach a 1% false discovery rate at protein level and 10% at peptide level. Occupancy of each N-linked glycosylation site was calculated using spectral counts assigned to the 18O-Asp-containing (PNGaseF-cleaved) and/or HexNAc-modified (EndoH-cleaved) peptides and their unmodified counterparts.

For time-course reactions, quantitation was performed through manual inspection of MS1 spectra using Thermo Freestyle 1.7 (Thermo Fischer Scientific). The intensities of monoisotopic peak heights for all observable charge states for reactants and products were determined and then summed and averaged in triplicate to determine percent conversion to product over time. Plots generated using RStudio (1.4.1717).

## Data availability

All data generated or analyzed during this study are included in this article and supporting information files. The glycopeptide analysis MS data have been deposited to the ProteomeXchange Consortium via the PRIDE partner repository with the dataset identifier PXD032149. MS data for time-course reactions available upon request.

## Supporting information

This article contains supporting figures (**Figs**. S1-S42) and tables (Tables S1-S3).

## Acknowledgements

We thank Dr. Henrik Clausen for providing cell lines for this project.

## Funding and additional information

This work was supported in part by National Institutes of Health Grant 5R01GM130915 from NIGMS (to K.W.M. and L.W.) and the Glycoscience Training Program (5T32GM107004) from NIGMS. The content is solely the responsibility of the authors and does not necessarily represent the official views of the National Institutes of Health.

## Abbreviations and nomenclature

ER: endoplasmic reticulum
GlcNAc: N-acetylglucosamine
MS: mass spectrometry
MS/MS: tandem mass spectrometry
LC: liquid chromatography

## References

1. Varki, A. (2017) Biological roles of glycans. Glycobiology. 27, 3–49

2. Moremen, K. W., Tiemeyer, M., and Nairn, A. V. (2012) Vertebrate protein glycosylation: diversity, synthesis and function. Nat. Rev. Mol. Cell Biol. 13, 448–462

3. Stanley, P., Taniguchi, N., and Aebi, M. (2015) N-Glycans. in Essentials of Glycobiology, 3rd Edition, p. Chapter 9, Cold Spring Harbor Laboratory Press, Cold Spring Harbor (NY)

4. Solá, R. J., and Griebenow, K. (2009) Effects of glycosylation on the stability of protein pharmaceuticals. J. Pharm. Sci. 98, 1223–1245

5. Wild, R., Kowal, J., Eyring, J., Ngwa, E. M., Aebi, M., and Locher, K. P. (2018) Structure of the yeast oligosaccharyltransferase complex gives insight into eukaryotic N-glycosylation. Science. 359, 545–550

6. Kasturi, L., Eshleman, J. R., Wunner, W. H., and Shakin-Eshleman, S. H. (1995) The hydroxy amino acid in an Asn-X-Ser/Thr sequon can influence N-linked core glycosylation efficiency and the level of expression of a cell surface glycoprotein. J. Biol. Chem. 270, 14756–14761

7. Hart, G. W., Brew, K., Grant, G. A., Bradshaw, R. A., and Lennarz, W. J. (1979) Primary structural requirements for the enzymatic formation of the N-glycosidic bond in glycoproteins. Studies with natural and synthetic peptides. J. Biol. Chem. 254, 9747–9753

8. Stanley, P., Moremen, K. W., Lewis, N. E., Taniguchi, N., and Aebi, M. (2022) N-Glycans. in Essentials of Glycobiology, 4th Ed. (Varki, A., Cummings, R. D., Esko, J. D., Stanley, P., Hart, G. W., Aebi, M., Mohnen, D., Kinoshita, T., Packer, N. H., Prestegard, J. H., Schnaar, R. L., and Seeberger, P. H. eds), Cold Spring Harbor Laboratory Press, Cold Spring Harbor (NY), [online] http://www.ncbi.nlm.nih.gov/books/NBK579964/ (Accessed July 18, 2022)

9. Losfeld, M.-E., Scibona, E., Lin, C.-W., Villiger, T. K., Gauss, R., Morbidelli, M., and Aebi, M. (2017) Influence of protein/glycan interaction on site-specific glycan heterogeneity. FASEB J. 31, 4623–4635

10. Arigoni-Affolter, I., Scibona, E., Lin, C.-W., Brühlmann, D., Souquet, J., Broly, H., and Aebi, M. (2019) Mechanistic reconstruction of glycoprotein secretion through monitoring of intracellular N-glycan processing. Sci. Adv. 5, eaax8930

11. Harpaz, N., and Schachter, H. (1980) Control of glycoprotein synthesis. Processing of asparagine-linked oligosaccharides by one or more rat liver Golgi alpha-D-mannosidases dependent on the prior action of UDP-N-acetylglucosamine: alpha-D-mannoside beta 2-N-acetylglucosaminyltransferase I. J. Biol. Chem. 255, 4894–4902

12. Narasimhan, S., Stanley, P., and Schachter, H. (1977) Control of glycoprotein synthesis. Lectin-resistant mutant containing only one of two distinct N-acetylglucosaminyltransferase activities present in wild type Chinese hamster ovary cells. J. Biol. Chem. 252, 3926–3933

13. Harpaz, N., and Schachter, H. (1980) Control of glycoprotein synthesis. Bovine colostrum UDP-N-acetylglucosamine:alpha-D-mannoside beta 2-N-acetylglucosaminyltransferase I. Separation from UDP-N-acetylglucosamine:alpha-D-mannoside beta 2-N-acetylglucosaminyltransferase II, partial purification, and substrate specificity. J. Biol. Chem. 255, 4885–4893

14. Gordon, R. D., Sivarajah, P., Satkunarajah, M., Ma, D., Tarling, C. A., Vizitiu, D., Withers,S. G., and Rini, J. M. (2006) X-ray Crystal Structures of Rabbit N-acetylglucosaminyltransferase I (GnT I) in Complex with Donor Substrate Analogues. J. Mol. Biol. 360, 67–79

15. Cymer, F., Beck, H., Rohde, A., and Reusch, D. (2018) Therapeutic monoclonal antibody N-glycosylation –Structure, function and therapeutic potential. Biologicals. 52, 1–11

16. Subedi, G. P., and Barb, A. W. (2015) The Structural Role of Antibody N-Glycosylation in Receptor Interactions. Structure. 23, 1573–1583

17. Li, T., DiLillo, D. J., Bournazos, S., Giddens, J. P., Ravetch, J. V., and Wang, L.-X. (2017) Modulating IgG effector function by Fc glycan engineering. Proc. Natl. Acad. Sci. 114, 3485–3490

18. Hayes, J. M., Frostell, A., Karlsson, R., Müller, S., Martín, S. M., Pauers, M., Reuss, F., Cosgrave, E. F., Anneren, C., Davey, G. P., and Rudd, P. M. (2017) Identification of Fc Gamma Receptor Glycoforms That Produce Differential Binding Kinetics for Rituximab. Mol. Cell. Proteomics. 16, 1770–1788

19. Patel, K. R., Roberts, J. T., Subedi, G. P., and Barb, A. W. (2018) Restricted processing of CD16a/Fc γ receptor IIIa N-glycans from primary human NK cells impacts structure and function. J. Biol. Chem. 293, 3477–3489

20. Higel, F., Seidl, A., Sörgel, F., and Friess, W. (2016) N-glycosylation heterogeneity and the influence on structure, function and pharmacokinetics of monoclonal antibodies and Fc fusion proteins. Eur. J. Pharm. Biopharm. 100, 94–100

21. Azevedo, V., Hassett, B., Fonseca, J. E., Atsumi, T., Coindreau, J., Jacobs, I., Mahgoub, E., O’Brien, J., Singh, E., Vicik, S., and Fitzpatrick, B. (2016) Differentiating biosimilarity and comparability in biotherapeutics. Clin. Rheumatol. 35, 2877–2886

22. Zhou, T., Doria-Rose, N. A., Cheng, C., Stewart-Jones, G. B. E., Chuang, G.-Y., Chambers, M., Druz, A., Geng, H., McKee, K., Kwon, Y. D., O’Dell, S., Sastry, M., Schmidt, S. D., Xu, K., Chen, L., Chen, R. E., Louder, M. K., Pancera, M., Wanninger, T. G., Zhang, B., Zheng, A., Farney, S. K., Foulds, K. E., Georgiev, I. S., Joyce, M. G., Lemmin, T., Narpala, S., Rawi, R., Soto, C., Todd, J.-P., Shen, C.-H., Tsybovsky, Y., Yang, Y., Zhao, P., Haynes, B. F., Stamatatos, L., Tiemeyer, M., Wells, L., Scorpio, D. G., Shapiro, L., McDermott, A. B., Mascola, J. R., and Kwong, P. D. (2017) Quantification of the Impact of the HIV-1-Glycan Shield on Antibody Elicitation. Cell Rep. 19, 719–732

23. Broszeit, F., van Beek, R. J., Unione, L., Bestebroer, T. M., Chapla, D., Yang, J.-Y., Moremen, K. W., Herfst, S., Fouchier, R. A. M., de Vries, R. P., and Boons, G.-J. (2021) Glycan remodeled erythrocytes facilitate antigenic characterization of recent A/H3N2 influenza viruses. Nat. Commun. 12, 5449

24. Zhao, P., Praissman, J. L., Grant, O. C., Cai, Y., Xiao, T., Rosenbalm, K. E., Aoki, K., Kellman, B. P., Bridger, R., Barouch, D. H., Brindley, M. A., Lewis, N. E., Tiemeyer, M., Chen, B., Woods, R. J., and Wells, L. (2020) Virus-Receptor Interactions of Glycosylated SARS-CoV-2 Spike and Human ACE2 Receptor. Cell Host Microbe. 28, 586-601.e6

25. Swiedler, S. J., Freed, J. H., Tarentino, A. L., Plummer, T. H., and Hart, G. W. (1985) Oligosaccharide microheterogeneity of the murine major histocompatibility antigens. Reproducible site-specific patterns of sialylation and branching in asparagine-linked oligosaccharides. J. Biol. Chem. 260, 4046–4054

26. Brisson, J. R., and Carver, J. P. (1983) The relation of three-dimensional structure to biosynthesis in the N-linked oligosaccharides. Can. J. Biochem. Cell Biol. Rev. Can. Biochim. Biol. Cell. 61, 1067–1078

27. Savvidou, G., Klein, M., Grey, A. A., Dorrington, K. J., and Carver, J. P. (1984) Possible role for peptide-oligosaccharide interactions in differential oligosaccharide processing at asparagine-107 of the light chain and asparagine-297 of the heavy chain in a monoclonal IgG1 kappa. Biochemistry. 23, 3736–3740

28. Watanabe, Y., Bowden, T. A., Wilson, I. A., and Crispin, M. (2019) Exploitation of glycosylation in enveloped virus pathobiology. Biochim. Biophys. Acta Gen. Subj. 1863, 1480–1497

29. Patel, K. R., Nott, J. D., and Barb, A. W. (2019) Primary Human Natural Killer Cells Retain Proinflammatory IgG1 at the Cell Surface and Express CD16a Glycoforms with Donor-dependent Variability. Mol. Cell. Proteomics. 18, 2178–2190

30. Hang, I., Lin, C., Grant, O. C., Fleurkens, S., Villiger, T. K., Soos, M., Morbidelli, M., Woods, R. J., Gauss, R., and Aebi, M. (2015) Analysis of site-specific N - glycan remodeling in the endoplasmic reticulum and the Golgi. Glycobiology. 25, 1335–1349

31. Weiß, R. G., Losfeld, M.-E., Aebi, M., and Riniker, S. (2021) N-Glycosylation Enhances Conformational Flexibility of Protein Disulfide Isomerase Revealed by Microsecond Molecular Dynamics and Markov State Modeling. J. Phys. Chem. B. 125, 9467–9479

32. Mathew, C., Weiß, R. G., Giese, C., Lin, C., Losfeld, M.-E., Glockshuber, R., Riniker, S., and Aebi, M. (2021) Glycan–protein interactions determine kinetics of N - glycan remodeling. RSC Chem. Biol. 2, 917–931

33. Patel, K. R., Rodriguez Benavente, M. C., Lorenz, W. W., Mace, E. M., and Barb, A. W. (2020) Fc γ receptor IIIa / CD16a processing correlates with the expression of glycan-related genes in human natural killer cells. J. Biol. Chem. 10.1074/jbc.RA120.015516

34. Liu, L., Gomathinayagam, S., Hamuro, L., Prueksaritanont, T., Wang, W., Stadheim, T. A., and Hamilton, S. R. (2013) The Impact of Glycosylation on the Pharmacokinetics of a TNFR2:Fc Fusion Protein Expressed in Glycoengineered Pichia Pastoris. Pharm. Res. 30, 803–812

35. Čaval, T., Tian, W., Yang, Z., Clausen, H., and Heck, A. J. R. (2018) Direct quality control of glycoengineered erythropoietin variants. Nat. Commun. 10.1038/s41467-018-05536-3

36. Sinclair, A. M., and Elliott, S. (2005) Glycoengineering: The effect of glycosylation on the properties of therapeutic proteins. J. Pharm. Sci. 94, 1626–1635

37. Banks, D. D. (2011) The Effect of Glycosylation on the Folding Kinetics of Erythropoietin. J. Mol. Biol. 412, 536–550

38. Sztain, T., Ahn, S.-H., Bogetti, A. T., Casalino, L., Goldsmith, J. A., Seitz, E., McCool, R. S., Kearns, F. L., Acosta-Reyes, F., Maji, S., Mashayekhi, G., McCammon, J. A., Ourmazd, A., Frank, J., McLellan, J. S., Chong, L. T., and Amaro, R. E. (2021) A glycan gate controls opening of the SARS-CoV-2 spike protein. Nat. Chem. 10.1038/s41557-021-00758-3

39. García-García, A., Serna, S., Yang, Z., Delso, I., Taleb, V., Hicks, T., Artschwager, R., Vakhrushev, S. Y., Clausen, H., Angulo, J., Corzana, F., Reichardt, N. C., and Hurtado-Guerrero, R. (2021) FUT8-Directed Core Fucosylation of N-glycans Is Regulated by the Glycan Structure and Protein Environment. ACS Catal. 10.1021/acscatal.1c01698

40. Narimatsu, Y., Büll, C., Chen, Y.-H., Wandall, H. H., Yang, Z., and Clausen, H. (2021) Genetic glycoengineering in mammalian cells. J. Biol. Chem. 296, 100448

41. Hirschberg, K., and Lippincott-Schwartz, J. (1999) Secretory pathway kinetics and in vivo analysis of protein traffic from the Golgi complex to the cell surface. FASEB J. Off. Publ. Fed. Am. Soc. Exp. Biol. 13 Suppl 2, S251–256

42. Burleigh, S. C., van de Laar, T., Stroop, C. J. M., van Grunsven, W. M. J., O’Donoghue, N., Rudd, P. M., and Davey, G. P. (2011) Synergizing metabolic flux analysis and nucleotide sugar metabolism to understand the control of glycosylation of recombinant protein in CHO cells. BMC Biotechnol. 11, 95

43. Yu, X., Baruah, K., Harvey, D. J., Vasiljevic, S., Alonzi, D. S., Song, B.-D., Higgins, M. K., Bowden, T. A., Scanlan, C. N., and Crispin, M. (2013) Engineering hydrophobic protein-carbohydrate interactions to fine-tune monoclonal antibodies. J. Am. Chem. Soc. 135, 9723–9732

44. Rose, D. R. (2012) Structure, mechanism and inhibition of Golgi α-mannosidase II. Curr. Opin. Struct. Biol. 22, 558–562

45. Mukai, Y., Nakamura, T., Yoshikawa, M., Yoshioka, Y., Tsunoda, S., Nakagawa, S., Yamagata, Y., and Tsutsumi, Y. (2010) Solution of the Structure of the TNF-TNFR2 Complex. Sci. Signal. 10.1126/scisignal.2000954

46. Syed, R. S., Reid, S. W., Li, C., Cheetham, J. C., Aoki, K. H., Liu, B., Zhan, H., Osslund, T. D., Chirino, A. J., Zhang, J., Finer-Moore, J., Elliott, S., Sitney, K., Katz, B. A., Matthews, D. J., Wendoloski, J. J., Egrie, J., and Stroud, R. M. (1998) Efficiency of signalling through cytokine receptors depends critically on receptor orientation. Nature. 395, 511–516

47. Isoda, Y., Yagi, H., Satoh, T., Shibata-Koyama, M., Masuda, K., Satoh, M., Kato, K., and Iida, S. (2015) Importance of the Side Chain at Position 296 of Antibody Fc in Interactions with FcγRIIIa and Other Fcγ Receptors. PLOS ONE. 10, e0140120

48. Tian, G., Xiang, S., Noiva, R., Lennarz, W. J., and Schindelin, H. (2006) The Crystal Structure of Yeast Protein Disulfide Isomerase Suggests Cooperativity between Its Active Sites. Cell. 124, 61–73

49. Moremen, K. W., Ramiah, A., Stuart, M., Steel, J., Meng, L., Forouhar, F., Moniz, H. A., Gahlay, G., Gao, Z., Chapla, D., Wang, S., Yang, J.-Y., Prabhakar, P. K., Johnson, R., Rosa, M. dela, Geisler, C., Nairn, A. V., Seetharaman, J., Wu, S.-C., Tong, L., Gilbert, H. J., LaBaer, J., and Jarvis, D. L. (2018) Expression system for structural and functional studies of human glycosylation enzymes. Nat. Chem. Biol. 14, 156–162

50. Xiao, T., Lu, J., Zhang, J., Johnson, R. I., McKay, L. G. A., Storm, N., Lavine, C. L., Peng, H., Cai, Y., Rits-Volloch, S., Lu, S., Quinlan, B. D., Farzan, M., Seaman, M. S., Griffiths, A., and Chen, B. (2021) A trimeric human angiotensin-converting enzyme 2 as an anti-SARS-CoV-2 agent. Nat. Struct. Mol. Biol. 28, 202–209

51. Meng, L., Forouhar, F., Thieker, D., Gao, Z., Ramiah, A., Moniz, H., Xiang, Y., Seetharaman, J., Milaninia, S., Su, M., Bridger, R., Veillon, L., Azadi, P., Kornhaber, G., Wells, L., Montelione, G. T., Woods, R. J., Tong, L., and Moremen, K. W. (2013) Enzymatic basis for N-glycan sialylation: structure of rat α2,6-sialyltransferase (ST6GAL1) reveals conserved and unique features for glycan sialylation. J. Biol. Chem. 288, 34680–34698

52. Zeng, W.-F., Cao, W.-Q., Liu, M.-Q., He, S.-M., and Yang, P.-Y. (2021) Precise, fast and comprehensive analysis of intact glycopeptides and modified glycans with pGlyco3. Nat. Methods. 18, 1515–1523

