## Supporting Information for "Sequential *in vitro* enzymatic N-glycoprotein modification reveals site-specific rates of glycoenzyme processing"

**S1. Protein sequences and purification.**
**S2. SARS-CoV-2 spike glycoprotein site occupancy**
**S3. Site-specific N-glycan processing rates of SARS-CoV-2 spike glycoprotein**
**S4. Levels of GlcNAc3Man4 intermediate during MAN2A1 processing**
**S5-9. N-glycan site occupancy on CD16a when expressed in WT-HEK293F and Lec1-HEK293F cells.**
**S10-14. N-glycan site occupancy on PDI when expressed in Lec1-HEK293F cells.**
**S15-17. N-glycan site occupancy on etanercept when expressed in WT-HEK293F and Lec1-HEK293F cells.**
**S18-20. N-glycan site occupancy on erythropoietin when expressed in WT-HEK293F and Lec1-HEK293F cells.**
**S21-42. N-glycan site occupancy on SARS-CoV-2 spike glycoprotein when expressed in WT-HEK293F and Lec1-HEK293F cells.**

**Table S1. Glycan occupancy at each site of protein disulfide-isomerase (PDI1), etanercept, erythropoietin (EPO), low affinity immunoglobulin gamma Fc region receptor III-A (FCGR3A/etanercept), and spike glycoprotein (S) from wild type and Lec1 (GnT1-/-) HEK293 cells.**

**Table S2. Glycan types identified at each site of protein disulfide-isomerase (PDI1), etanercept, erythropoietin (EPO), low affinity immunoglobulin gamma Fc region receptor III-A (FCGR3A/etanercept), and spike glycoprotein (S) from wild type and Lec1 (GnT1-/-) HEK293 cells.**

**Table S3: Glycan topologies identified at each site of protein disulfide-isomerase (PDI1), etanercept, erythropoietin (EPO), low affinity immunoglobulin gamma Fc region receptor III-A (FCGR3A/etanercept), and spike glycoprotein (S) from wild type and Lec1 (GnT1-/-) HEK293 cells.** Asn(N)# indicates the numbers of asparagines in protein sequences. In toplogies: N=HexNAc, H=Hexose (Hex), F=Fucose (Fuc), A=Neu5Ac. In fucosylation: NoFuc=No Fuc identified; 1Core=One Fuc identified at core position; 1Term=One Fuc identified at terminal position; 1Core and 1Term=One Fuc identified as a mixture of core and terminal positions; 1Core1Term=Two Fuc identified and one is at core and the other is at terminal; 2Term=Two Fuc identified at terminal positions; 1Core1Term and 2Term=Two Fuc identified as a mixture of core and terminal positions; 1Core2Term=Three Fuc identified and one is at core and the others are at terminal; 3Term=Three Fuc identified at terminal positions; 1Core2Term and 3Term=Three Fuc identified as a mixture of core and terminal positions; 1Core3Term=Four Fuc identified and one is at core and the others are at terminal; 4Term=Four Fuc identified at terminal positions; 1Core3Term and 4Term=Four Fuc identified as a mixture of core and terminal positions; 1Core4Term=Five Fuc identified and one is at core and the others are at terminal; 5Term=Five Fuc identified at terminal positions; 1Core4Term and 5Term=Five Fuc identified as a mixture of core and terminal positions; 1Core5Term=Six Fuc identified and one is at core and the others are at terminal; 6Term=Six Fuc identified at terminal positions; 1Core5Term and 6Term=Six Fuc identified as a mixture of core and terminal positions.

**S1: Protein sequences and purification.**
**S1a: Reporter protein sequences**

**Erythropoietin**
001 HHHHHHHHMS GLNDIFEAQK IEWHEMSKGE LFTGVVPILV ELDGDVNGHK
051 FSVRGEGEGD ATNGKLTLKF ICTTGKLPVP WPTLVTTLTY GVQCFSRYPD
101 HMKRHDFFKS AMPEGYVQER TISFKDDGTY KTRAEVKFEG DTLVNRIELK
151 GIDFKEDGNI LGHKLEYNFN SHNVYITADK QKNGIKANFK IRHNVEDGSV
201 QLADHYQQNT PIGDGPVLLP DNHYLSTQSV LSKDPNEKRD HMVLLEFVTA
251 AGITHGEFAS TSLYKKAGSE NLYFQGAPPR LICDSRVLER YLLEAKEAEN
301 ITTGCAEHCS LNENITVPDT KVNFYAWKRM EVGQQAVEVW QGLALLSEAV
351 LRGQALLVNS SQPWEPLQLH VDKAVSGLRS LTTLLRALGA QKEAISPPDA
401 ASAAPLRTIT ADTFRKLFRV YSNFLRGKLK LYTGEACRTG DR

**Etanercept**
001 HHHHHHHHMS GLNDIFEAQK IEWHEMSKGE ELFTGVVPIL VELDGDVNGH
051 KFSVRGEGEG DATNGKLTLK FICTTGKLPV PWPTLVTTLT YGVQCFSRYP
101 DHMKRHDFFK SAMPEGYVQE RTISFKDDGT YKTRAEVKFE GDTLVNRIEL
151 KGIDFKEDGN ILGHKLEYNF NSHNVYITAD KQKNGIKANF KIRHNVEDGS
201 VQLADHYQQN TPIGDGPVLL PDNHYLSTQS VLSKDPNEKR DHMVLLEFVT
251 AAGITHGEFA STSLYKKAGS ENLYFQGLPA QVAFTPYAPE PGSTCRLREY
301 YDQTAQMCCS KCSPGQHAKV FCTKTSDTVC DSCEDSTYTQ LWNWVPECLS
351 CGSRCSSDQV ETQACTREQN RICTCRPGWY CALSKQEGCR LCAPLRKCRP
401 GFGVARPGTE TSDVVCKPCA PGTFSNTTSS TDICRPHQIC NVVAIPGNAS
451 MDAVCTSTSP TRSMAPGAVH LPQPVSTRSQ HTQPTPEPST APSTSFLLPM
501 GPSPPAEGST GDEPKSCDKT HTCPPCPAPE LLGGPSVFLF PPKPKDTLMI
551 SRTPEVTCVV VDVSHEDPEV KFNWYVDGVE VHNAKTKPRE EQYNSTYRVV
601 SVLTVLHQDW LNGKEYKCKV SNKALPAPIE KTISKAKGQP REPQVYTLPP
651 SREEMTKNQV SLTCLVKGFY PSDIAVEWES NGQPENNYKT TPPVLDSDGS
701 FFLYSKLTVD KSRWQQGNVF SCSVMHEALH NHYTQKSLSL SPGK

**Protein disulfide isomerase**
001 MKFSAGAVLS WSSLLLASSV FAQQEAVAPE DSAVVKLATD SFNEYIQSHD
051 LVLAEFFAPW CGHCKNMAPE YVKAAETLVE KNITLAQIDC TENQDLCMEH
101 NIPGFPSLKI FKNSDVNNSI DYEGPRTAEA IVQFMIKQSQ PAVAVVADLP
151 AYLANETFVT PVIVQSGKID ADFNATFYSM ANKHFNDYDF VSAENADDDF
201 KLSIYLPSAM DEPVVYNGKK ADIADADVFE KWLQVEALPY FGEIDGSVFA
251 QYVESGLPLG YLFYNDEEEL EEYKPLFTEL AKKNRGLMNF VSIDARKFGR
301 HAGNLNMKEQ FPLFAIHDMT EDLKYGLPQL SEEAFDELSD KIVLESKAIE
351 SLVKDFLKGD ASPIVKSQEI FENQDSSVFQ LVGKNHDEIV NDPKKDVLVL
401 YYAPWCGHCK RLAPTYQELA DTYANATSDV LIAKLDHTEN DVRGVVIEGY
451 PTIVLYPGGK KSESVVYQGS RSLDSLFDFI KENGHFDVDG KALYEEAQEK
501 AAEEADADAE LADEEDAISG SGENLYFQGS SFLVQSGMSK GEELFTGVVP
551 ILVELDGDVN GHKFSVRGEG EGDATNGKLT LKFICTTGKL PVPWPTLVTT
601 LTYGVQCFSR YPDHMKRHDF FKSAMPEGYV QERTISFKDD GTYKTRAEVK

651 FEGDTLVNRI ELKGIDFKED GNILGHKLEY NFNSHNVYIT ADKQKNGIKA
701 NFKIRHNVED GSVQLADHYQ QNTPIGDGPV LLPDNHYLST QSVLSKDPNE
751 KRDHMVLLEF VTAAGITHGM SGLNDIFEAQ KIEWHEHHHH HHHH

**CD16a**
001 HHHHHHHHMS GLNDIFEAQK IEWHEMSKGE ELFTGVVPIL VELDGDVNGH
051 KFSVRGEGEG DATNGKLTLK FICTTGKLPV PWPTLVTTLT YGVQCFSRYP
101 DHMKRHDFFK SAMPEGYVQE RTISFKDDGT YKTRAEVKFE GDTLVNRIEL
151 KGIDFKEDGN ILGHKLEYNF NSHNVYITAD KQKNGIKANF KIRHNVEDGS
201 VQLADHYQQN TPIGDGPVLL PDNHYLSTQS VLSKDPNEKR DHMVLLEFVT
251 AAGITHGEFS SENLYFQGRT EDLPKAVVFL EPQWYRVLEK DSVTLKCQGA
301 YSPEDNSTQW FHNESLISSQ ASSYFIDAAT VDDSGEYRCQ TNLSTLSDPV
351 QLEVHIGWLL LQAPRWVFKE EDPIHLRCHS WKNTALHKVT YLQNGKGRKY
401 FHHNSDFYIP KATLKDSGSY FCRGLVGSKN VSSETVNITI TQG

**SARS-CoV-2 spike glycoprotein**
0001 MFVFLVLLPL VSSQCVNLTT RTQLPPAYTN SFTRGVYYPD KVFRSSVLHS
0051 TQDLFLPFFS NVTWFHAIHV SGTNGTKRFD NPVLPFNDGV YFASTEKSNI
0101 IRGWIFGTTL DSKTQSLLIV NNATNVVIKV CEFQFCNDPF LGVYYHKNNK
0151 SWMESEFRVY SSANNCTFEY VSQPFLMDLE GKQGNFKNLR EFVFKNIDGY
0201 FKIYSKHTPI NLVRDLPQGF SALEPLVDLP IGINITRFQT LLALHRSYLT
0251 PGDSSSGWTA GAAAYYVGYL QPRTFLLKYN ENGTITDAVD CALDPLSETK
0301 CTLKSFTVEK GIYQTSNFRV QPTESIVRFP NITNLCPFGE VFNATRFASV
0351 YAWNRKRISN CVADYSVLYN SASFSTFKCY GVSPTKLNDL CFTNVYADSF
0401 VIRGDEVRQI APGQTGKIAD YNYKLPDDFT GCVIAWNSNN LDSKVGGNYN
0451 YLYRLFRKSN LKPFERDIST EIYQAGSTPC NGVEGFNCYF PLQSYGFQPT
0501 NGVGYQPYRV VVLSFELLHA PATVCGPKKS TNLVKNKCVN FNFNGLTGTG
0551 VLTESNKKFL PFQQFGRDIA DTTDAVRDPQ TLEILDITPC SFGGVSVITP
0601 GTNTSNQVAV LYQDVNCTEV PVAIHADQLT PTWRVYSTGS NVFQTRAGCL
0651 IGAEHVNNSY ECDIPIGAGI CASYQTQTNS PGGSGSVASQ SIIAYTMSLG
0701 AENSVAYSNN SIAIPTNFTI SVTTEILPVS MTKTSVDCTM YICGDSTECS
0751 NLLLQYGSFC TQLNRALTGI AVEQDKNTQE VFAQVKQIYK TPPIKDFGGF
0801 NFSQILPDPS KPSKRSFIED LLFNKVTLAD AGFIKQYGDC LGDIAARDLI
0851 CAQKFNGLTV LPPLLTDEMI AQYTSALLAG TITSGWTFGA GAALQIPFAM
0901 QMAYRFNGIG VTQNVLYENQ KLIANQFNSA IGKIQDSLSS TASALGKLQD
0951 VVNQNAQALN TLVKQLSSNF GAISSVLNDI LSRLDPPEAE VQIDRLITGR
1001 LQSLQTYVTQ QLIRAAEIRA SANLAATKMS ECVLGQSKRV DFCGKGYHLM
1051 SFPQSAPHGV VFLHVTYVPA QEKNFTTAPA ICHDGKAHFP REGVFVSNGT
1101 HWFVTQRNFY EPQIITTDNT FVSGNCDVVI GIVNNTVYDP LQPELDSFKE
1151 ELDKYFKNHT SPDVDLGDIS GINASVVNIQ KEIDRLNEVA KNLNESLIDL
1201 QELGKYEQGS GGYIPEAPRD GQAYVRKDGE WVLLSTFLGG SHHHHHH

#### S1b: CD16a purification

**CD16A: Expression and purification in Lec1 cells**

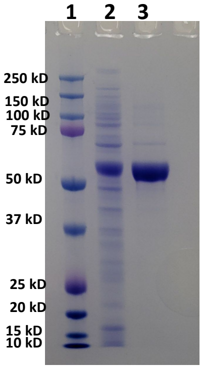


Lane 1: Marker
Lane 2: Crude :CD16A
Lane 3: Ni-NTA elution: CD16A

**CD16A: Expression and purification HEK 293f WT cells**

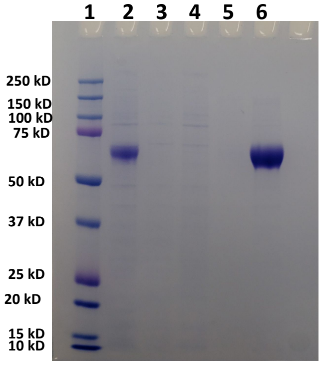


Lane 1: Marker
Lane 2: Crude media CD16A
Lane 3: Flow through
Lane 4: Wash 1
Lane 5: Wash 2
Lane 6: CD16A– Ni-NTA elution

#### S1c: PDI purification

**PDI: Expression and purification in Lec1 cells**

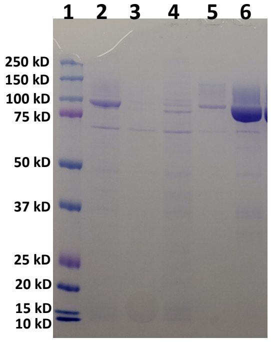


Lane 1: Marker
Lane 2: Crude Media : PDI
Lane 3: Flow through
Lane 4: Wash 1
Lane 5: Wash 2
Lane 6: Ni-NTA elution: PDI

**PDI: Expression and purification HEK 293f WT cells**

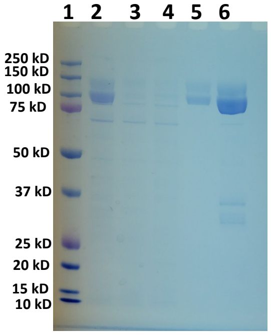


Lane 1: Marker
Lane 2: Crude media: PDI
Lane 3: Flow Through Ni-NTA
Lane 4: Wash 1
Lane 5: Wash 2
Lane 6: Ni-NTA elution: PDI

#### S1d: Etanercept purification

**ETANERCEPT: Expression and purification in Lec1 cells**

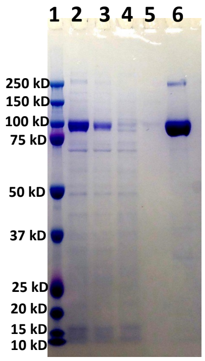


Lane 1: Marker
Lane 2: Crude media: ETNARCEPT
Lane 3: Flow Through Ni-NTA
Lane 4: Wash 1
Lane 5: Wash 2
Lane 6: Ni-NTA elution: ETANERCEPT

**ETANERCEPT: Expression and purification HEK 293f WT cells**

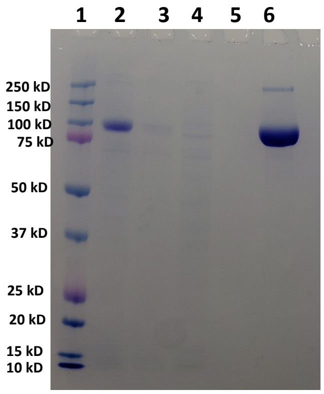


Lane 1: Marker
Lane 2: Crude media : ETANERCEPT
Lane 3: Flow through
Lane 4: Wash 1
Lane 5: Wash 2
Lane 6: Ni-NTA elution : ETANERCEPT

#### S1e: Spike purification

**Spike: Expression and Purification in Lec1 cells**

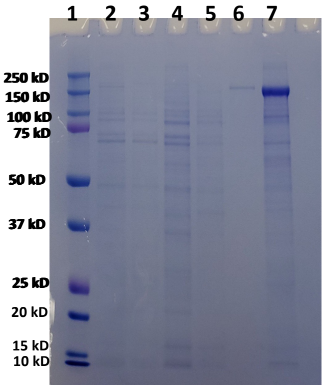


Lane 1: Marker
Lane 2: Crude media : Spike
Lane 3: Flow through
Lane 4: Wash 1
Lane 5: Wash 2
Lane 6: Ni-NTA elution
Lane 7: Spike After concentration and buffer exchange

**Spike: Expression and Purification in HEK 293FWT cells**

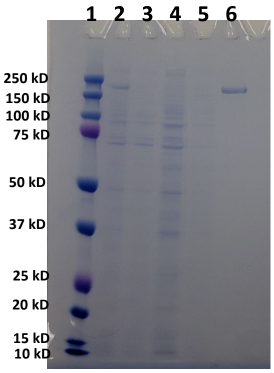


Lane 1: Marker
Lane 2: Crude media : Spike
Lane 3: Flow through
Lane 4: Wash 1
Lane 5: Wash 2
Lane 6: Ni-NTA elution: Spike

#### S1f: Enzyme purification

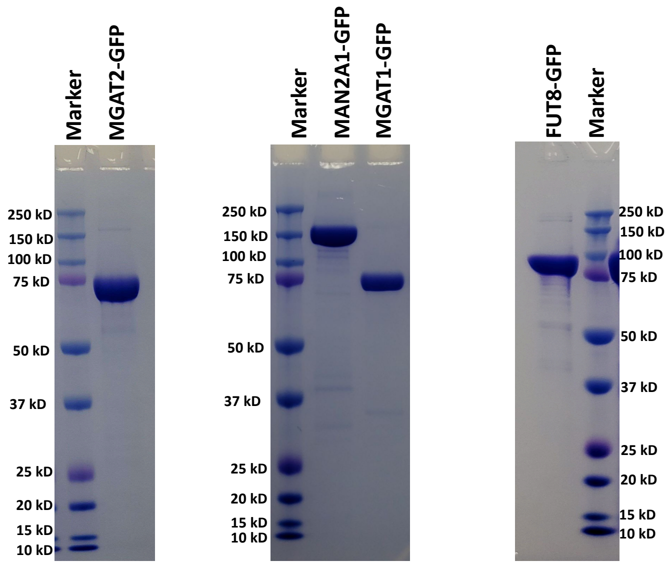

#### S2: SARS-CoV-2 spike glycoprotein site occupancy

a) SARS-CoV-2 spike glycoprotein expressed in WT-HEK293F cells

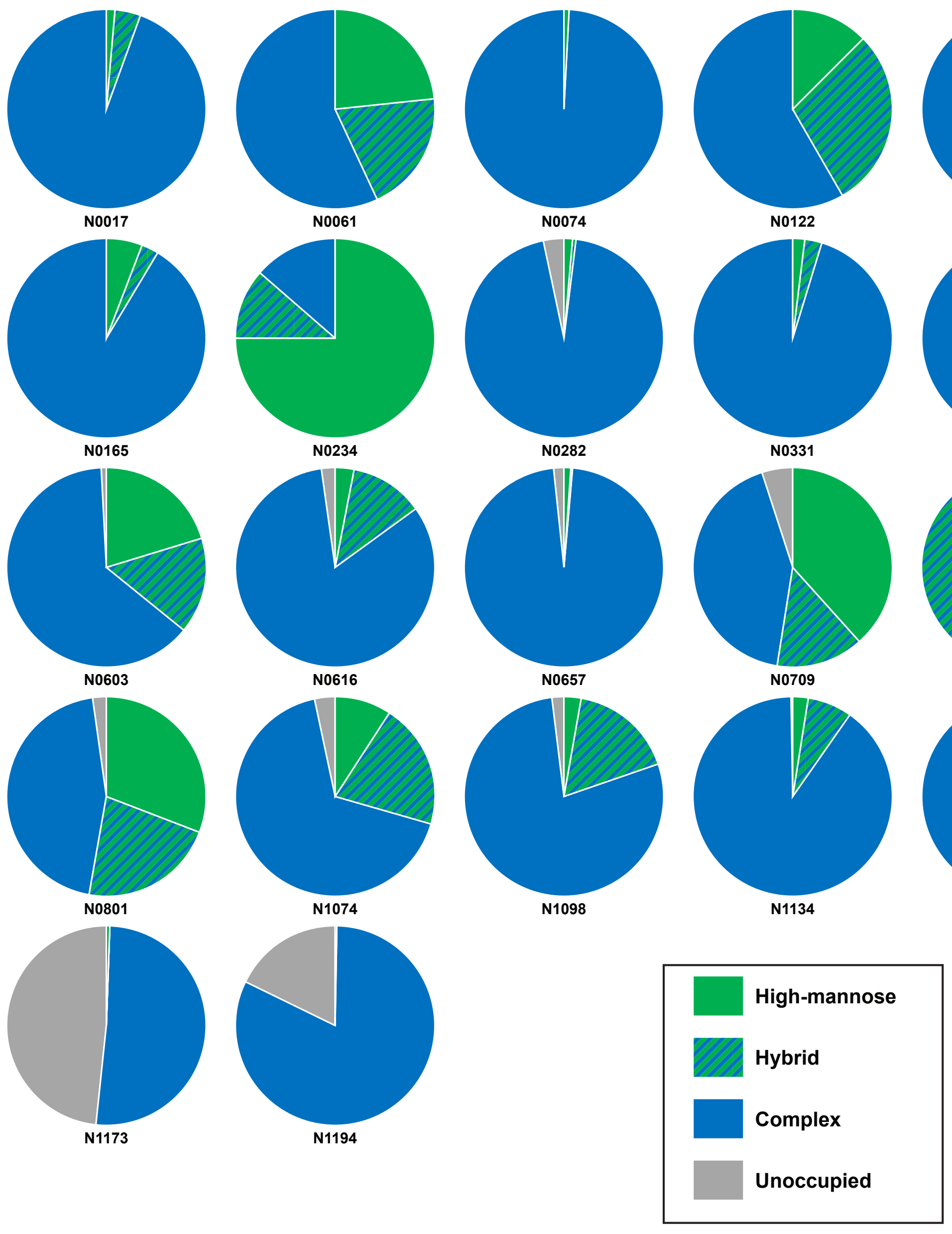

b) SARS-CoV-2 spike glycoprotein expressed in Lec1-HEK293F cells

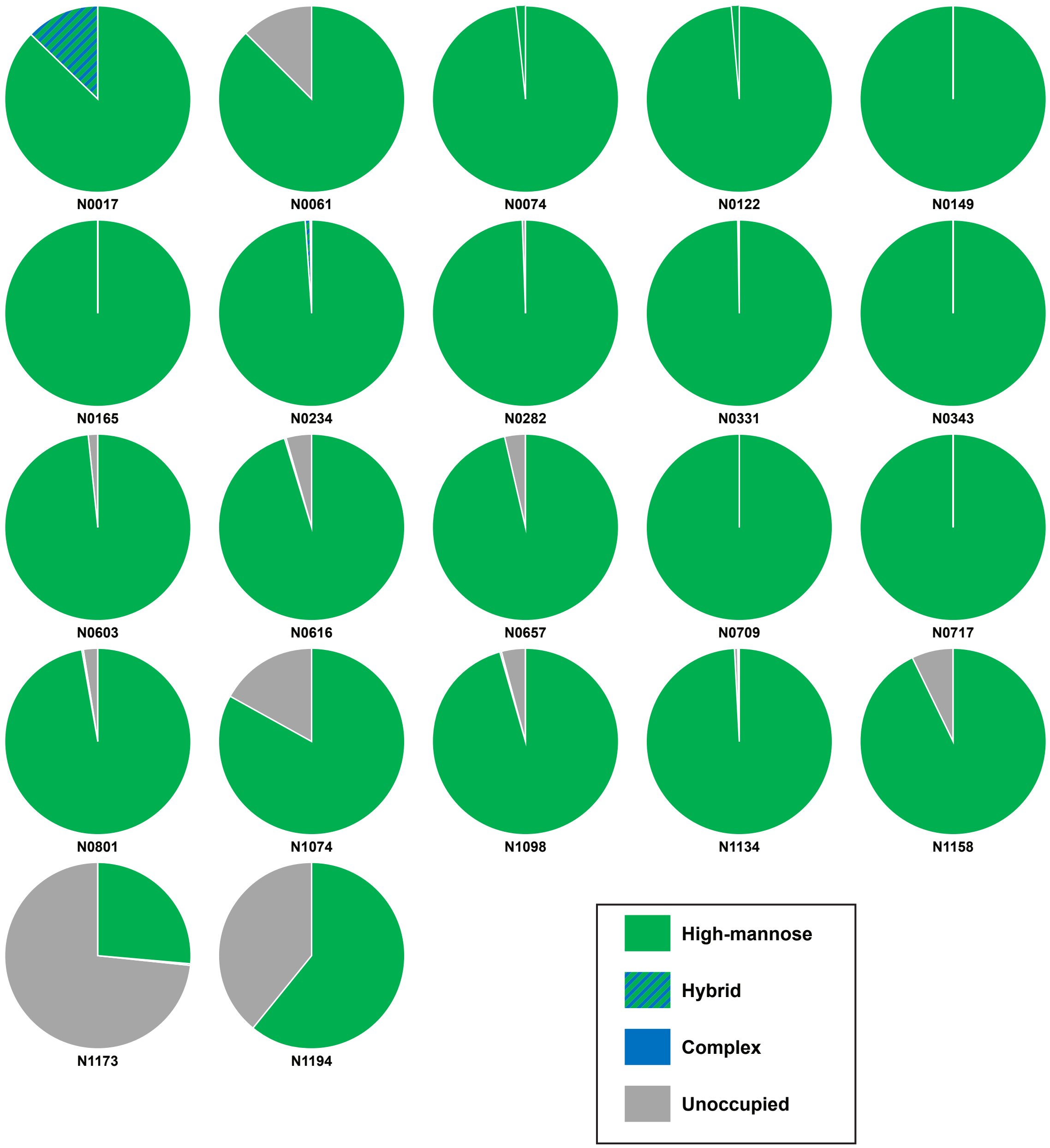


Figure S2: Site occupancy of SARS-CoV-2 spik glycoprotein expressed in a, WT-HEK293F and b, Lec1-HEK293F cells. Relative populations were ascertained with glycopeptide analysis and quantified with spectral counts.

#### S3: Site-specific N-glycan processing rates of SARS-CoV-2 spike glycoprotein

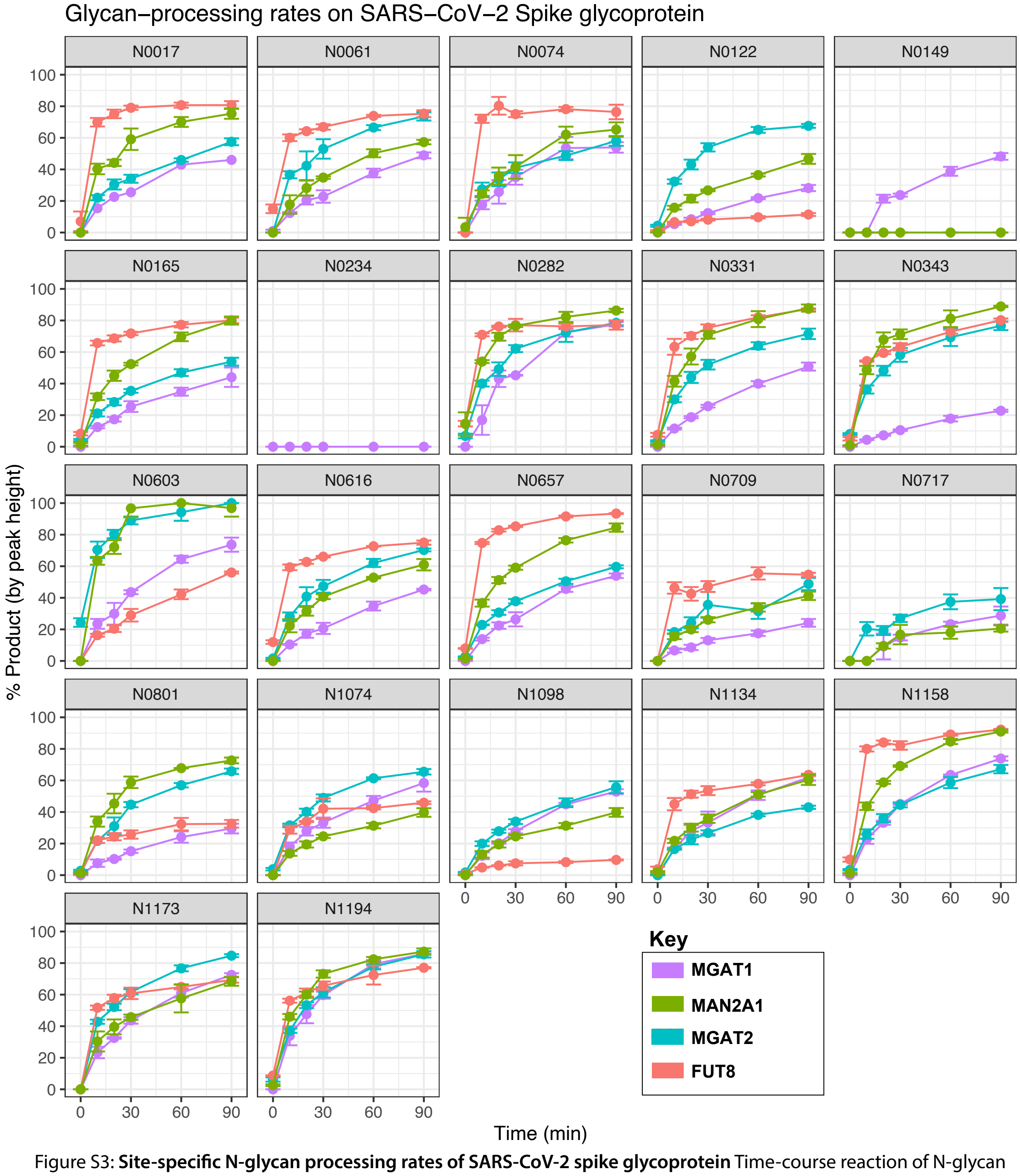


Figure S3: **Site-specific N-glycan processing rates of SARS-CoV-2 spike glycoprotein** Time-course reaction of N-glycan processing steps with recombinant enzymes. Reaction progress calculated as proportion of the sum of monoisotpic peak heights of product vs. the sum of product and reactant peak heights. Experiments performed in triplicate, error bars represent standard error of the mean.

### S4: Levels of GlcNAc3Man4 intermediate during MAN2A1 processing

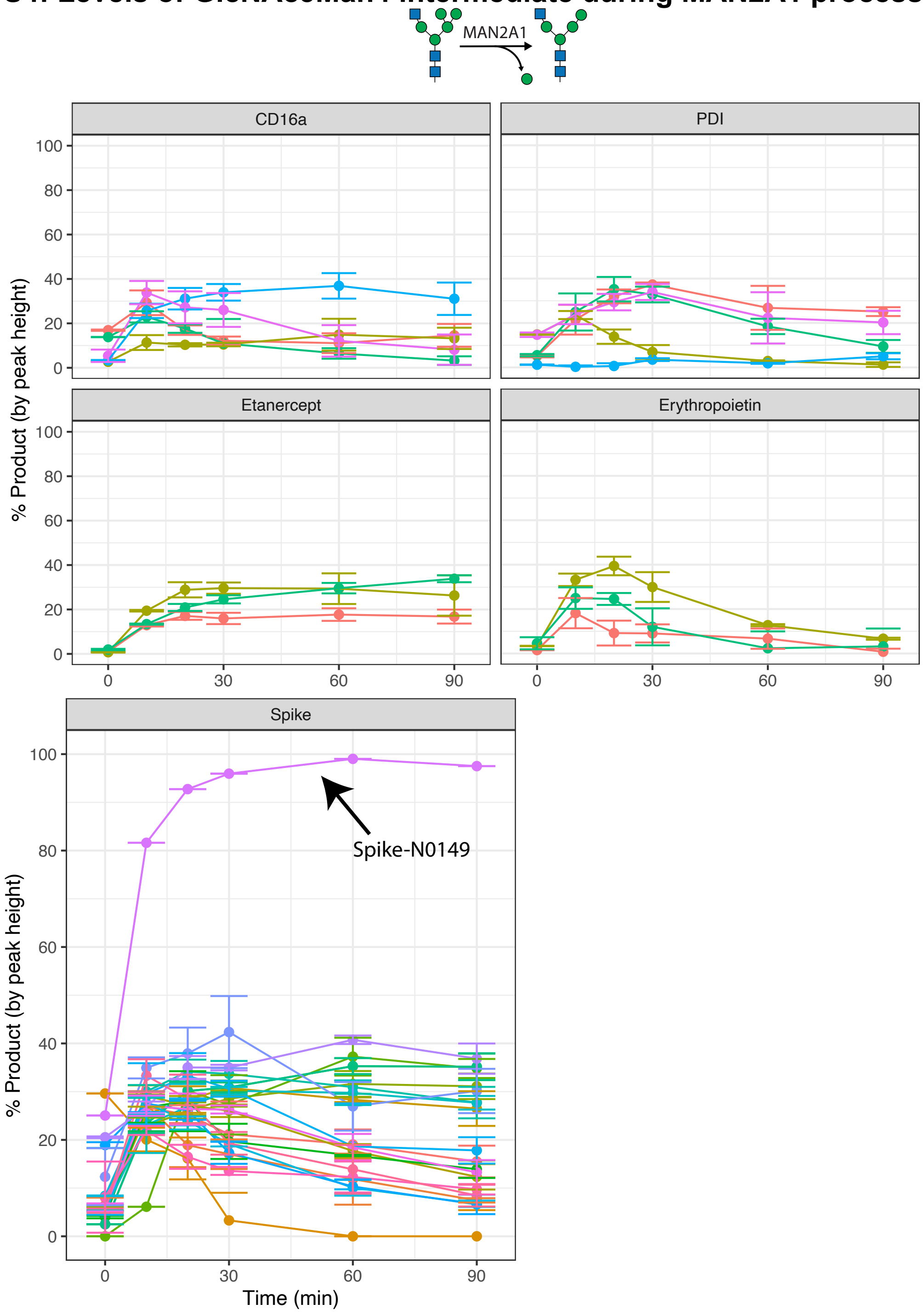


Figure S4: **Site-specific monitoring of production of MAN2A1 intermediate ($GlcNAc_3Man_4$).** Time-course reaction of mannose trimming of reporter proteins with recombinant MAN2A1. Legend omitted for SARS-CoV-2 spike glycoprotein due to large number of sites. Reaction progress calculated as proportion of the sum of monoisotpic peak heights of intermediate vs. the sum of product, intermediate, and reactant peak heights. Experiments performed in triplicate, error bars represent standard error of the mean.

#### S5: Site occupancy of CD16a Sequon 1 (N056) expressed in HEK293F cells.

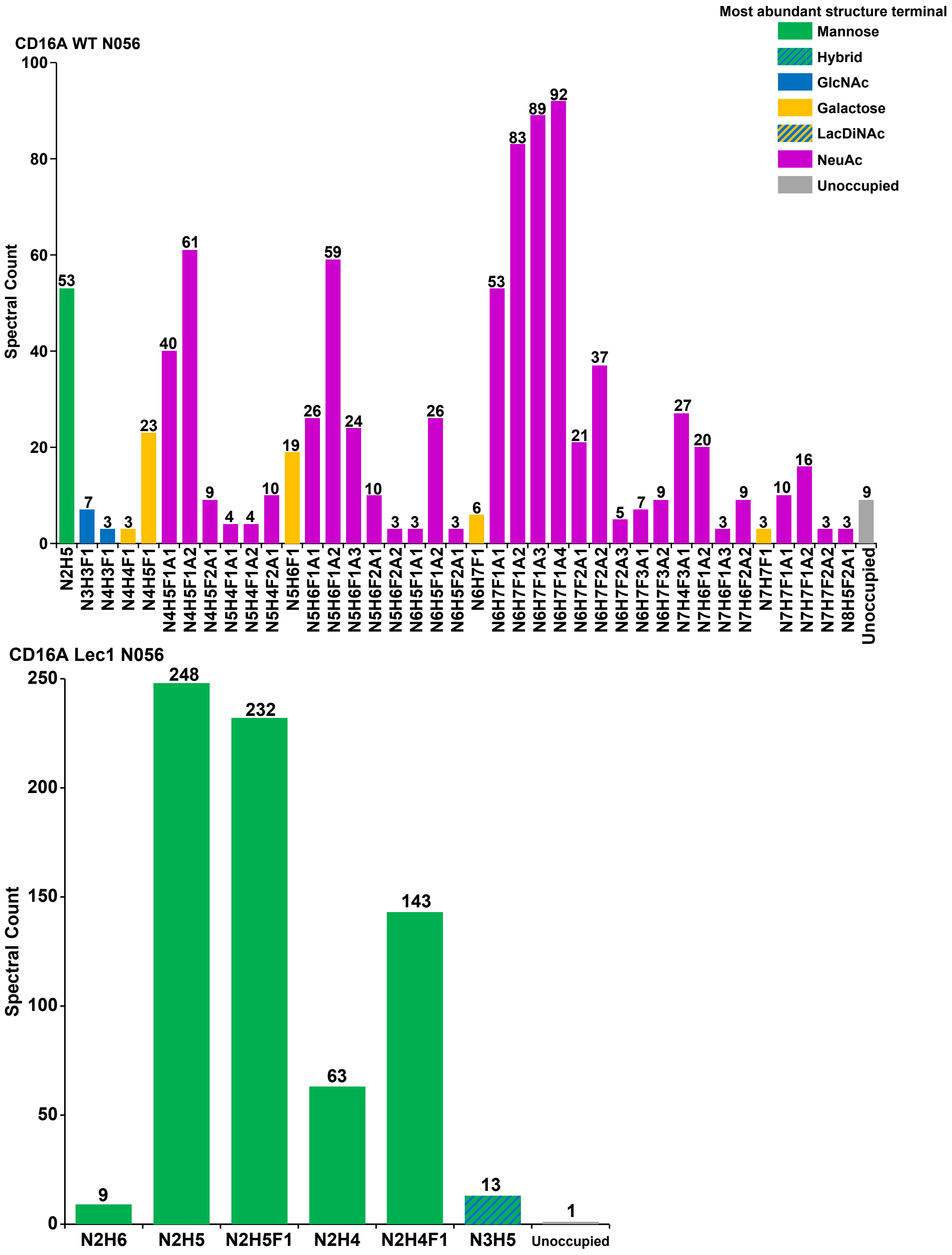

### S6: Site occupancy of CD16a Sequon 2 (N063) expressed in HEK293F cells.

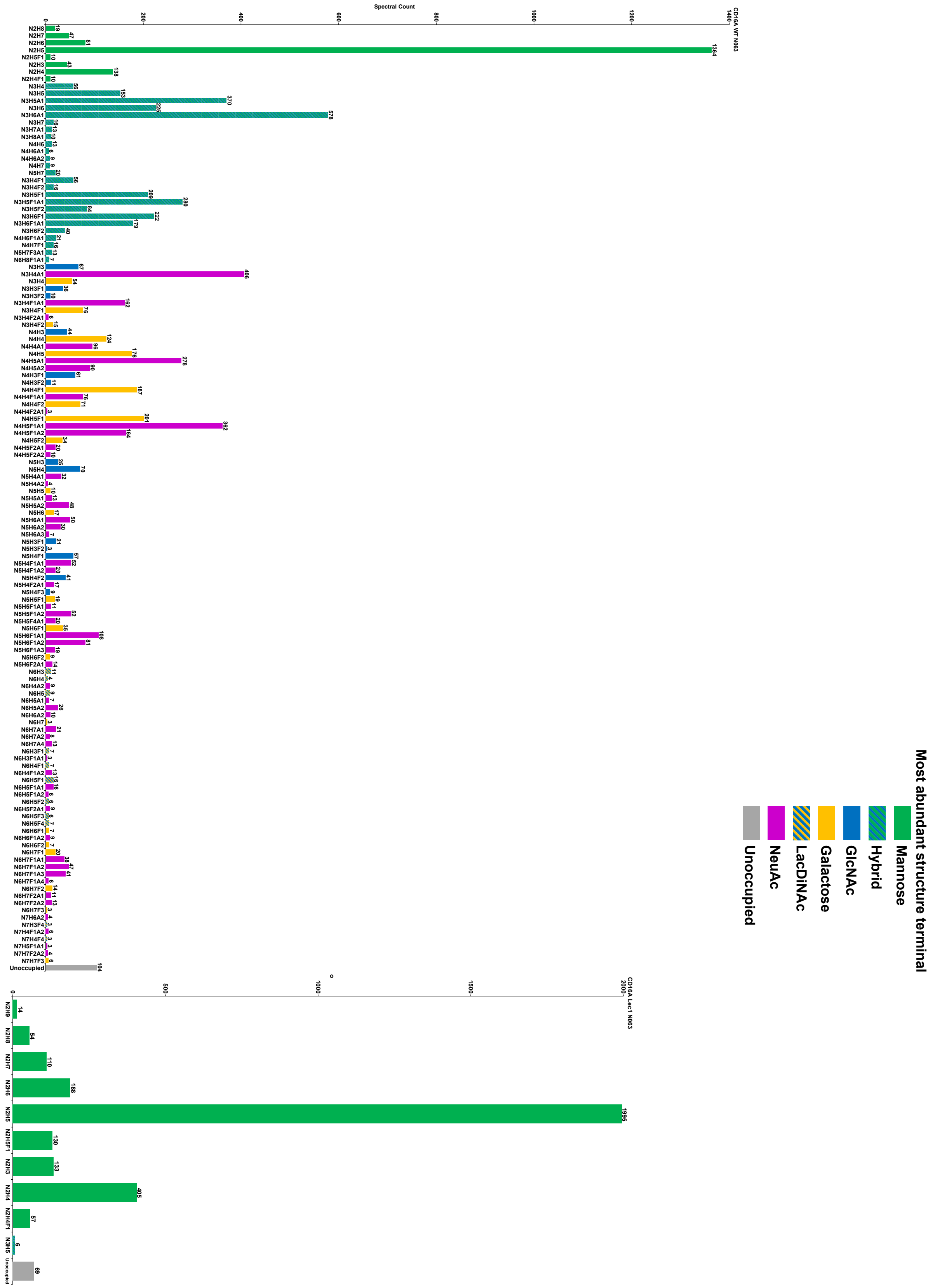

### S7: Site occupancy of CD16a Sequon 3 (N092) expressed in HEK293F cells.

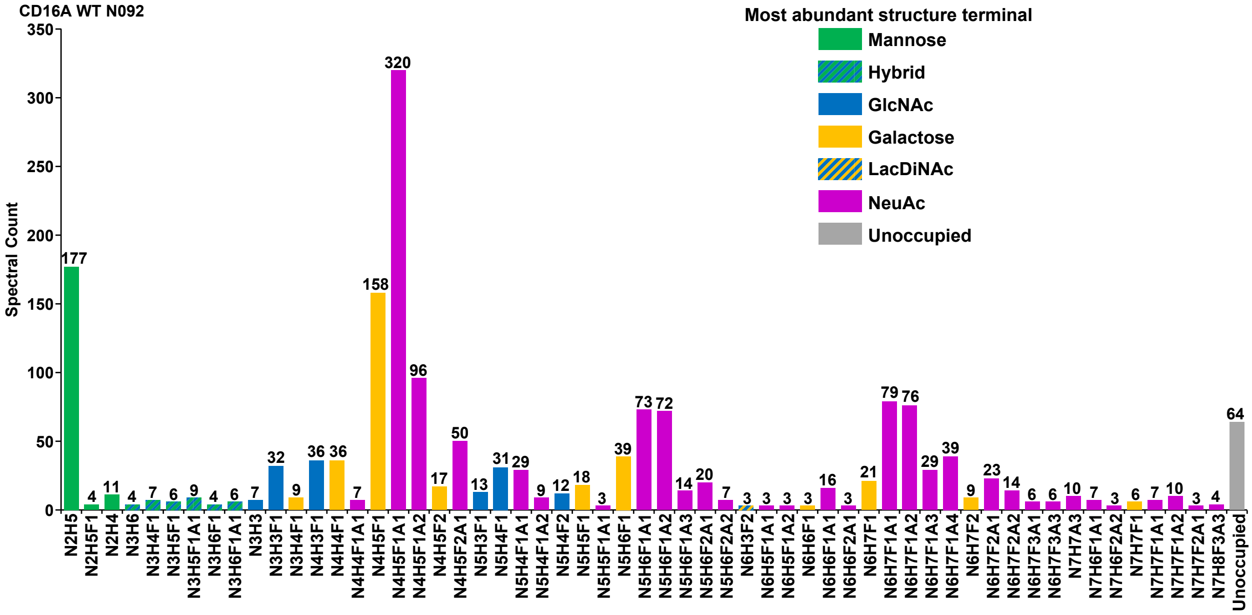


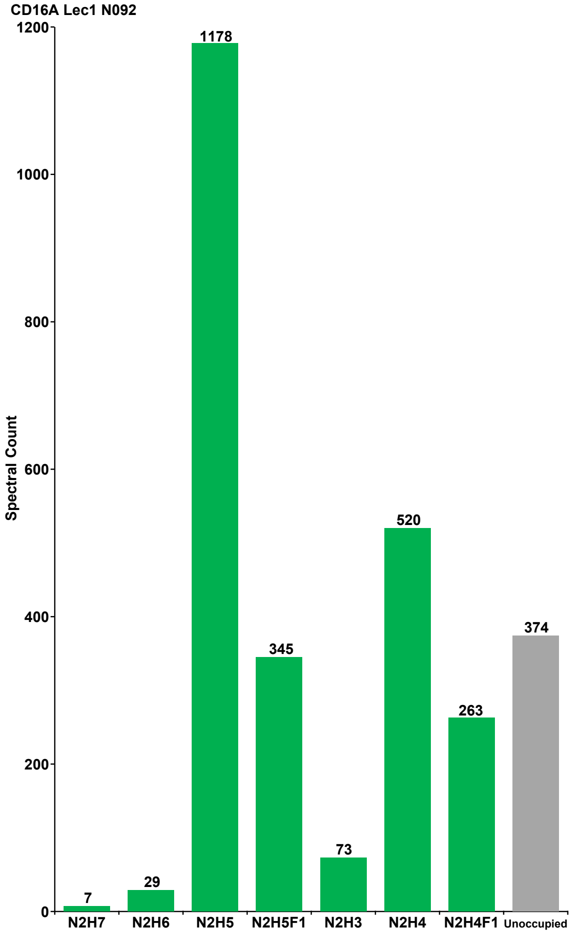

**S8: Site occupancy of CD16a Sequon 4 (N180) expressed in HEK293F cells.**

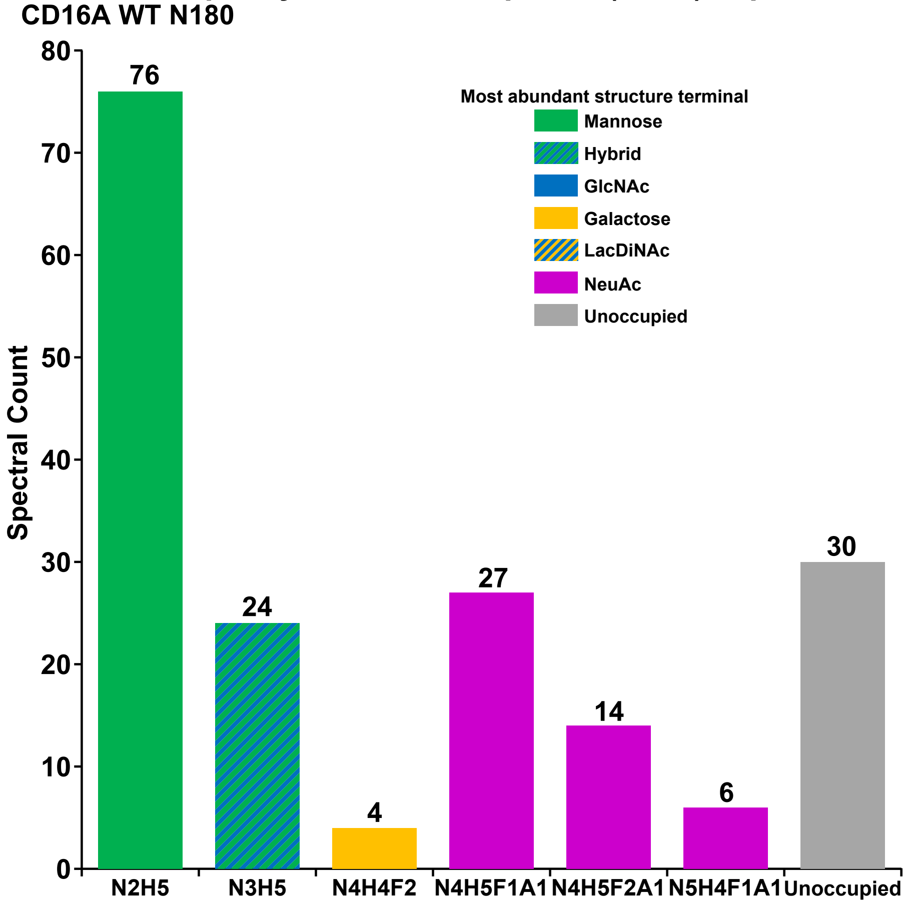


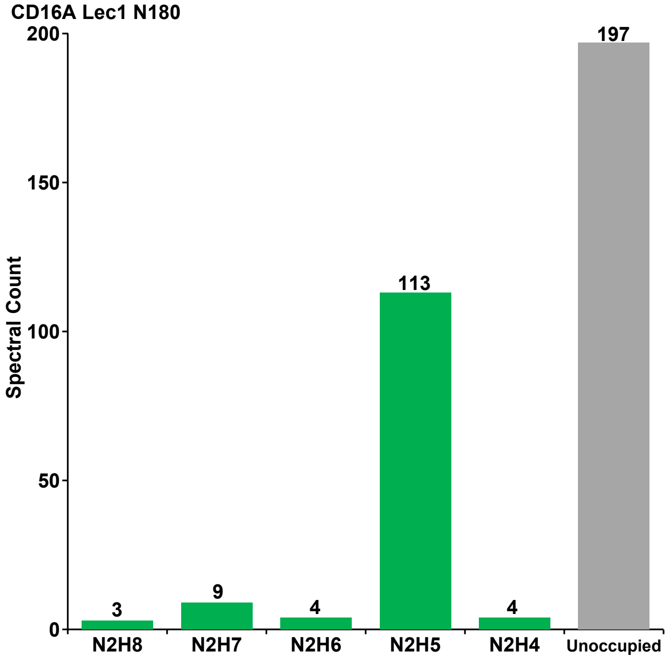

### S9: Site occupancy of CD16a Sequon 5 (N187) expressed in HEK293F cells.

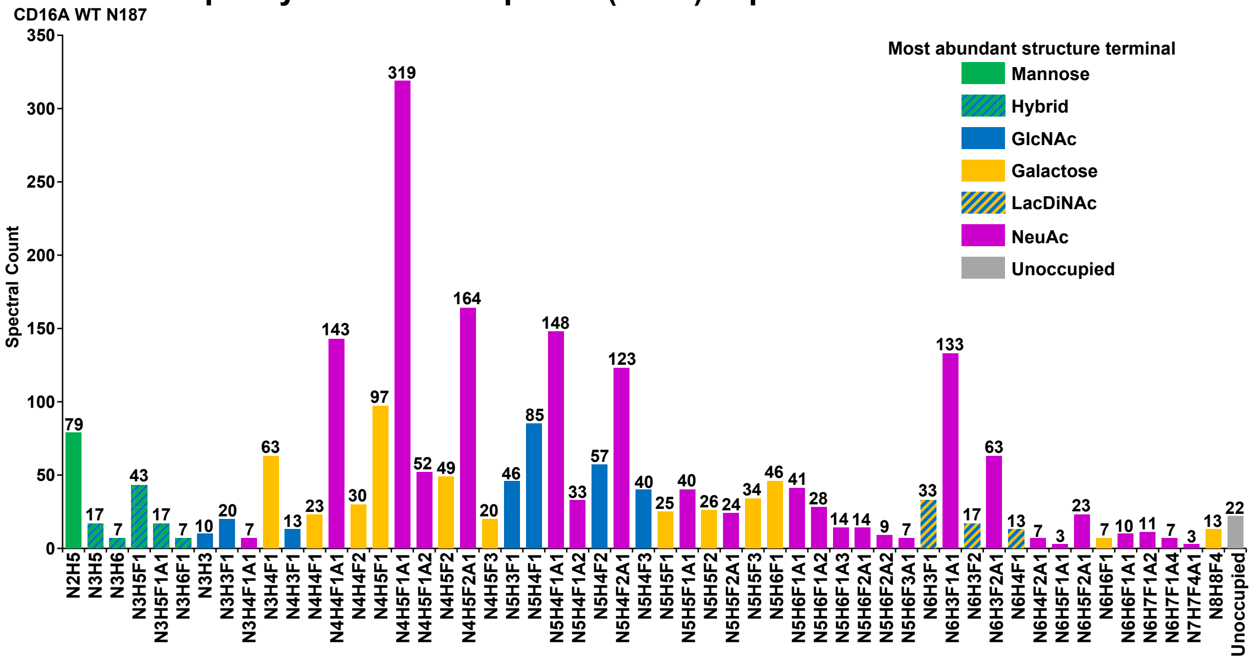


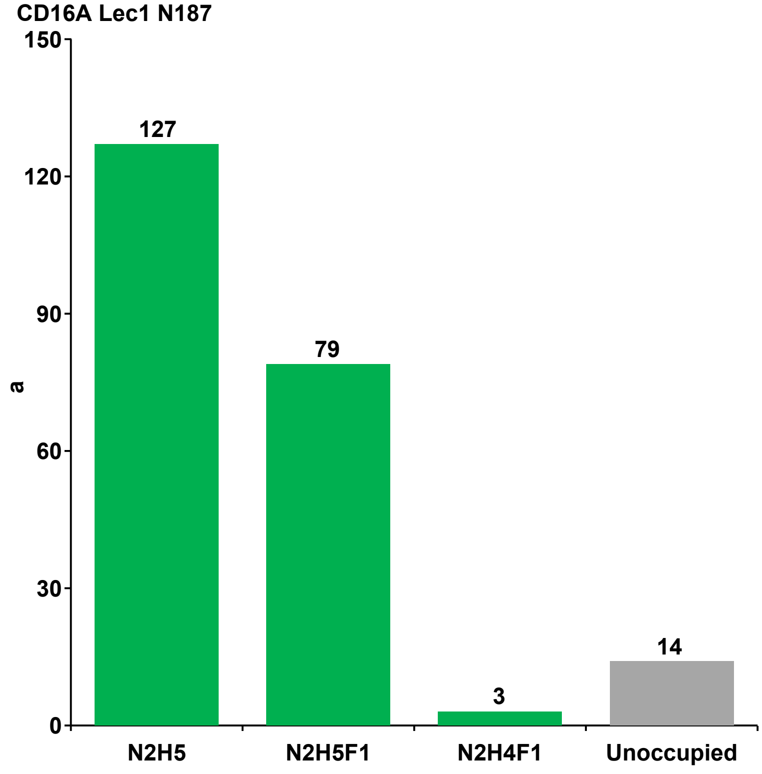

#### S10: Site occupancy of PDI Sequon 1 (N082) expressed in HEK293F cells

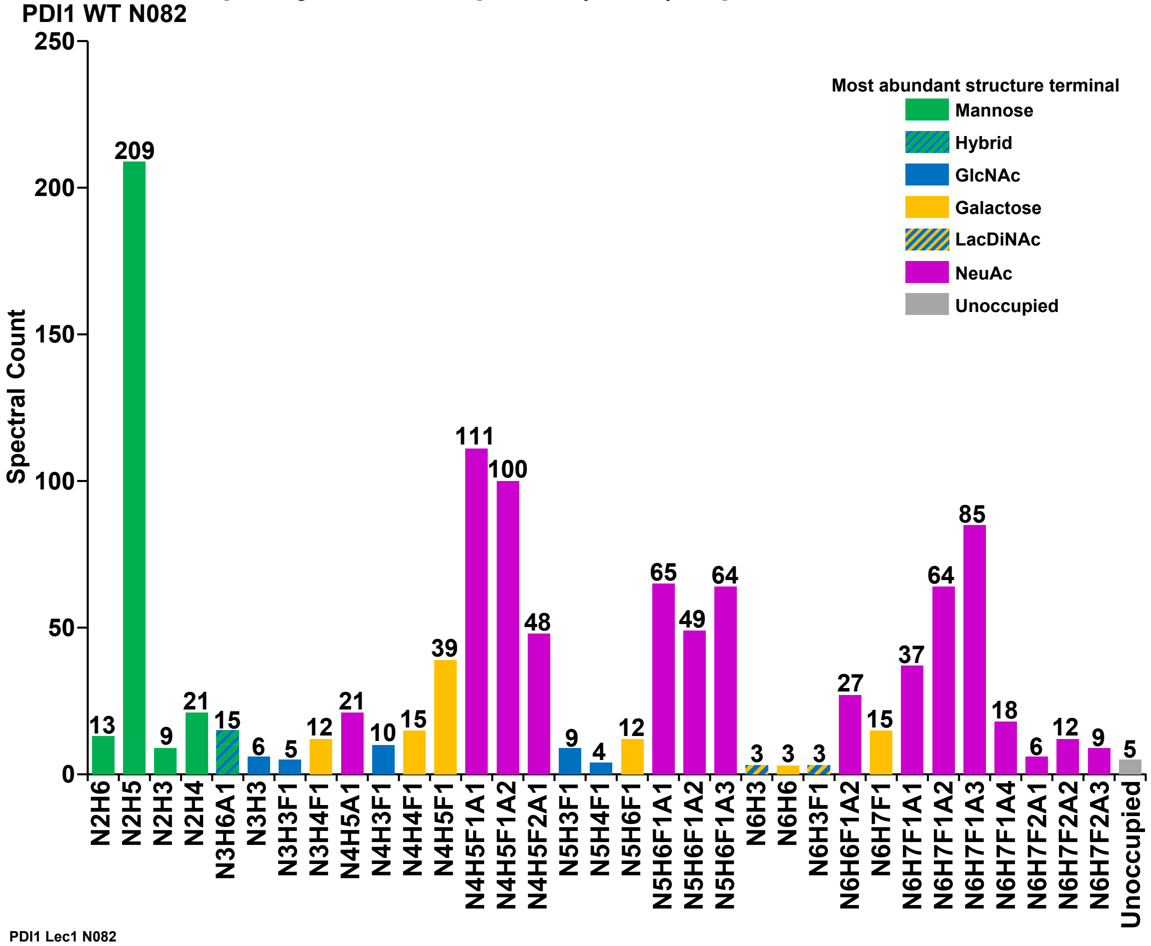


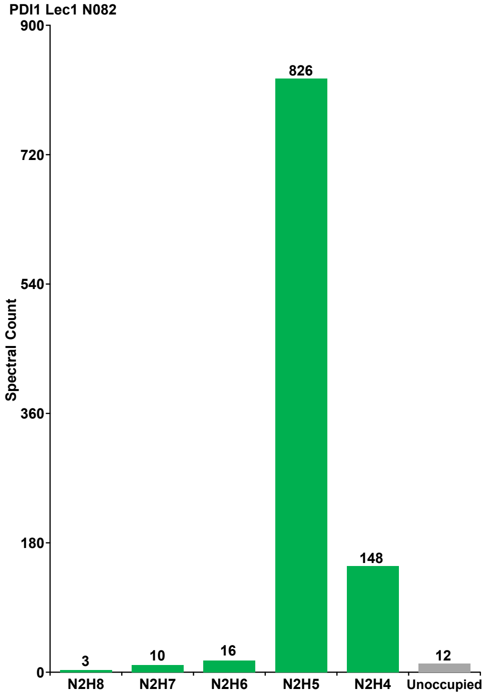

### S11: Site occupancy of PDI Sequon 2 (N187) expressed in HEK293F cells

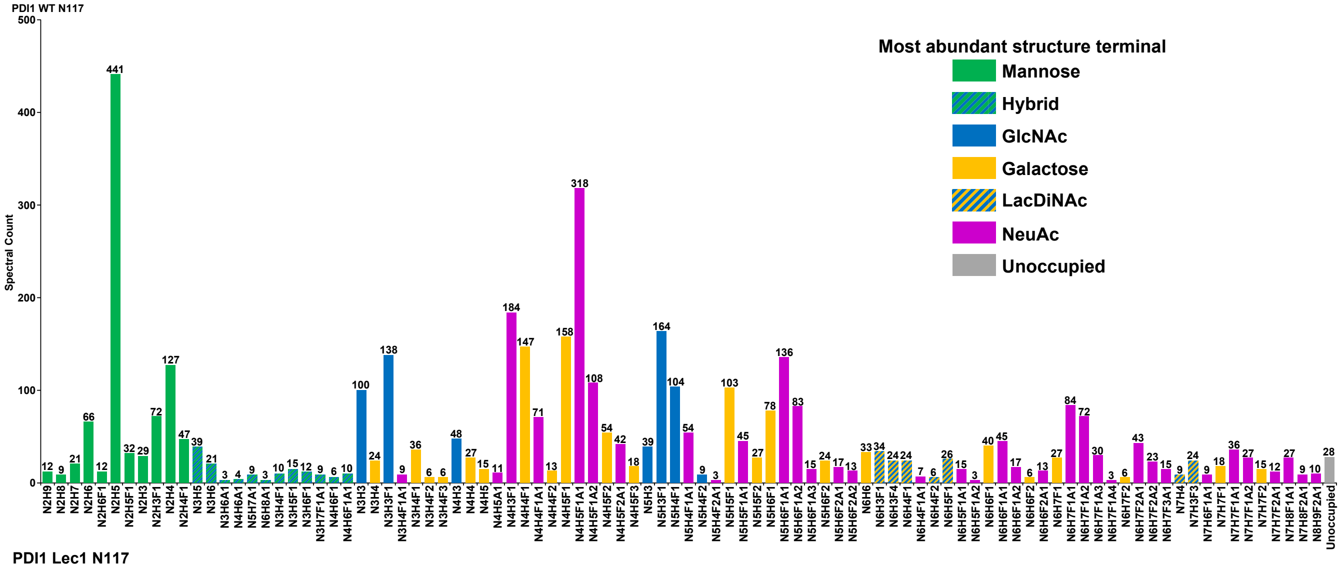


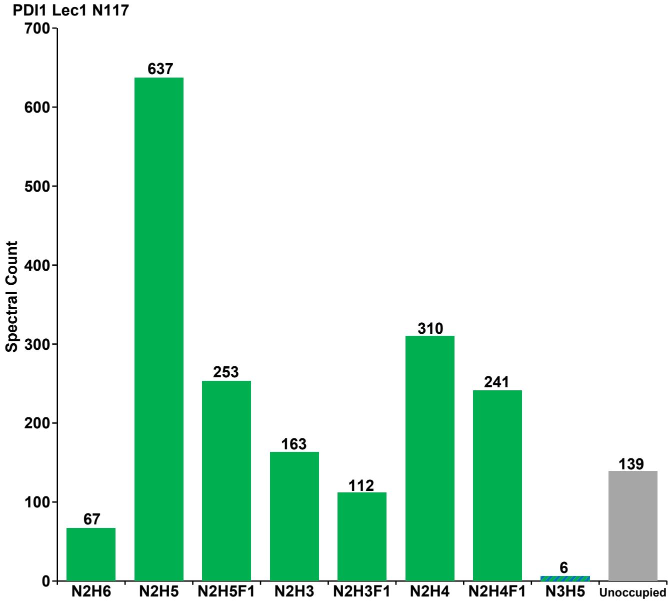

### S12: Site occupancy of PDI Sequon 3 (N155) expressed in HEK293F cells

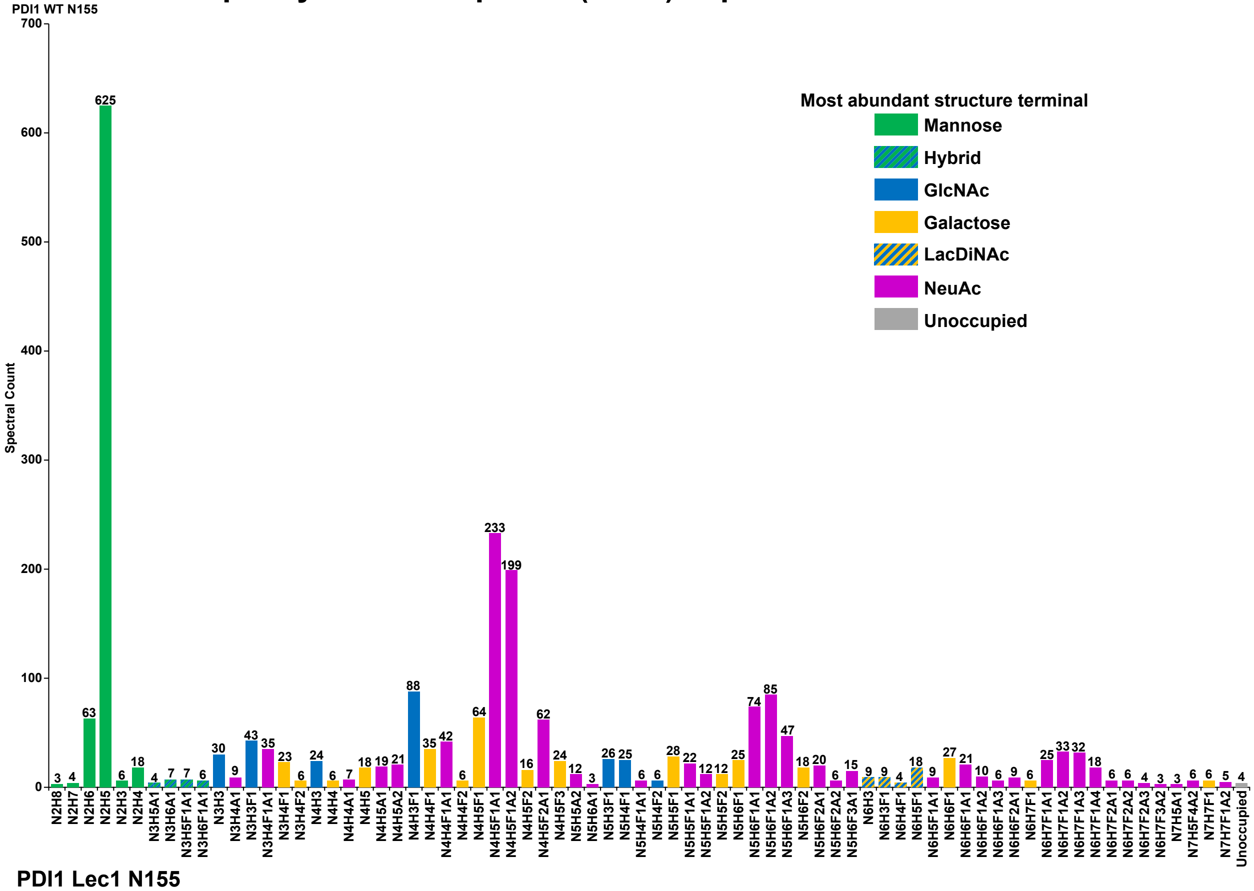


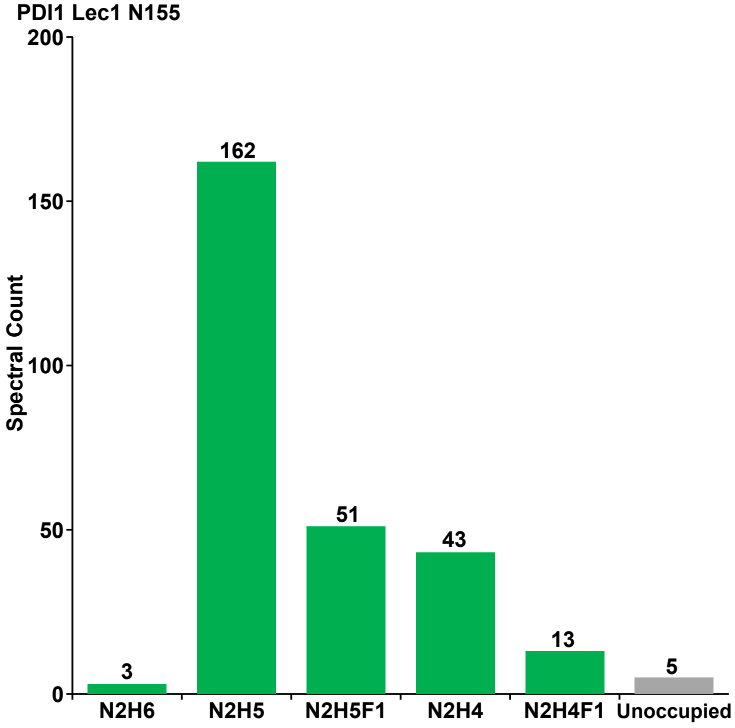

### S13: Site occupancy of PDI Sequon 4 (N174) expressed in HEK293F cells

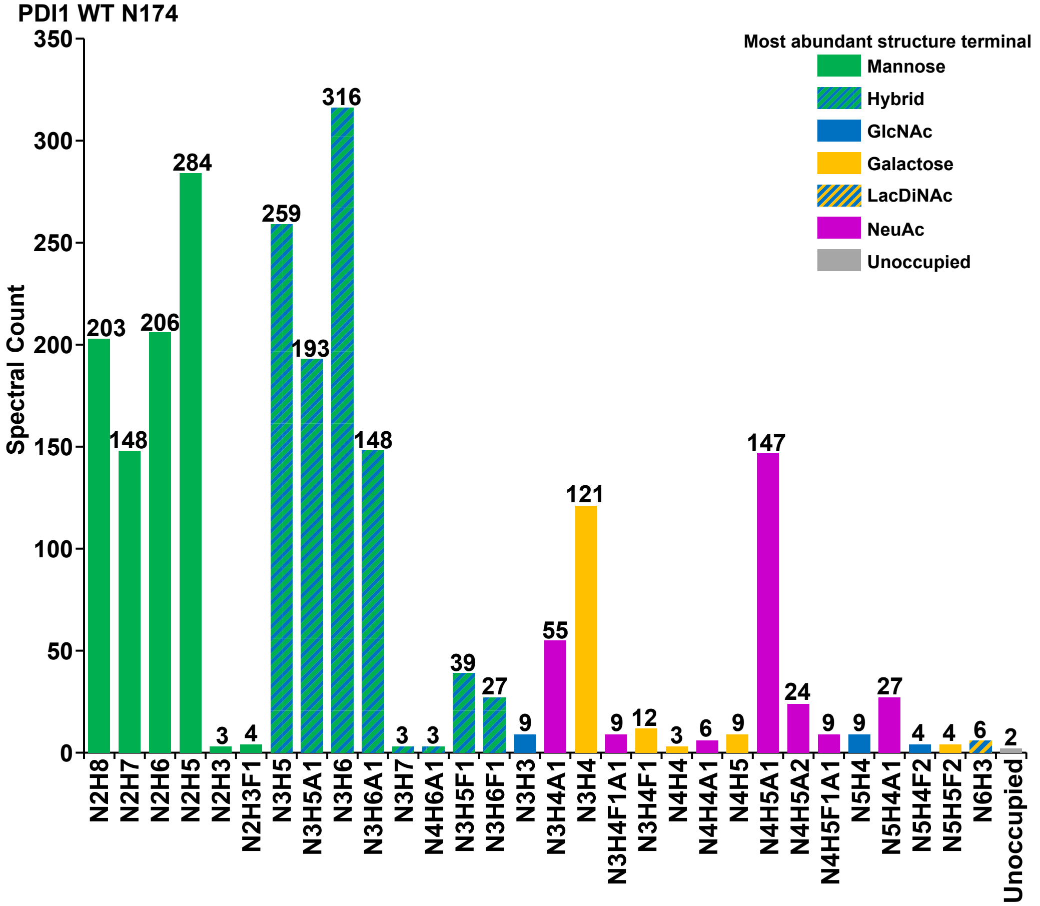


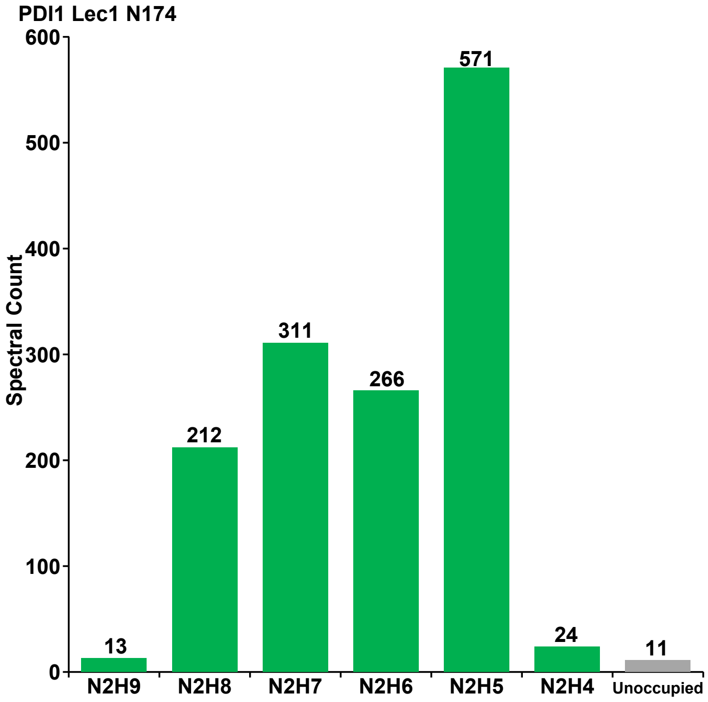

### S14: Site occupancy of PDI Sequon 5 (N425) expressed in HEK293F cells

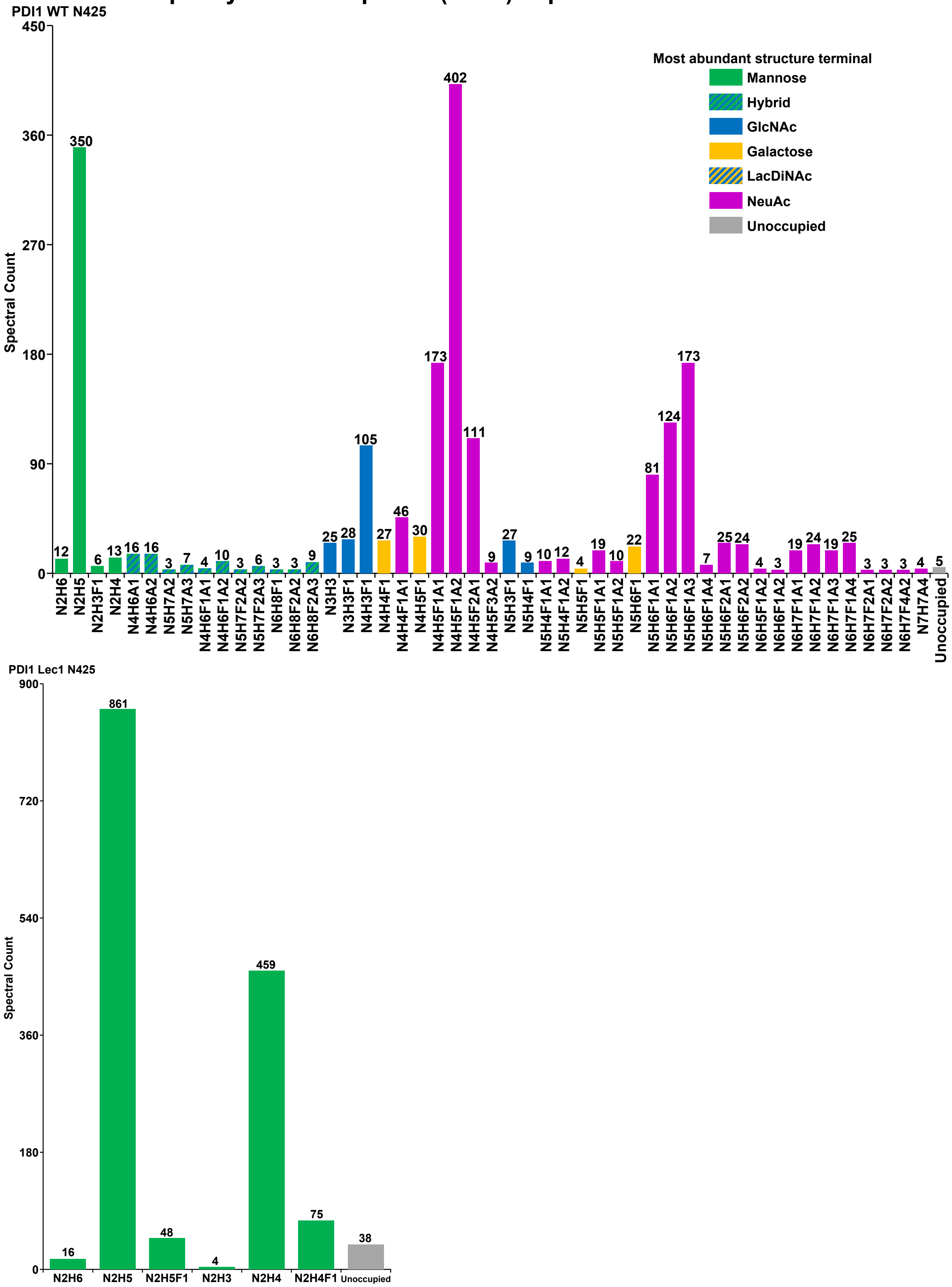

### S15: Site occupancy of etanercept sequon 1 (N149) expressed in HEK293F cells

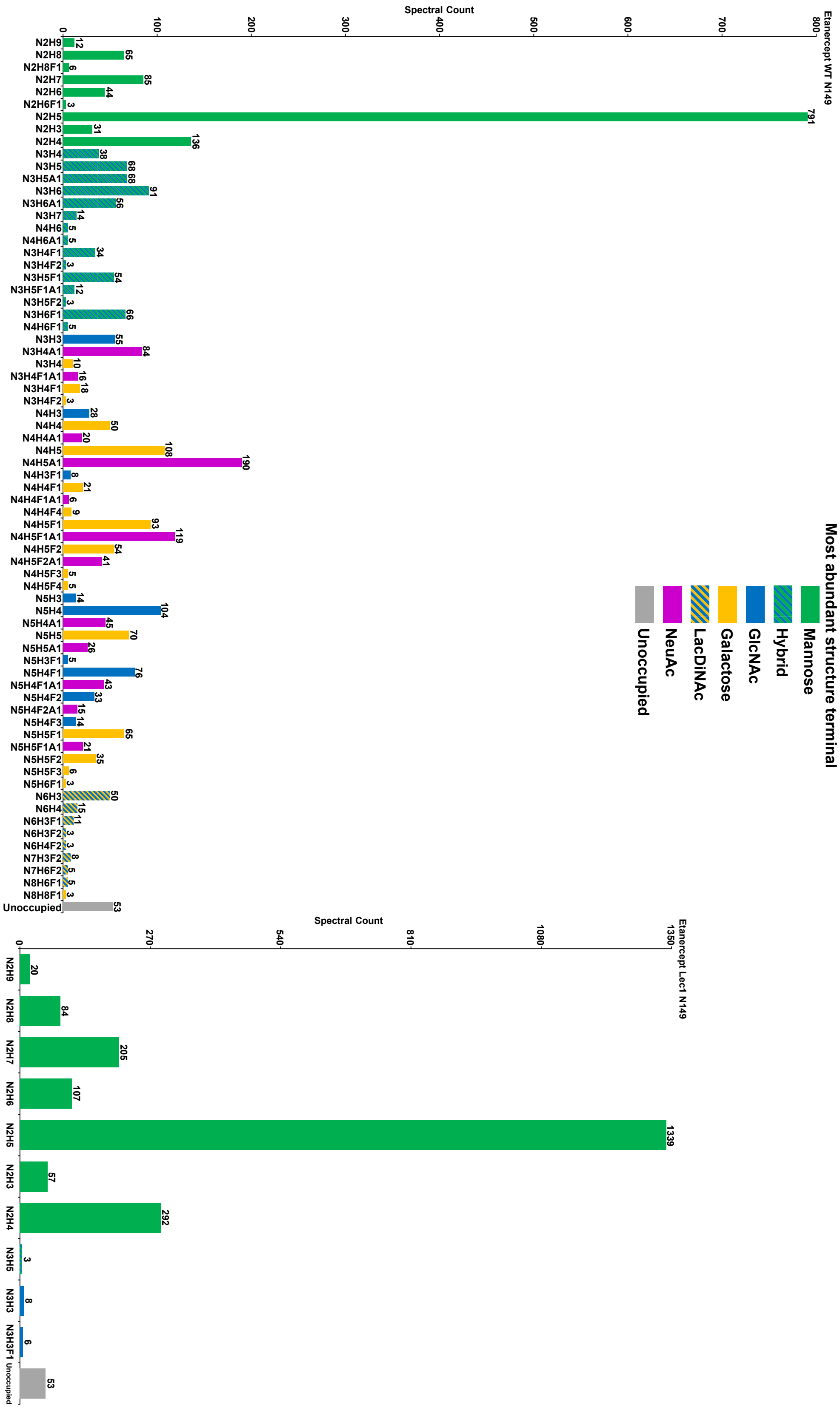

### S16: Site occupancy of etanercept sequon 2 (N171) expressed in HEK293F cells

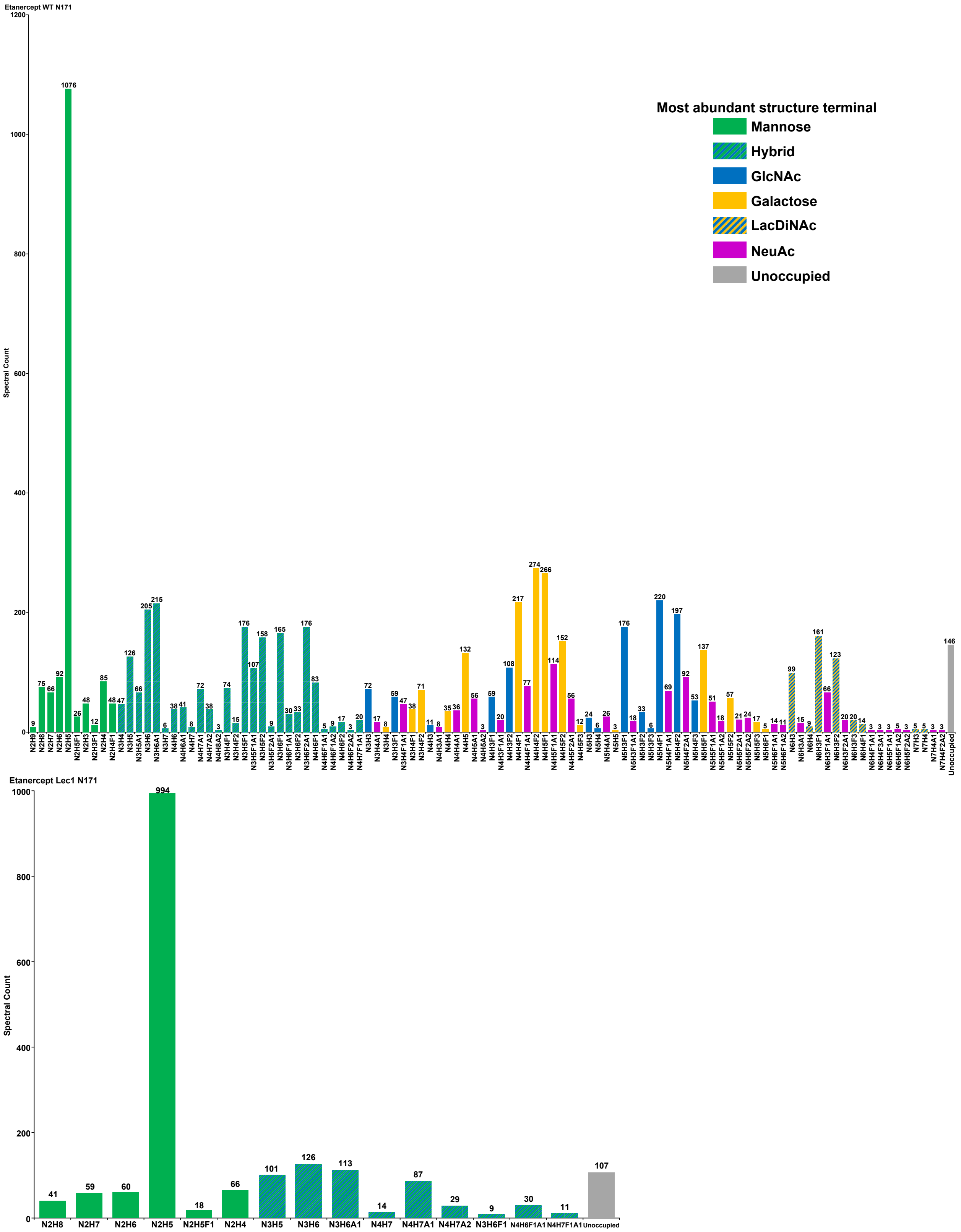

### S17: Site occupancy of etanercept sequon 3 (N317) expressed in HEK293F cells

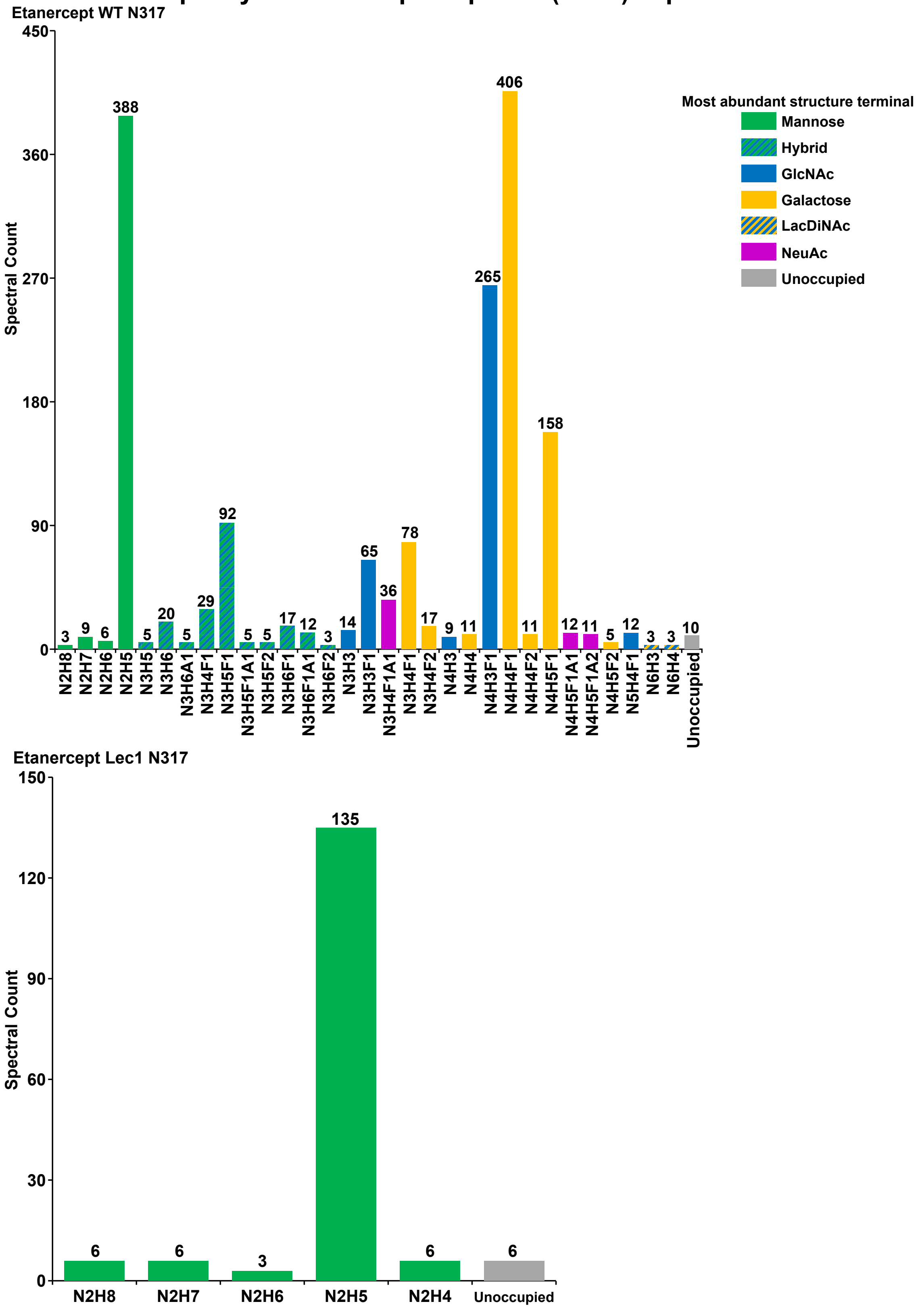

### S18: Site occupancy of erythropoietin sequon 1 (N051) expressed in HEK293F cells

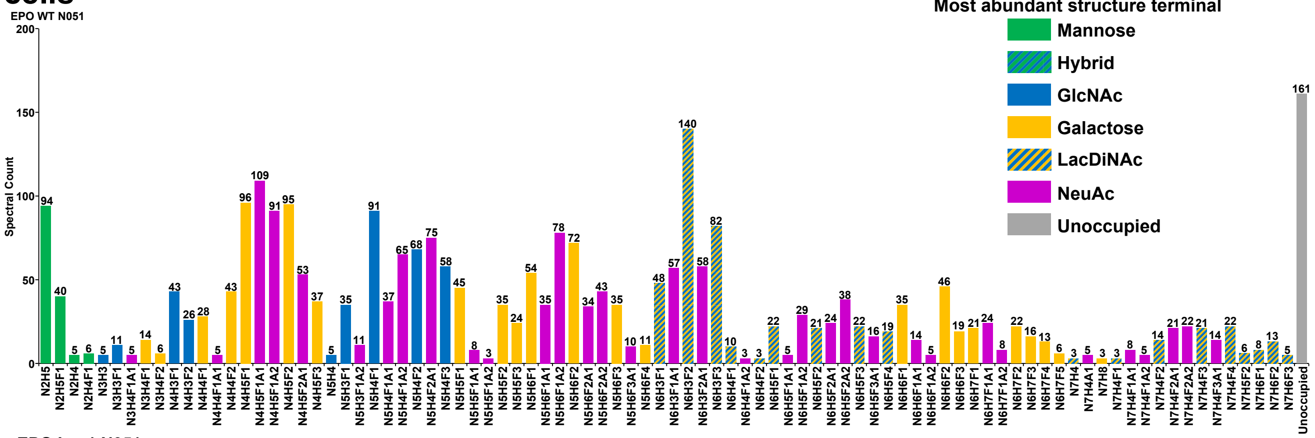


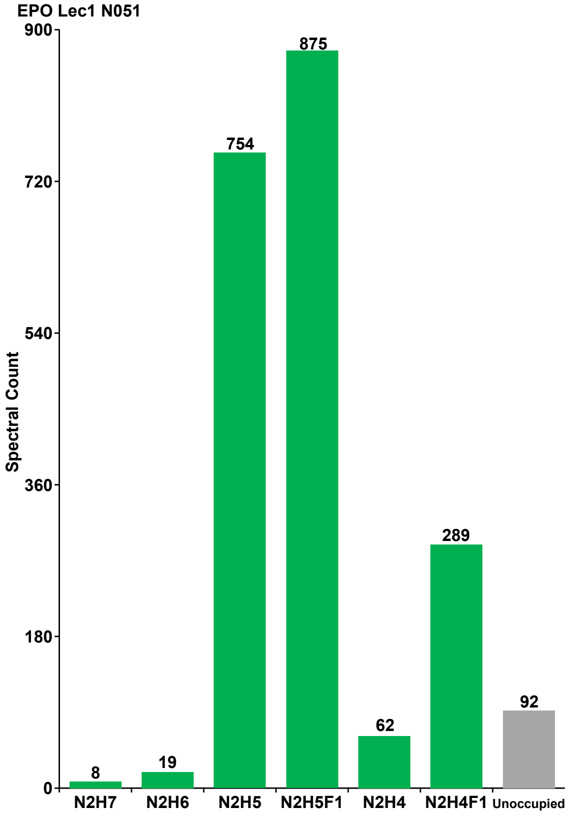

### S19: Site occupancy of erythropoietin sequon 2 (N065) expressed in HEK293F cells

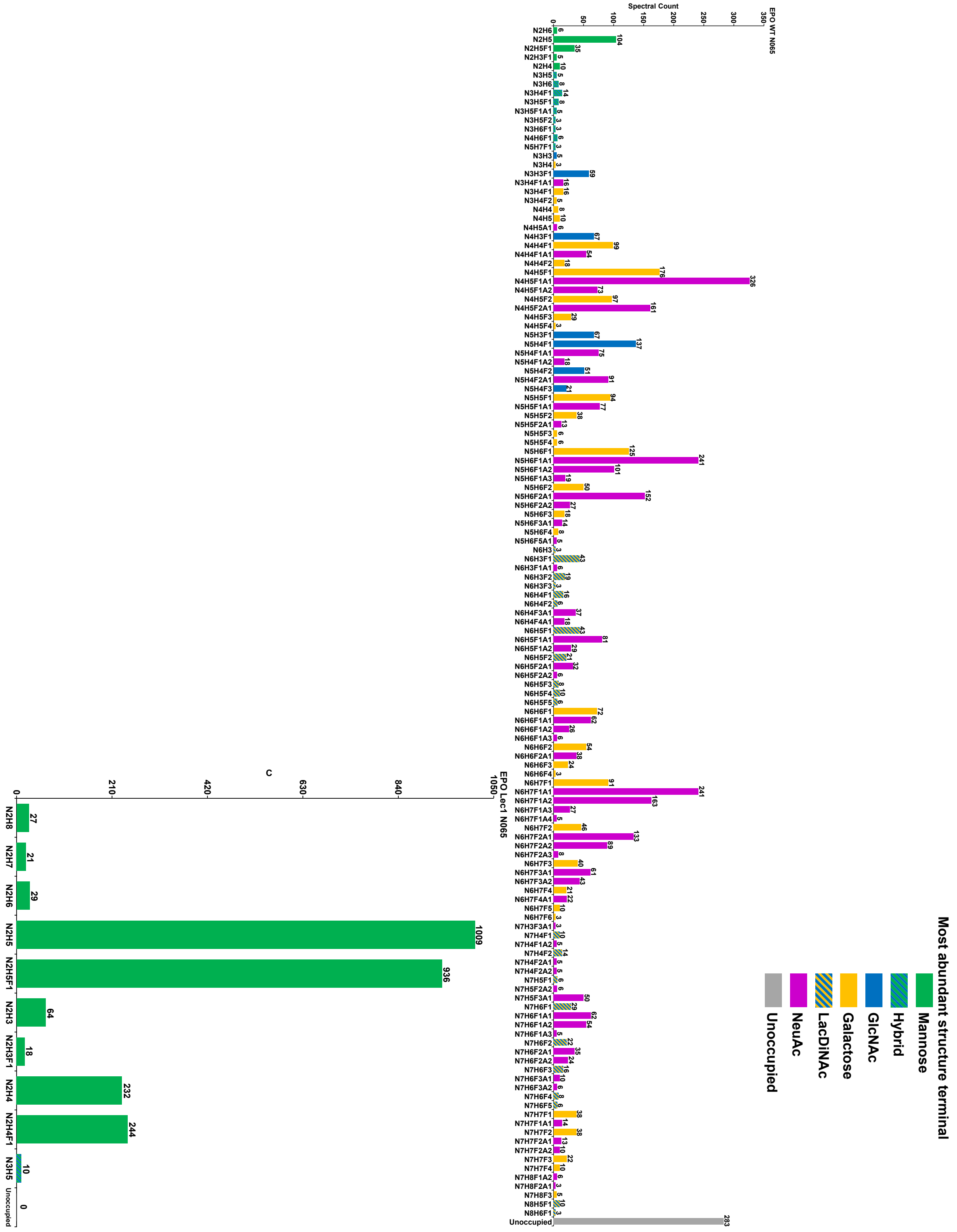

### S20: Site occupancy of erythropoietin sequon 3 (N110) expressed in HEK293F cells

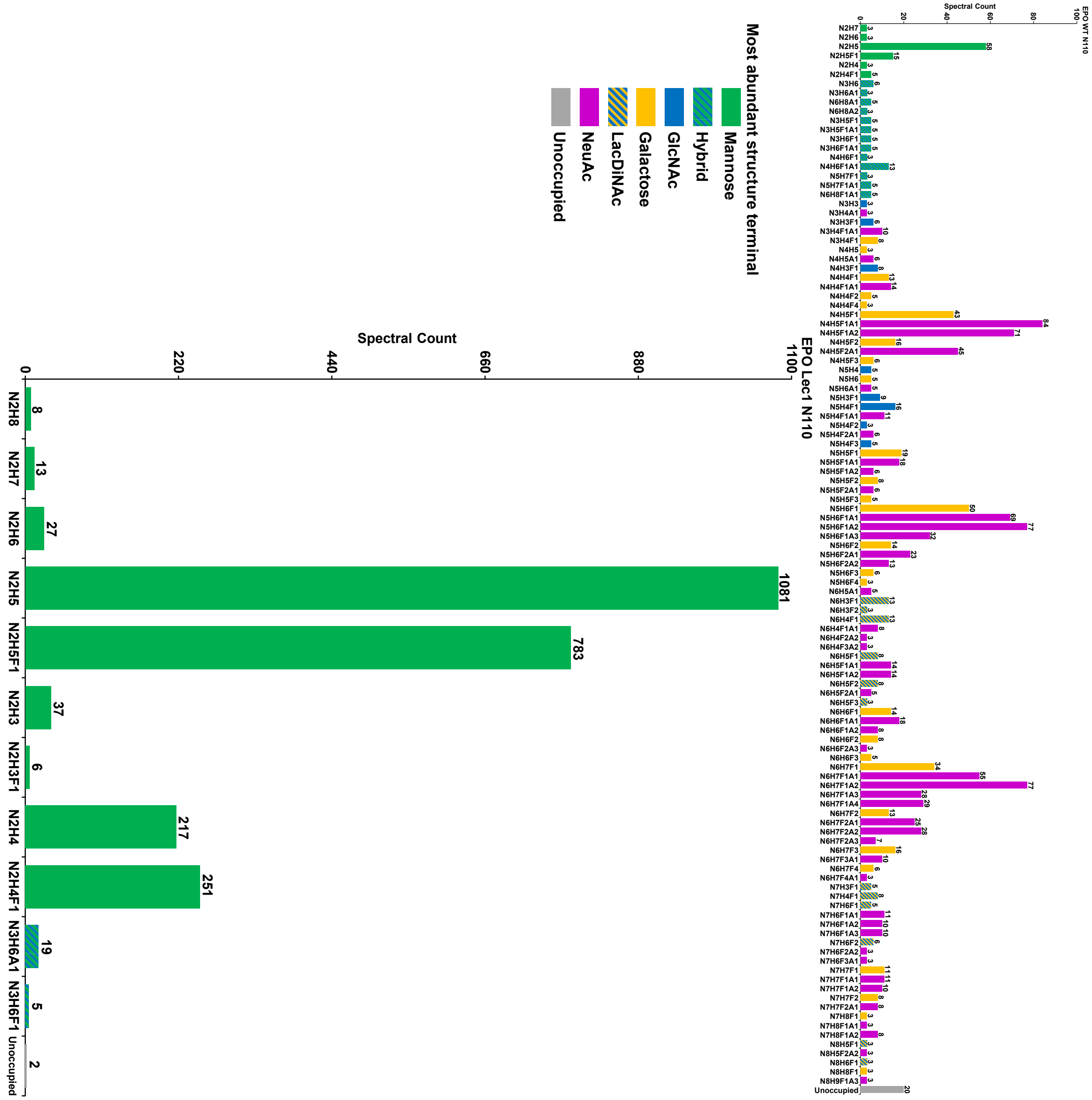

### S21: Site occupancy of SARS-CoV-2 spike glycoprotein sequon 1 (N0017) expressed in HEK293F cells

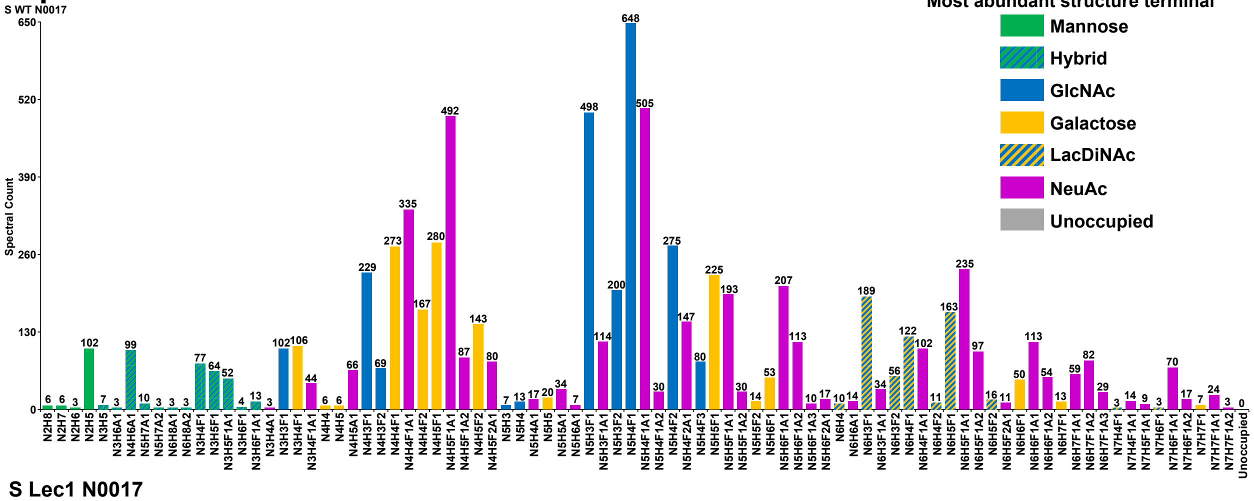


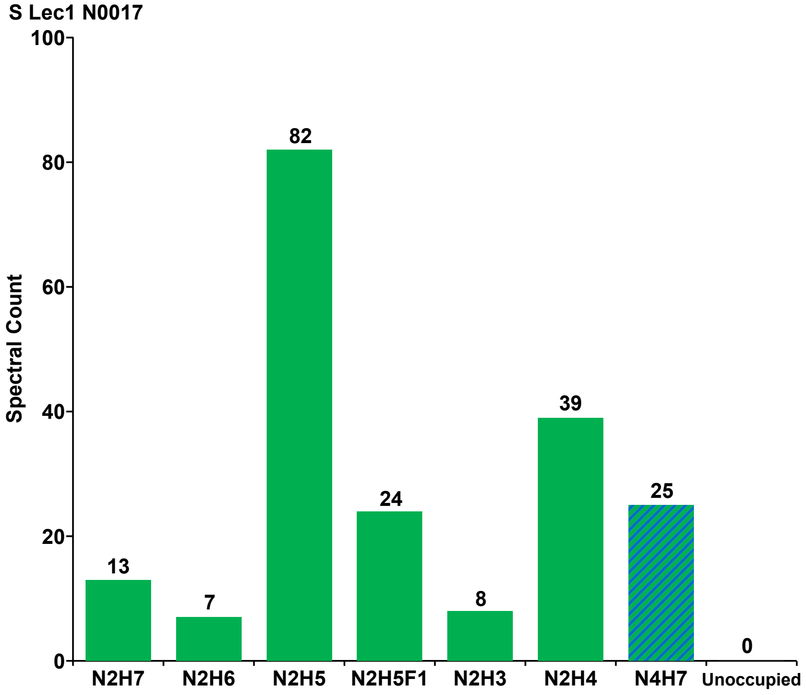

### S22: Site occupancy of SARS-CoV-2 spike glycoprotein sequon 2 (N0061) expressed in HEK293F cells

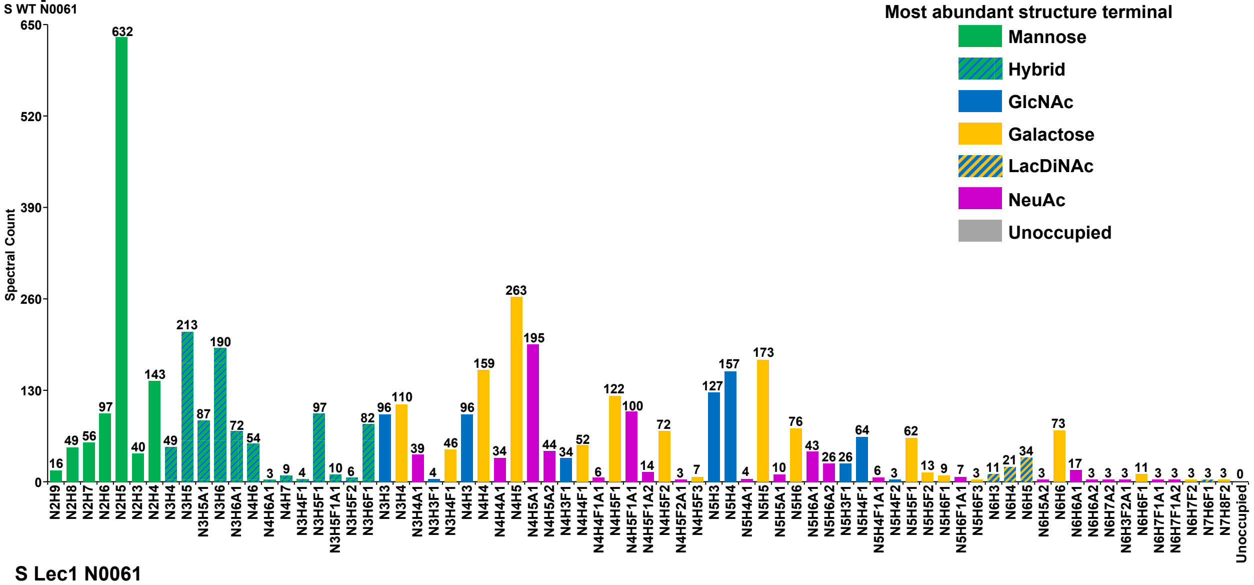


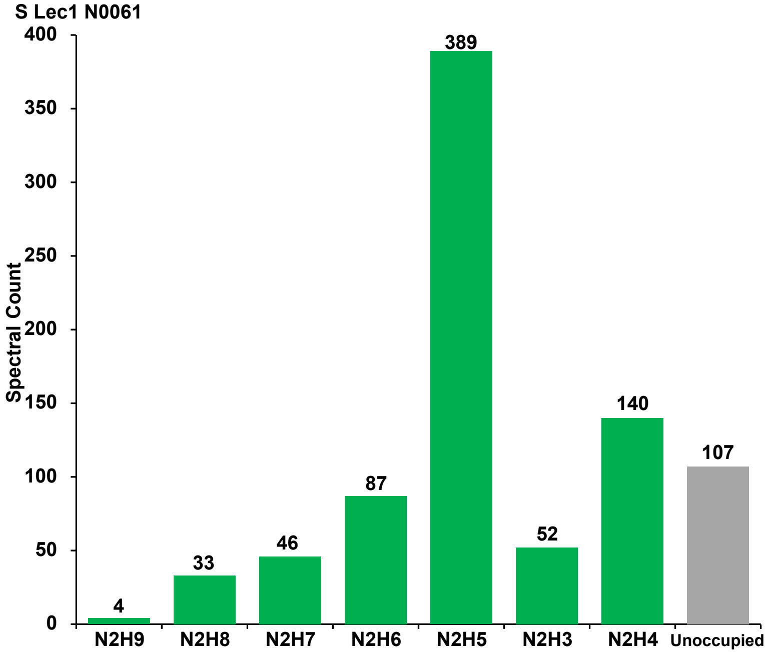

#### S23: Site occupancy of SARS-CoV-2 spike glycoprotein sequon 3 (N0074) expressed in HEK293F cells

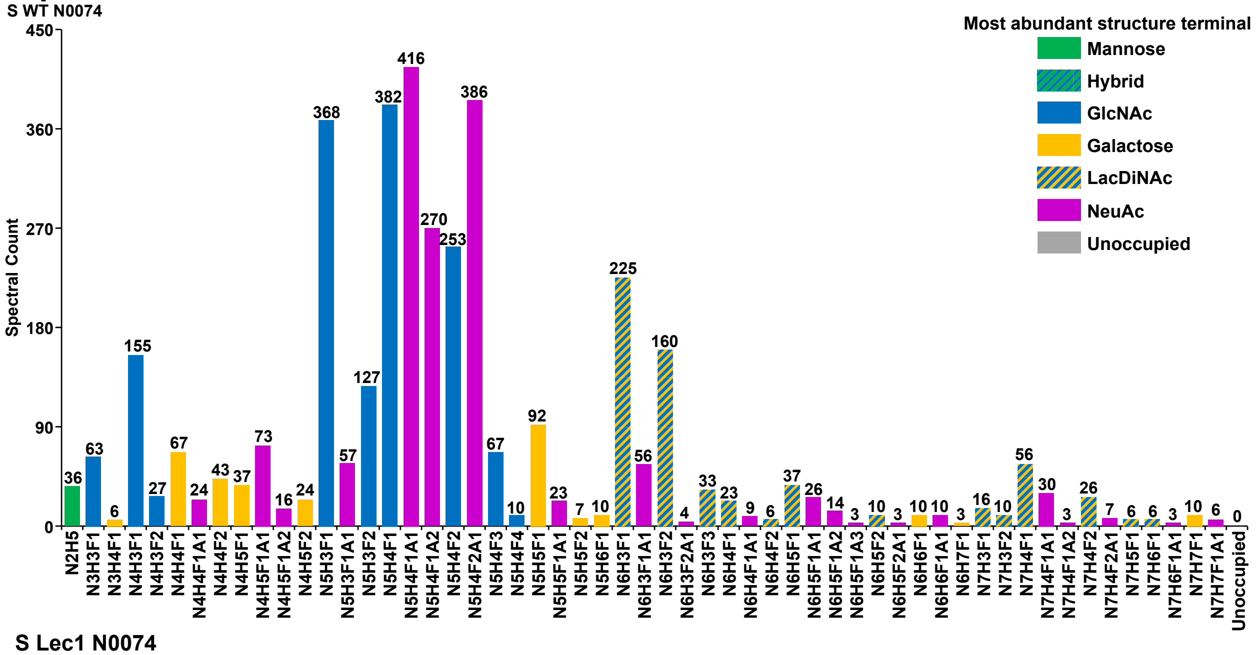


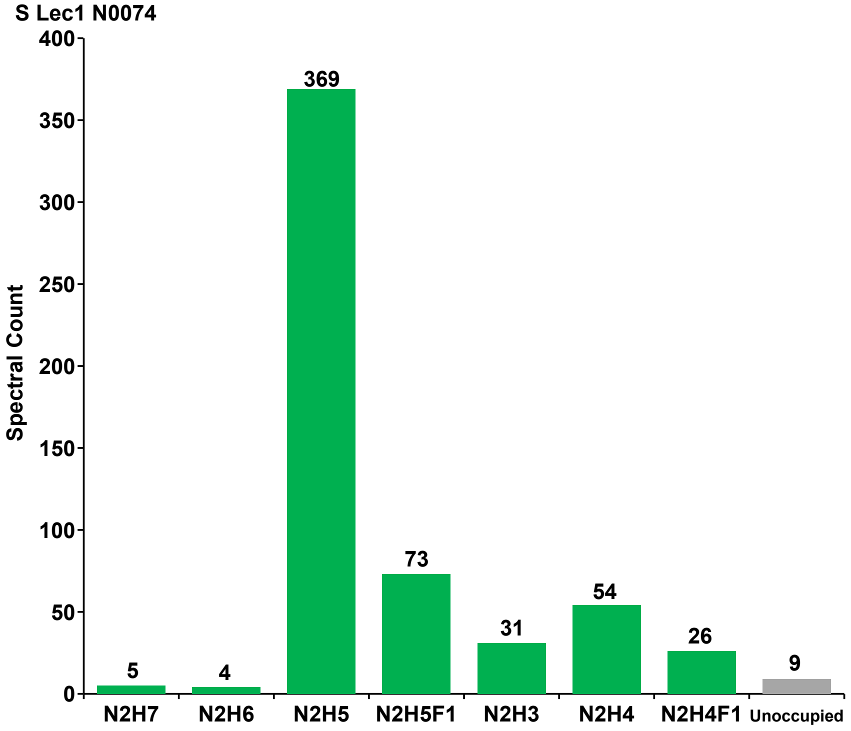

#### S24: Site occupancy of SARS-CoV-2 spike glycoprotein sequon 4 (N0122) expressed in HEK293F cells

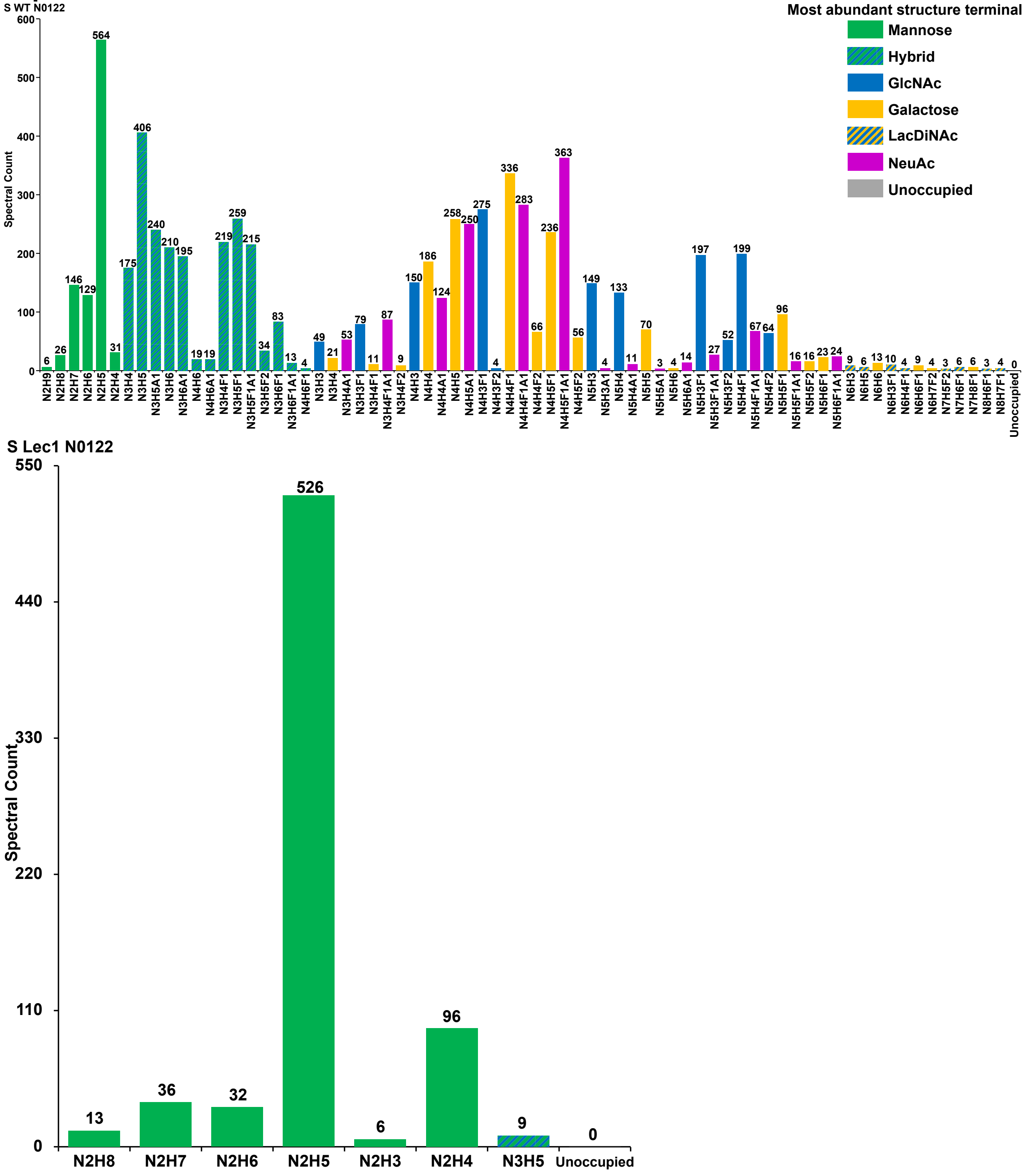

#### S25: Site occupancy of SARS-CoV-2 spike glycoprotein sequon 5 (N0149) expressed in HEK293F cells

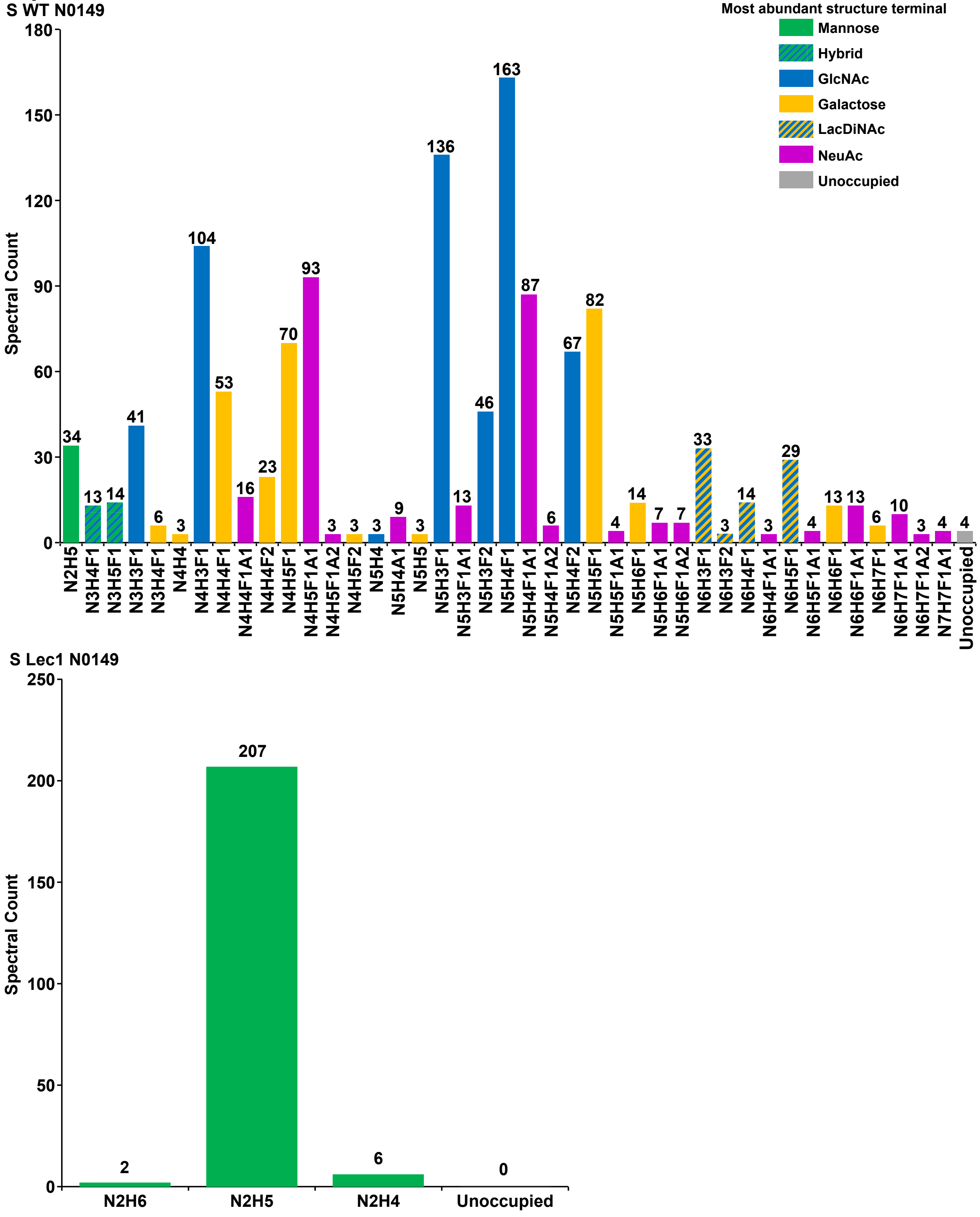

### S26: Site occupancy of SARS-CoV-2 spike glycoprotein sequon 6 (N0165) expressed in HEK293F cells

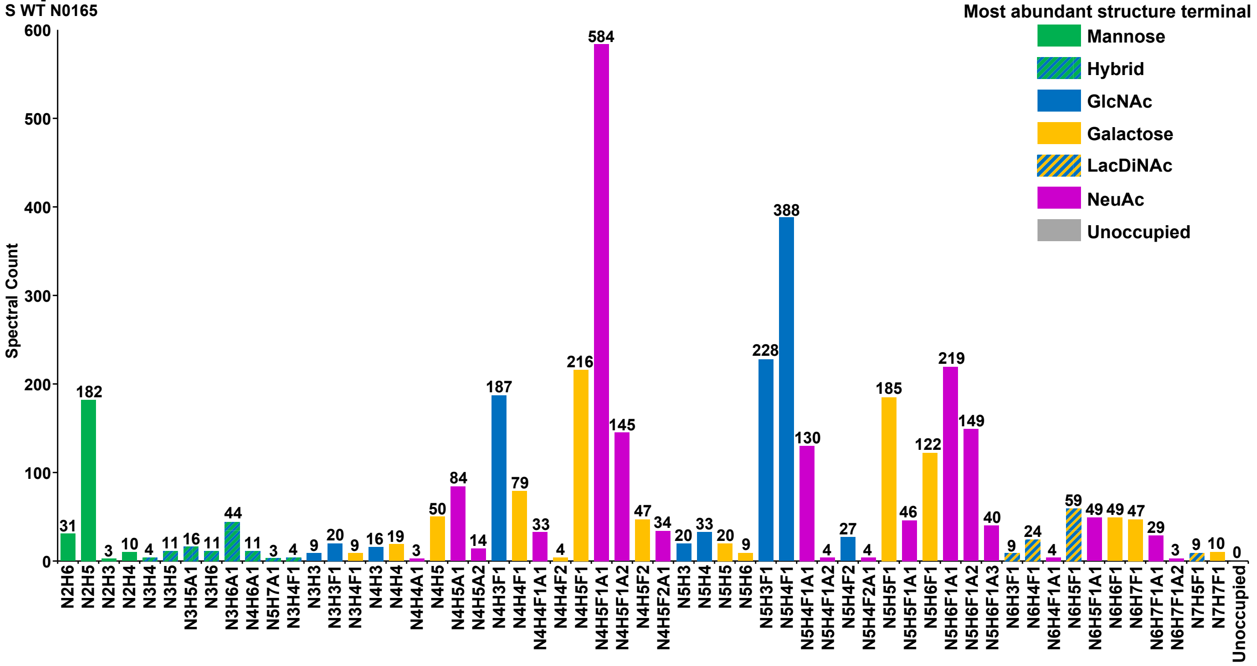


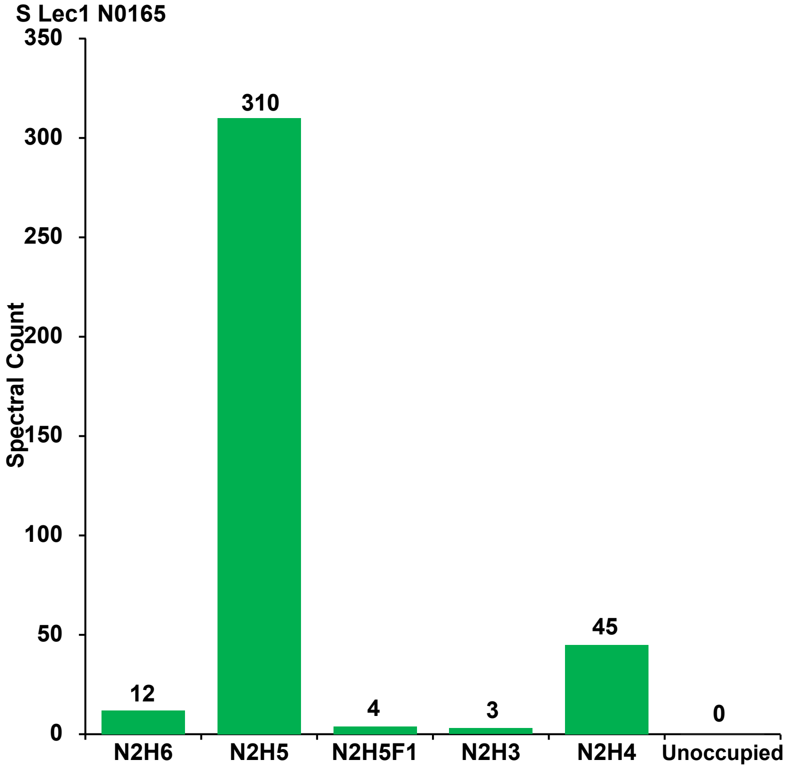

### S27: Site occupancy of SARS-CoV-2 spike glycoprotein sequon 7 (N0234) expressed in HEK293F cells

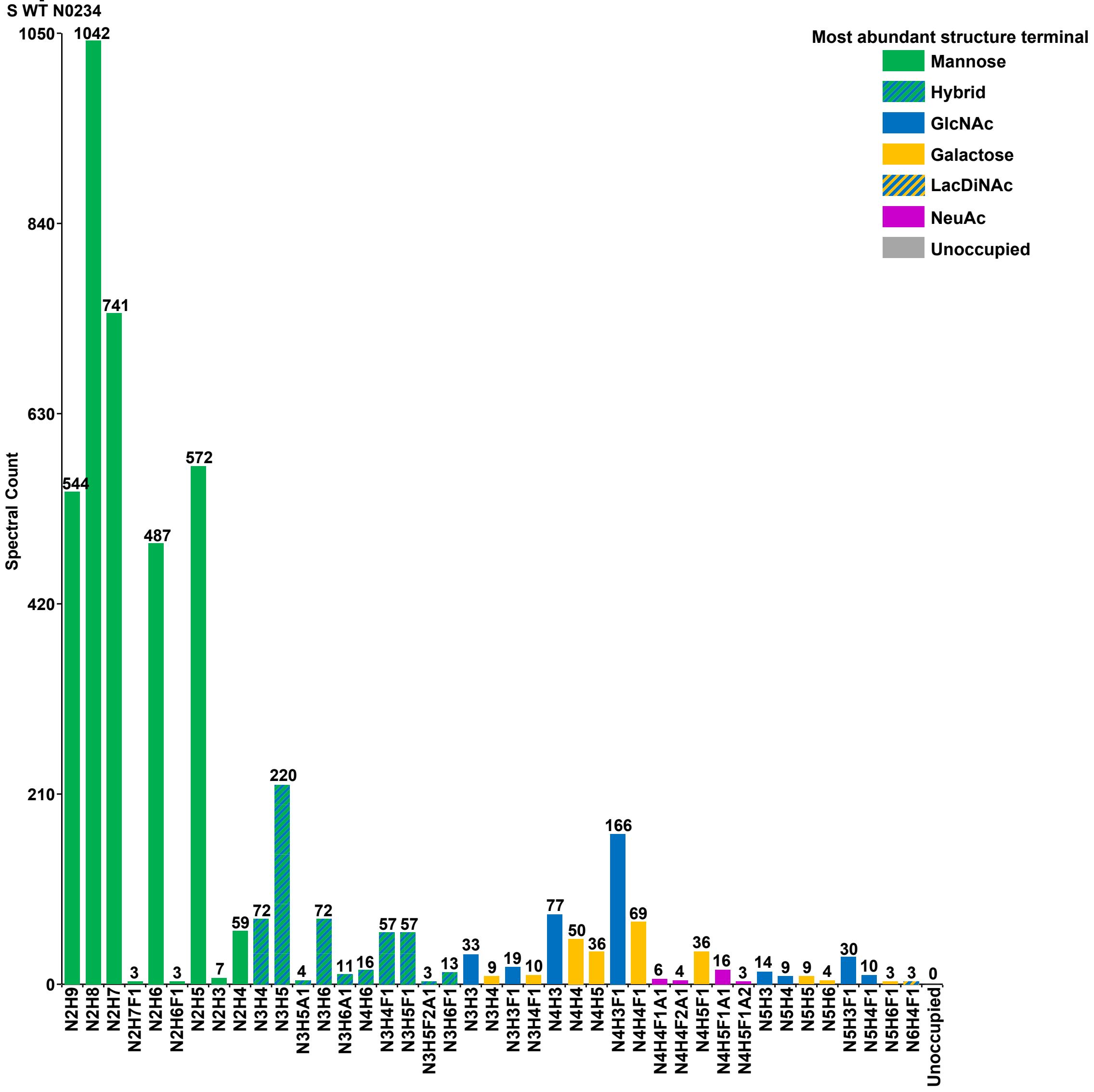


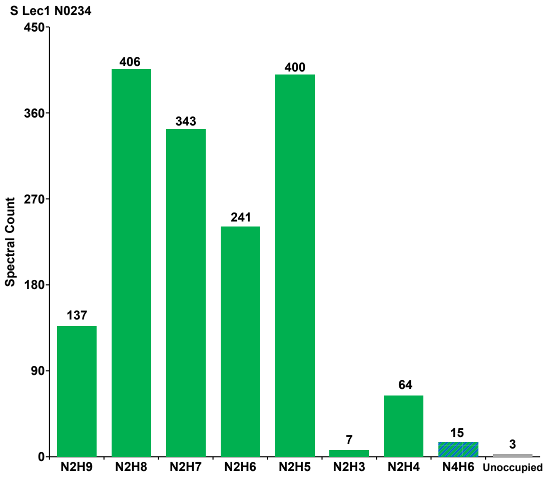

#### S28: Site occupancy of SARS-CoV-2 spike glycoprotein sequon 8 (N0282) expressed in HEK293F cells

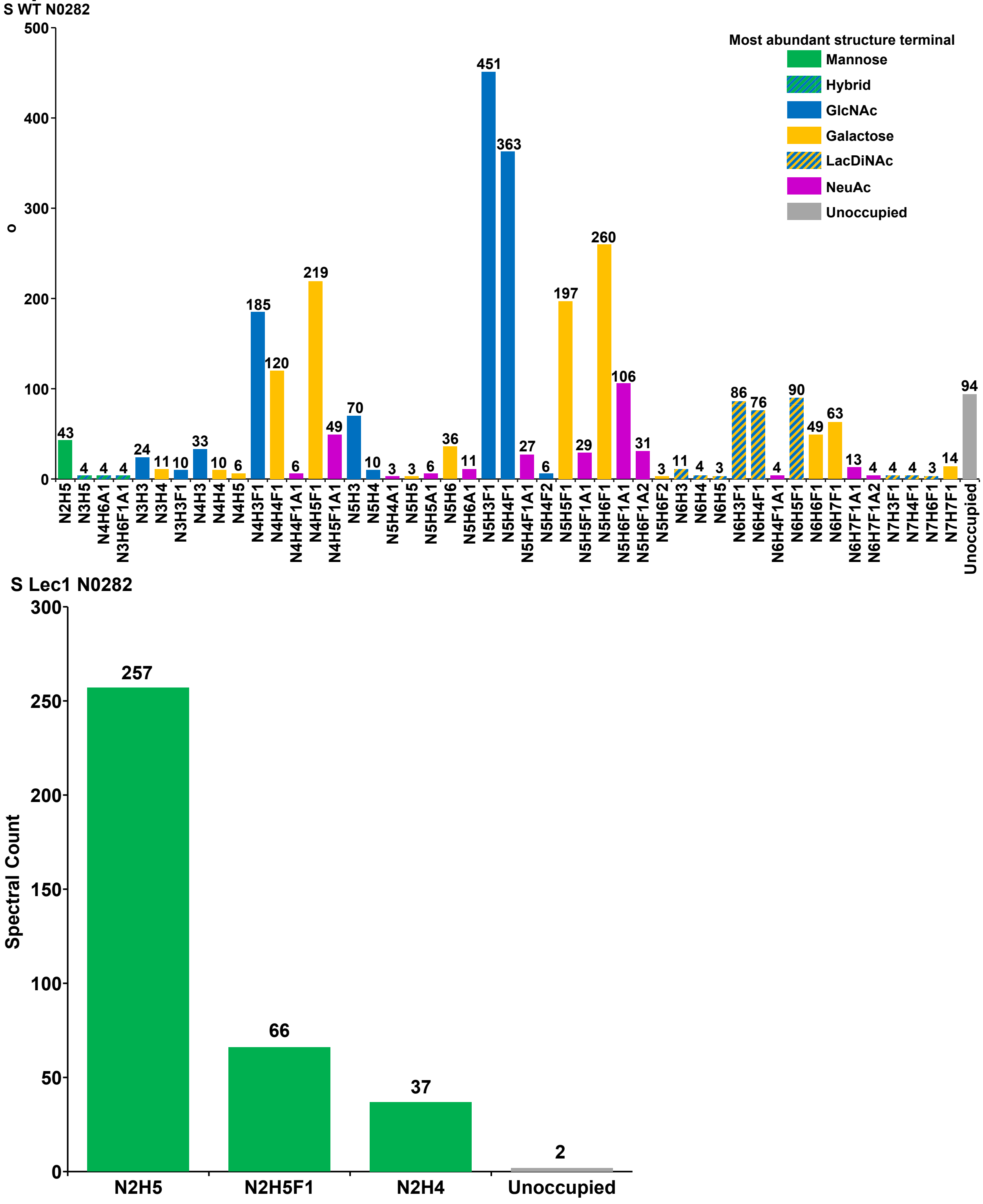

### S29: Site occupancy of SARS-CoV-2 spike glycoprotein sequon 9 (N0331) expressed in HEK293F cells

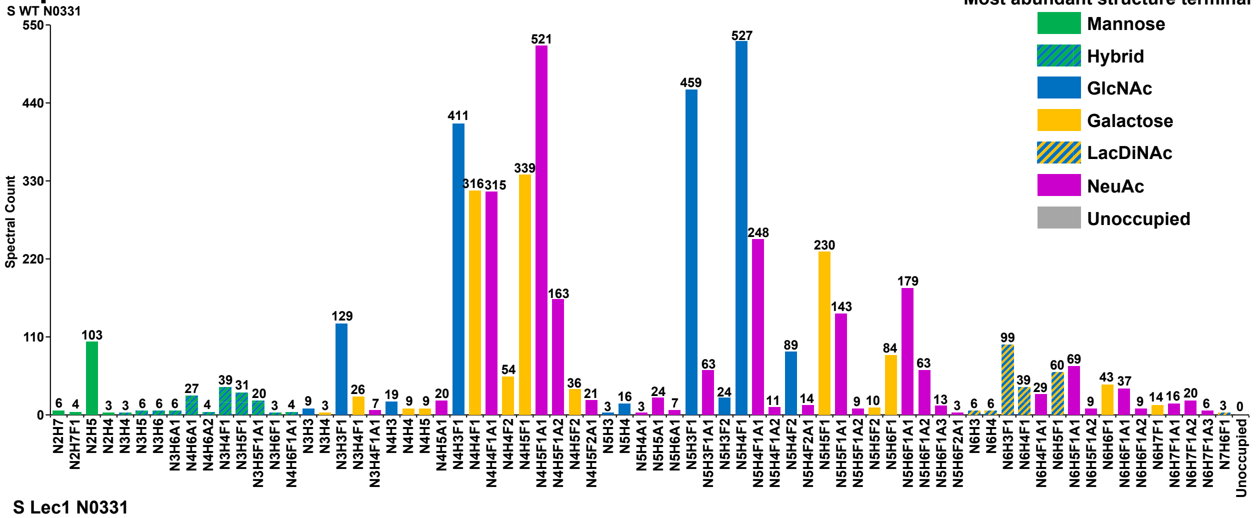


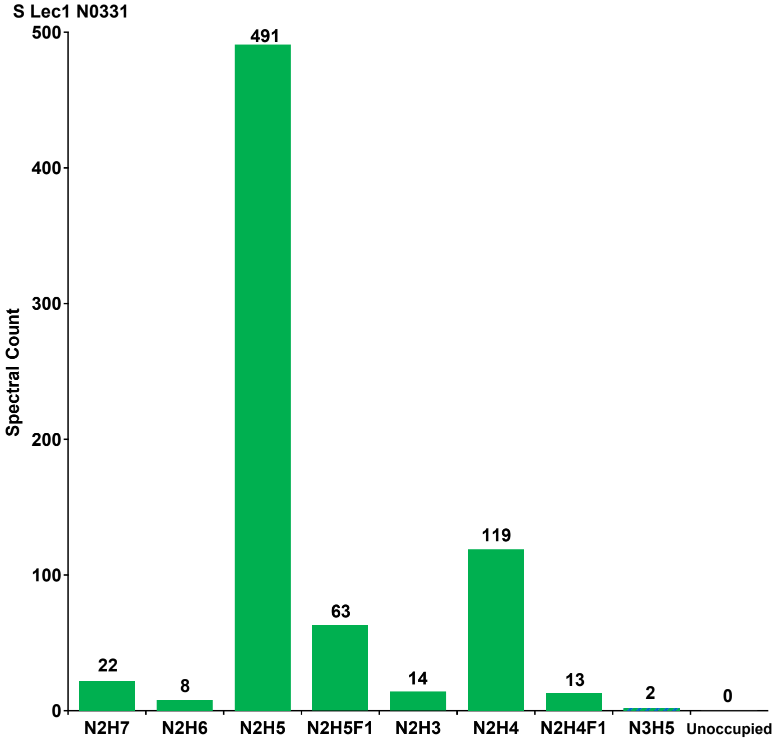

### S30: Site occupancy of SARS-CoV-2 spike glycoprotein sequon 10 (N0343) expressed in HEK293F cells

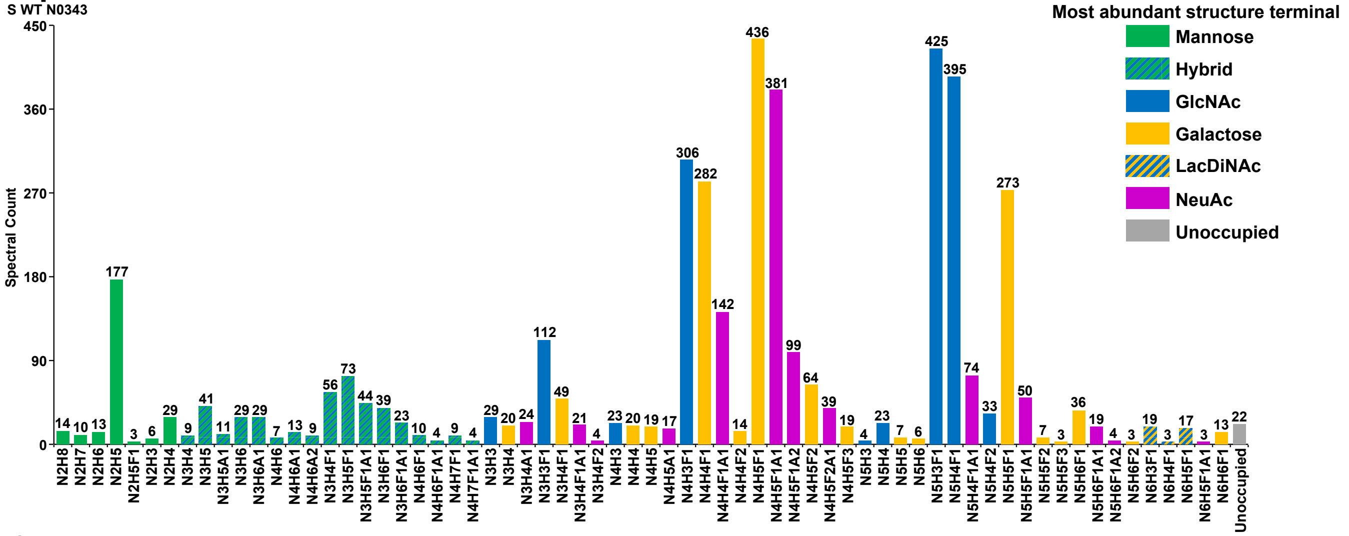


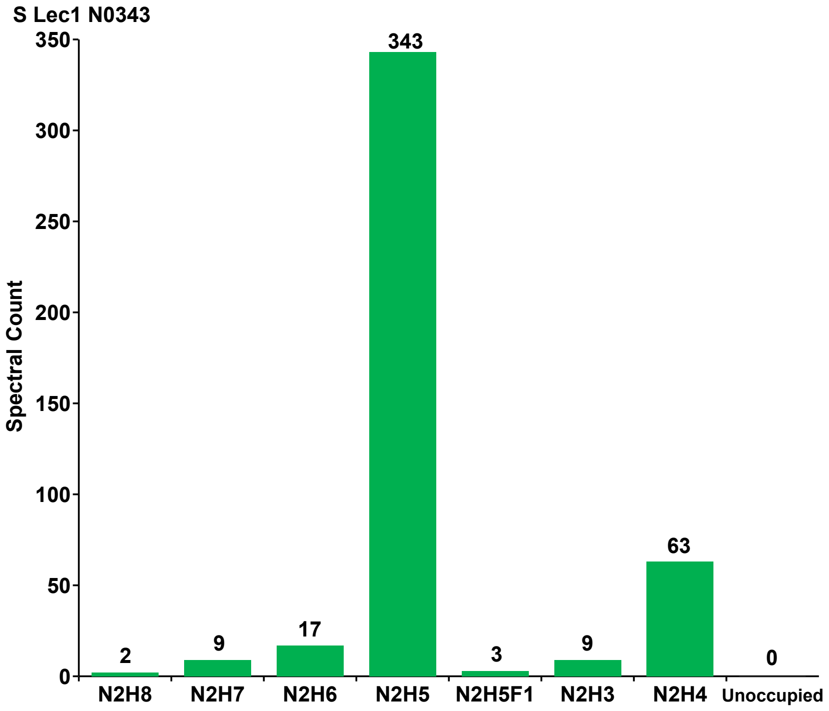

#### S31: Site occupancy of SARS-CoV-2 spike glycoprotein sequon 11 (N0603) expressed in HEK293F cells

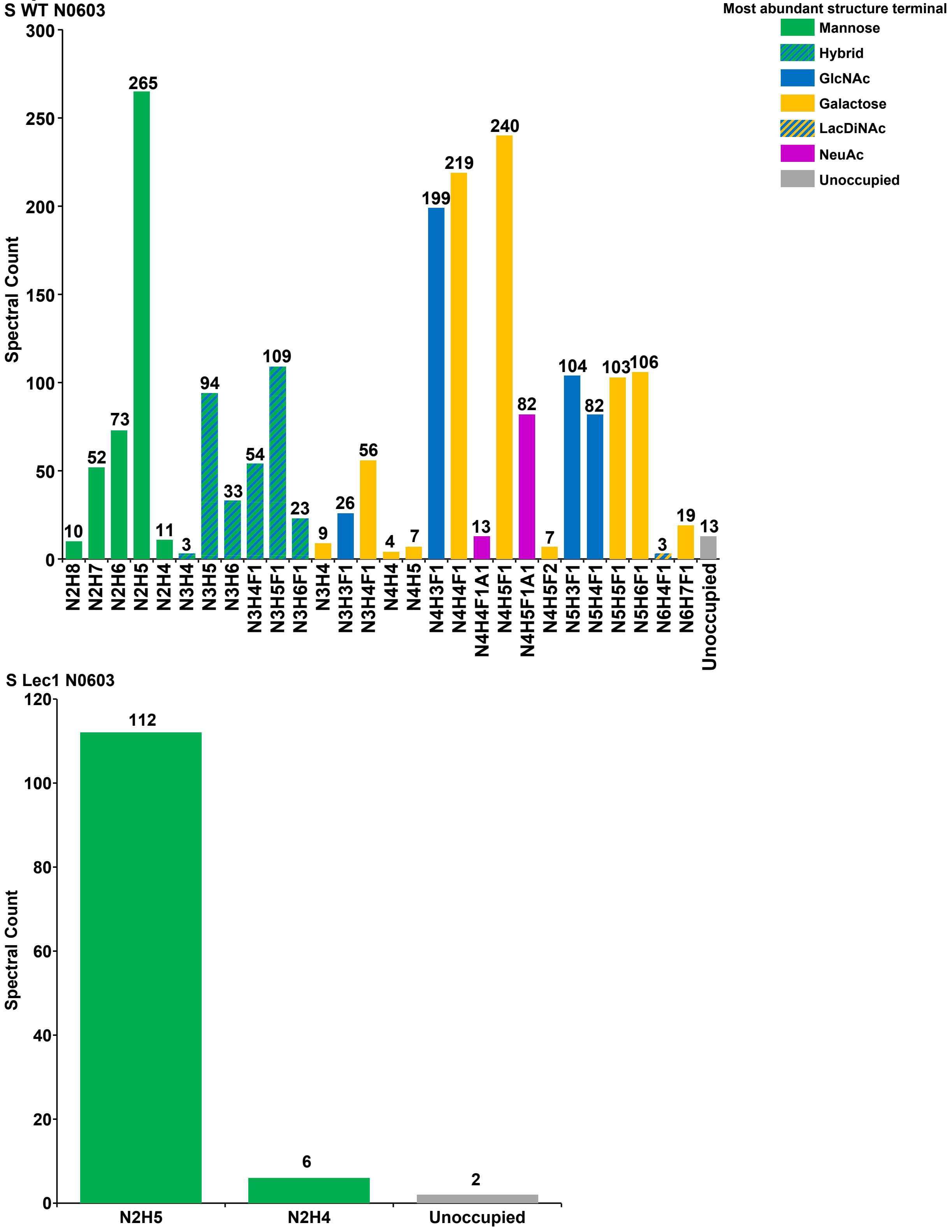

### S32: Site occupancy of SARS-CoV-2 spike glycoprotein sequon 12 (N0616) expressed in HEK293F cells

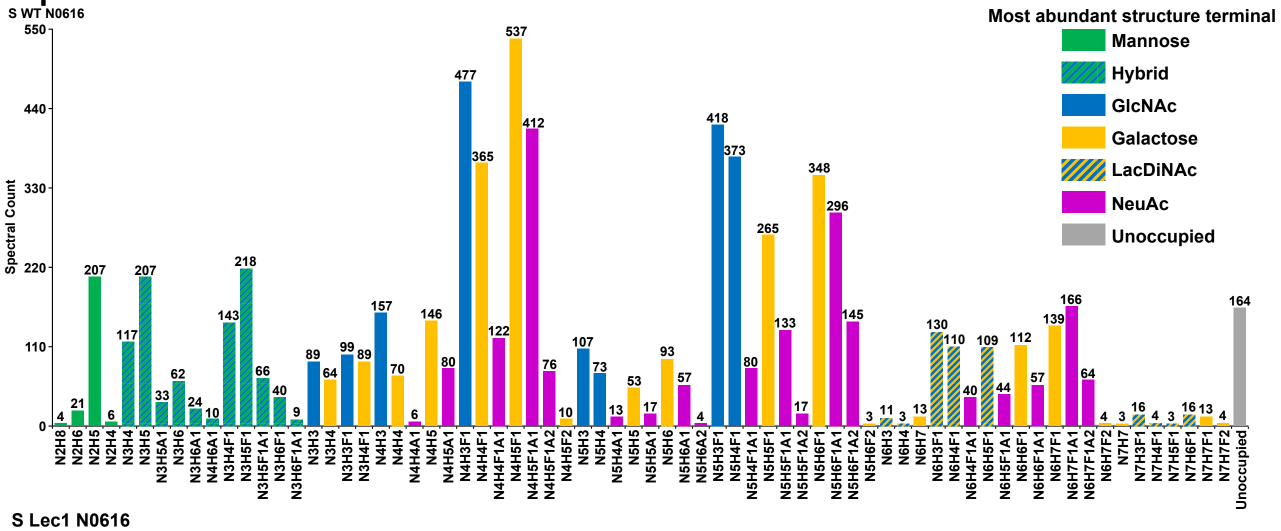


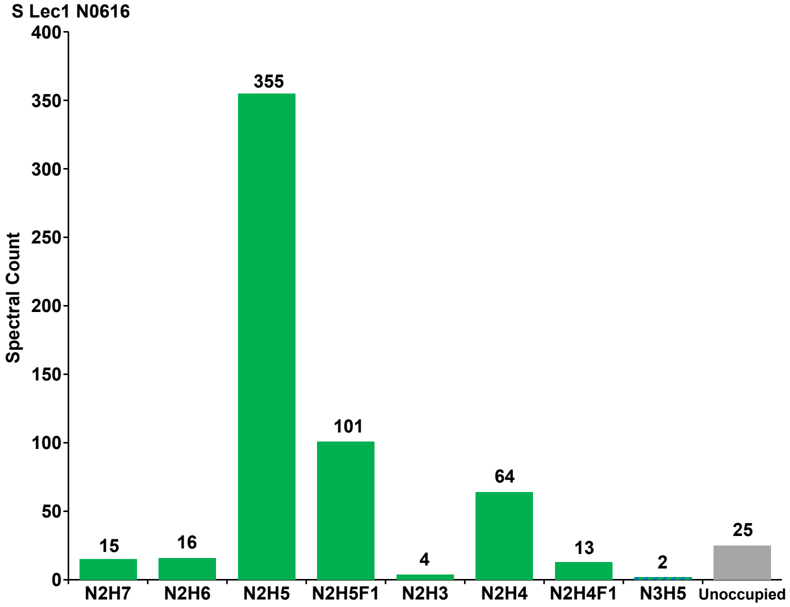

### S33: Site occupancy of SARS-CoV-2 spike glycoprotein sequon 13 (N0657) expressed in HEK293F cells

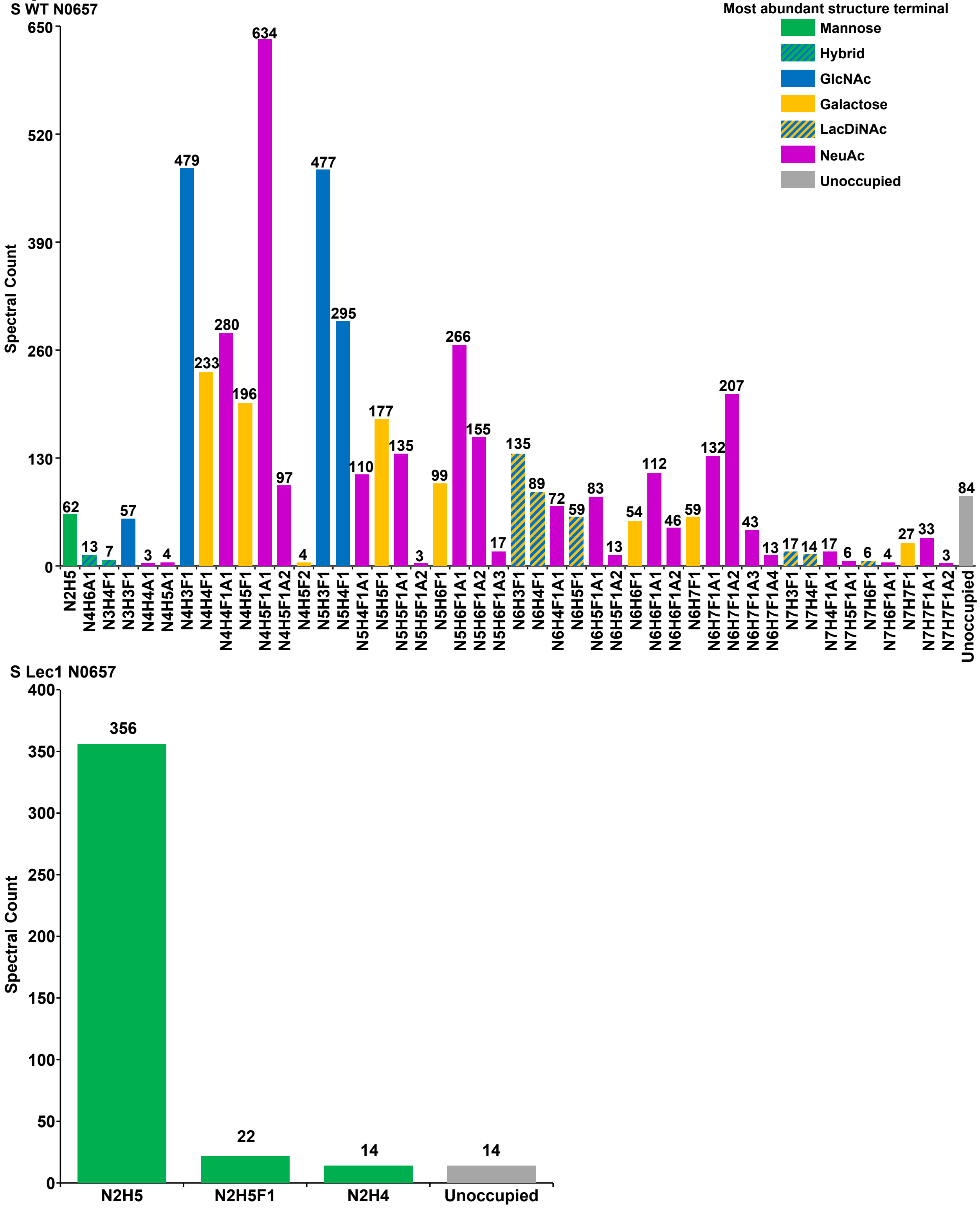

#### S34: Site occupancy of SARS-CoV-2 spike glycoprotein sequon 14 (N0709) expressed in HEK293F cells

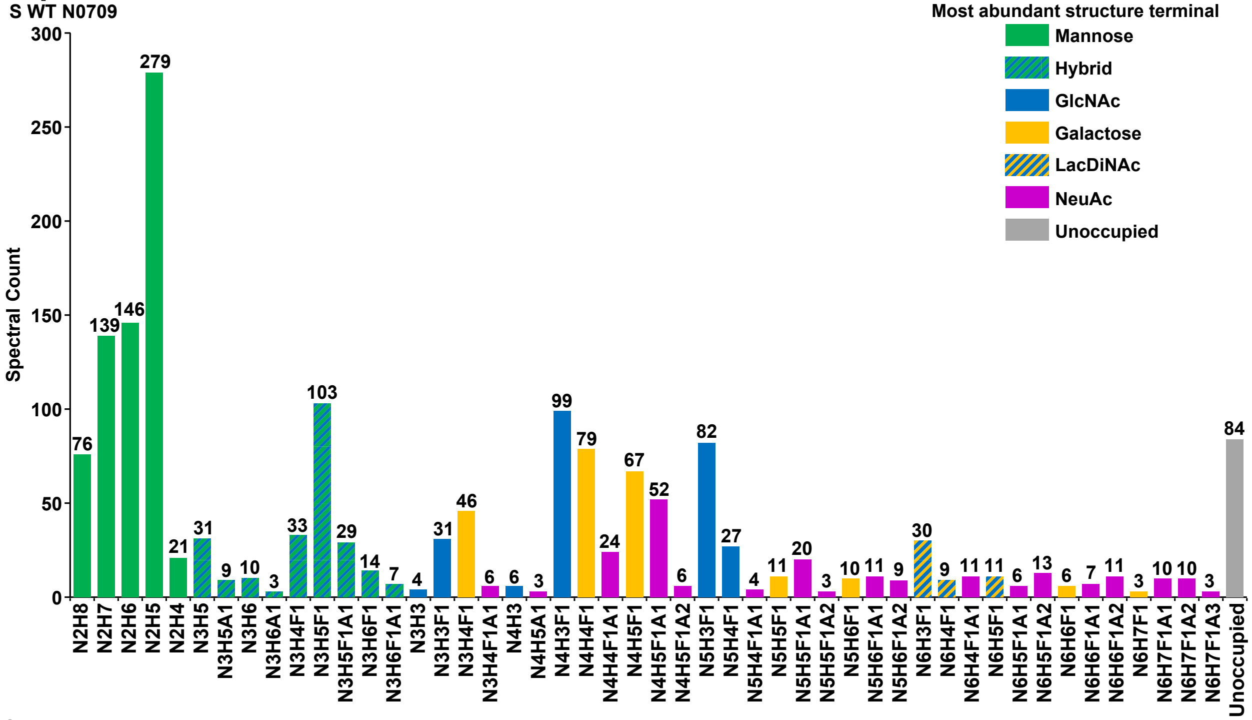


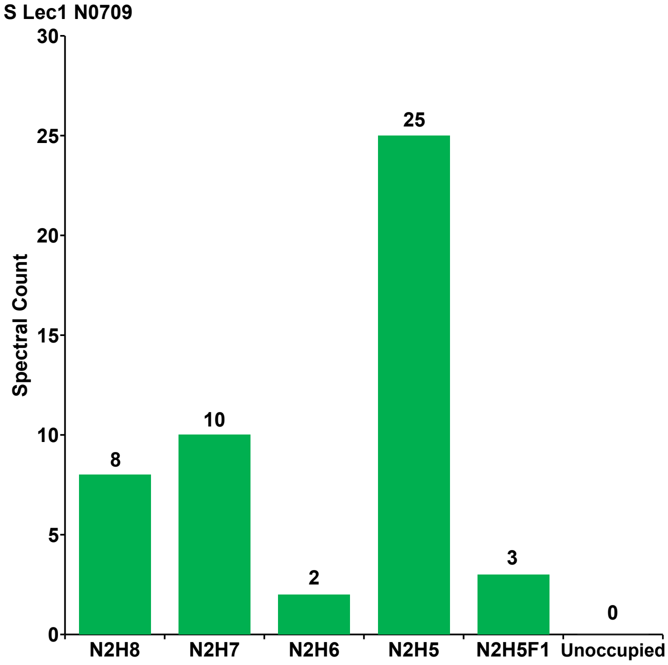

#### S35: Site occupancy of SARS-CoV-2 spike glycoprotein sequon 15 (N0717) expressed in HEK293F cells

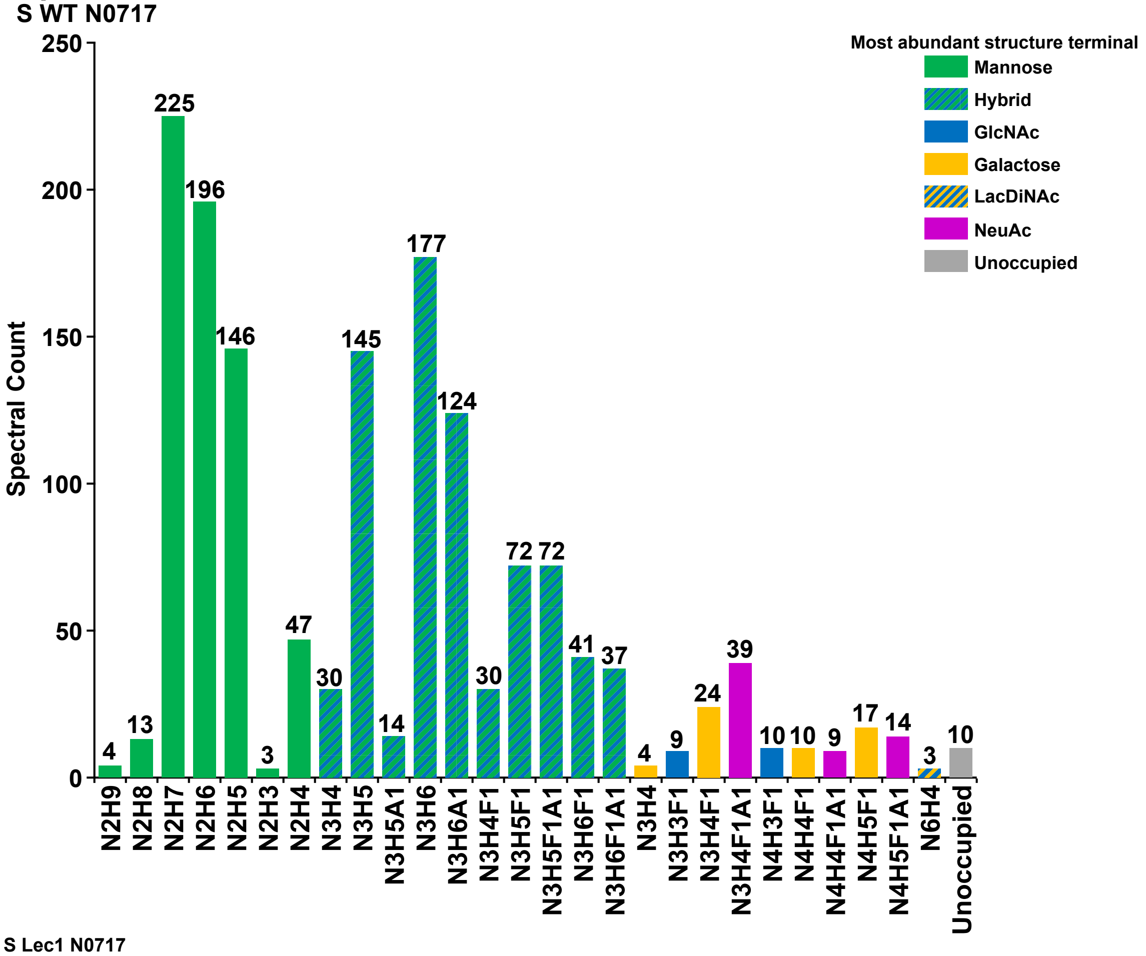


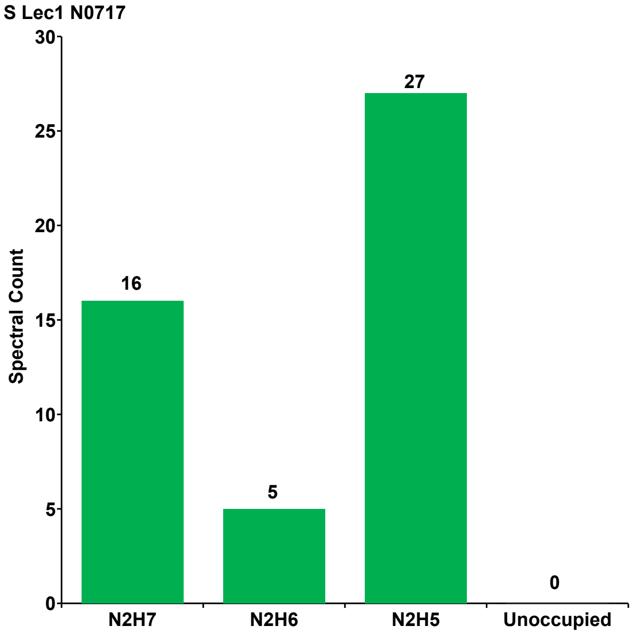

### S36: Site occupancy of SARS-CoV-2 spike glycoprotein sequon 16 (N0801) expressed in HEK293F cells

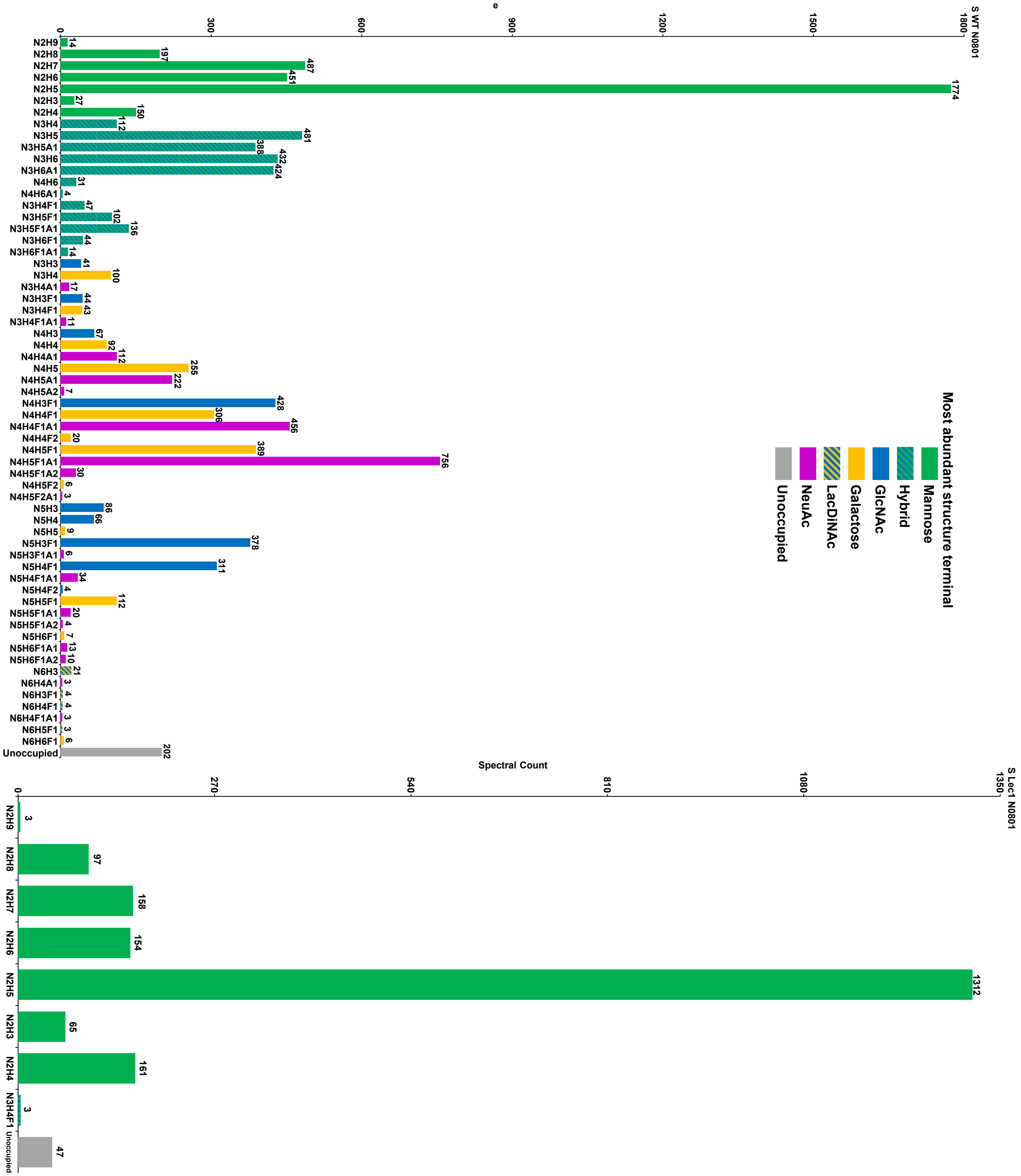

### S37: Site occupancy of SARS-CoV-2 spike glycoprotein sequon 17 (N1074) expressed in HEK293F cells

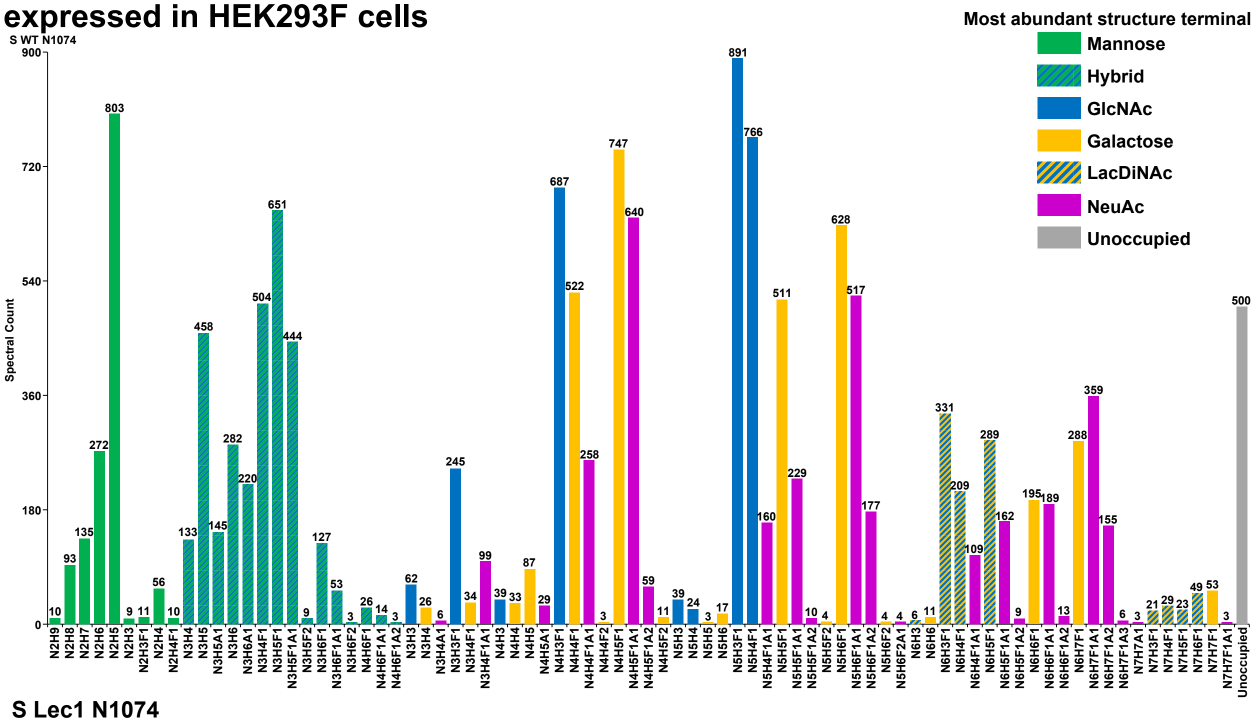


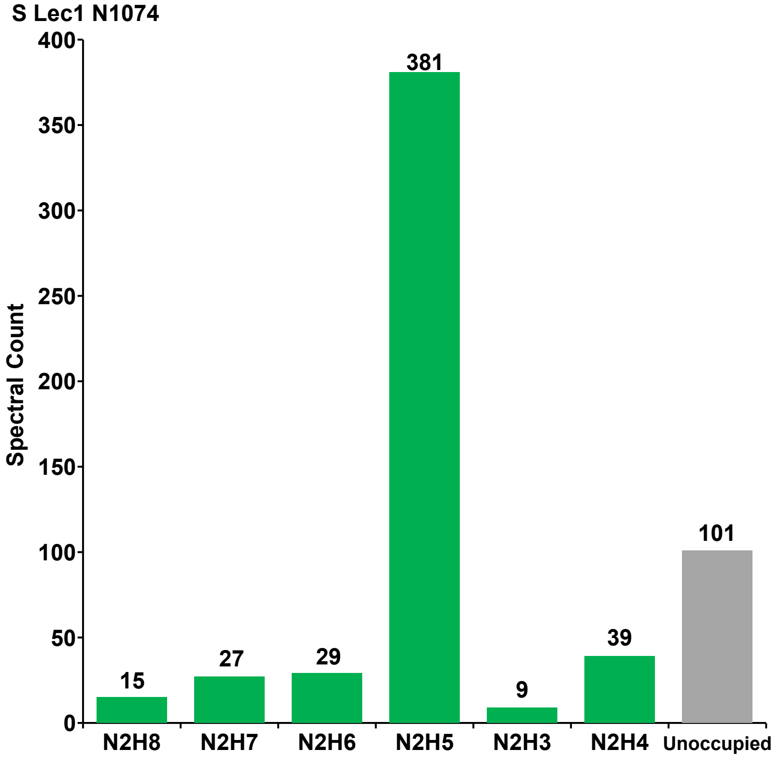

### S38: Site occupancy of SARS-CoV-2 spike glycoprotein sequon 18 (N1098) expressed in HEK293F cells

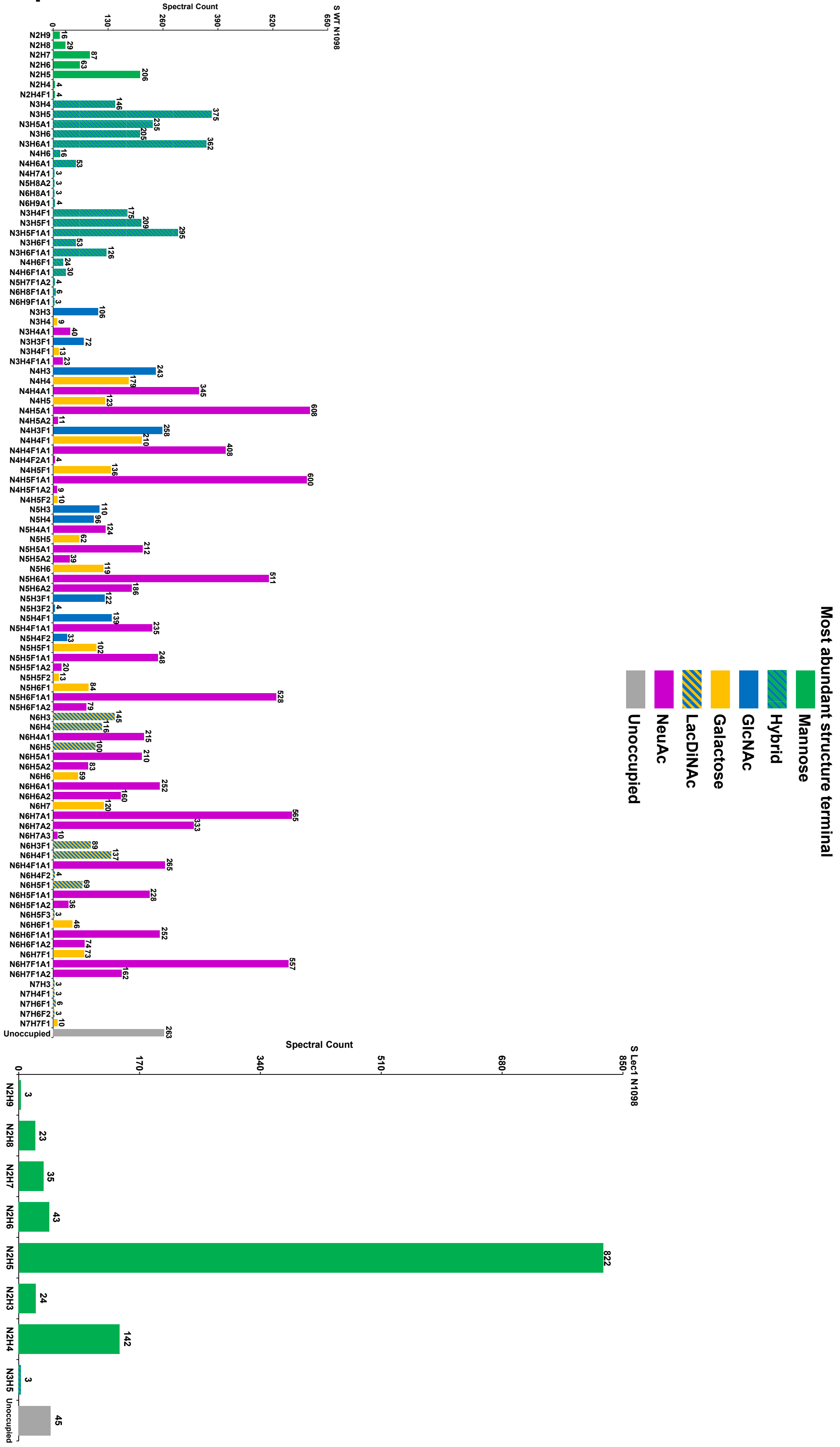

### S39: Site occupancy of SARS-CoV-2 spike glycoprotein sequon 19 (N1134) expressed in HEK293F cells

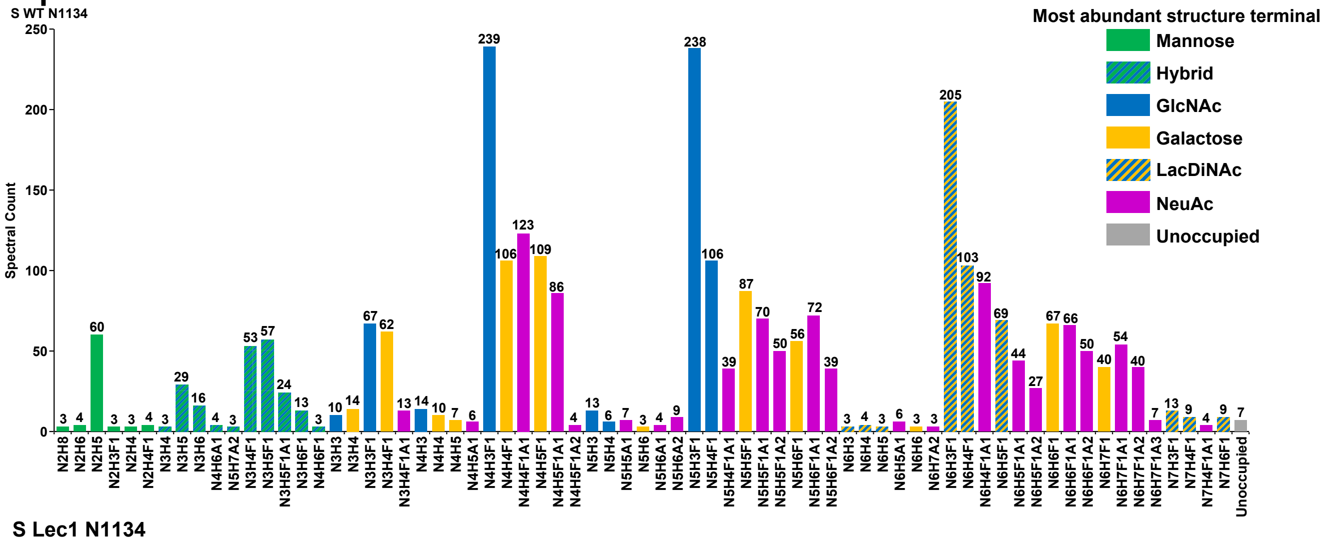


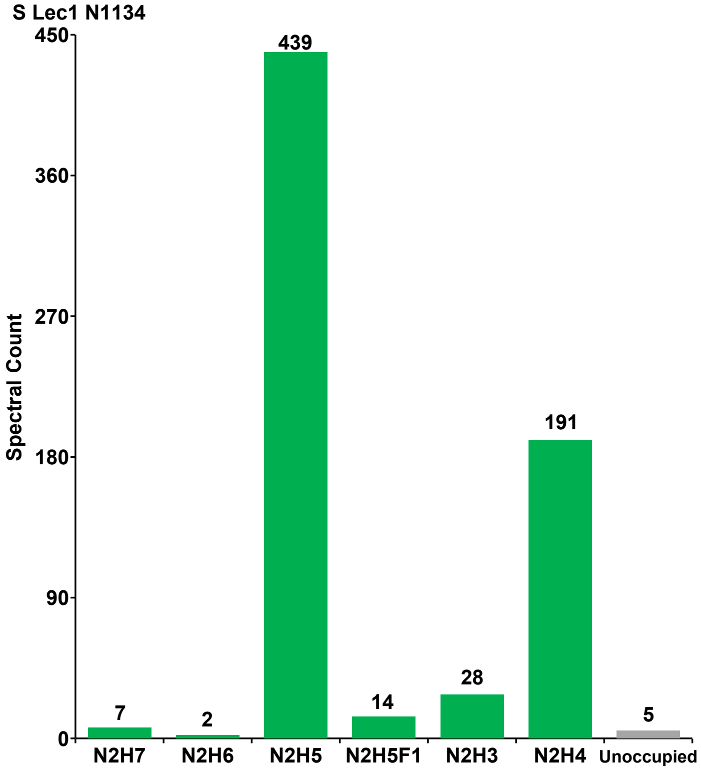

#### S40: Site occupancy of SARS-CoV-2 spike glycoprotein sequon 20 (N1158) expressed in HEK293F cells

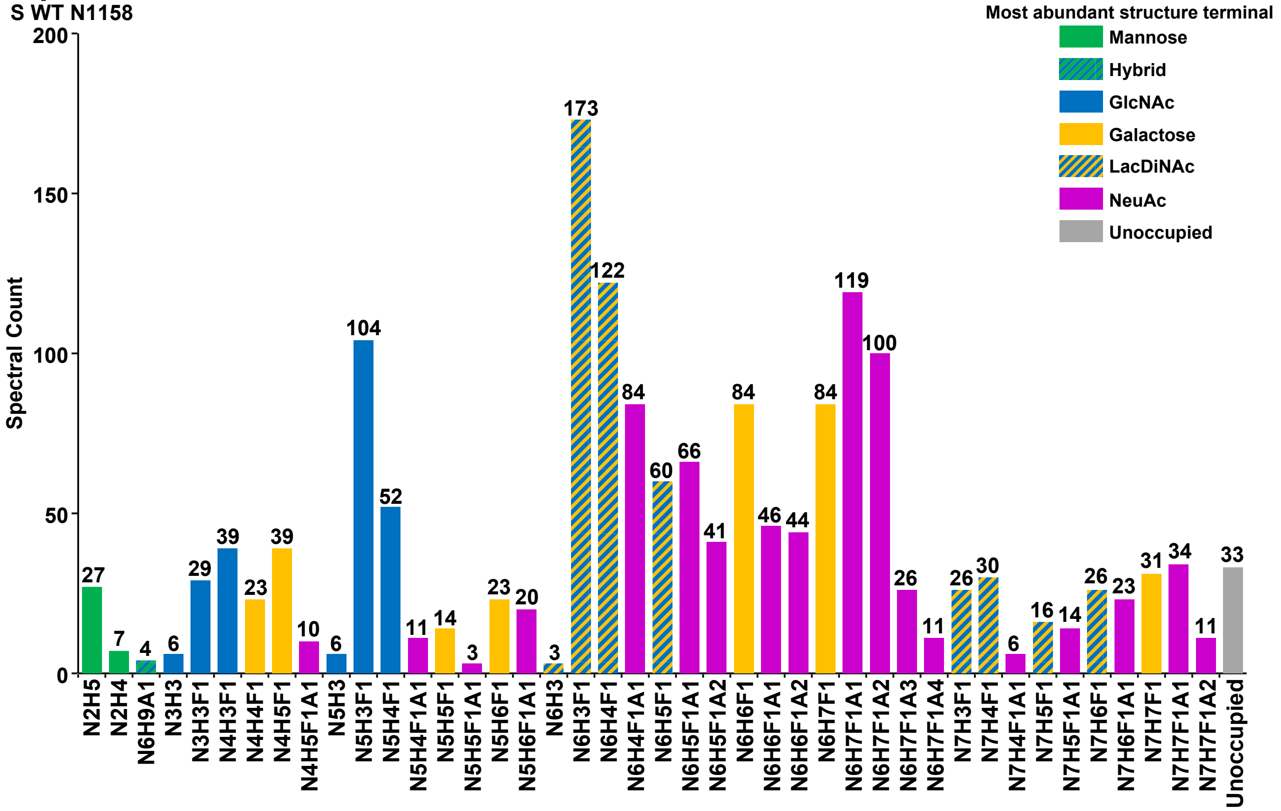


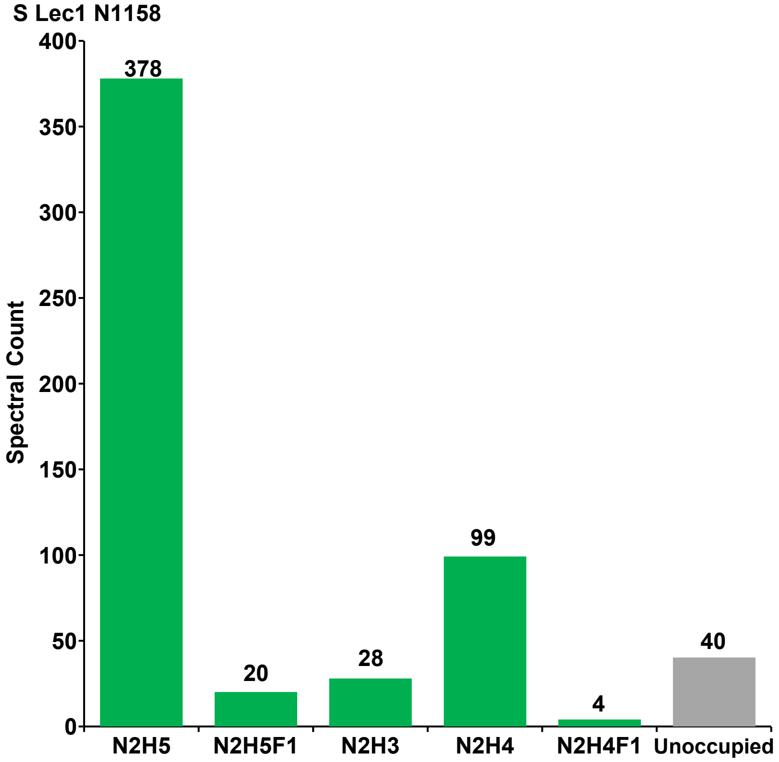

### S41: Site occupancy of SARS-CoV-2 spike glycoprotein sequon 21 (N1173) expressed in HEK293F cells

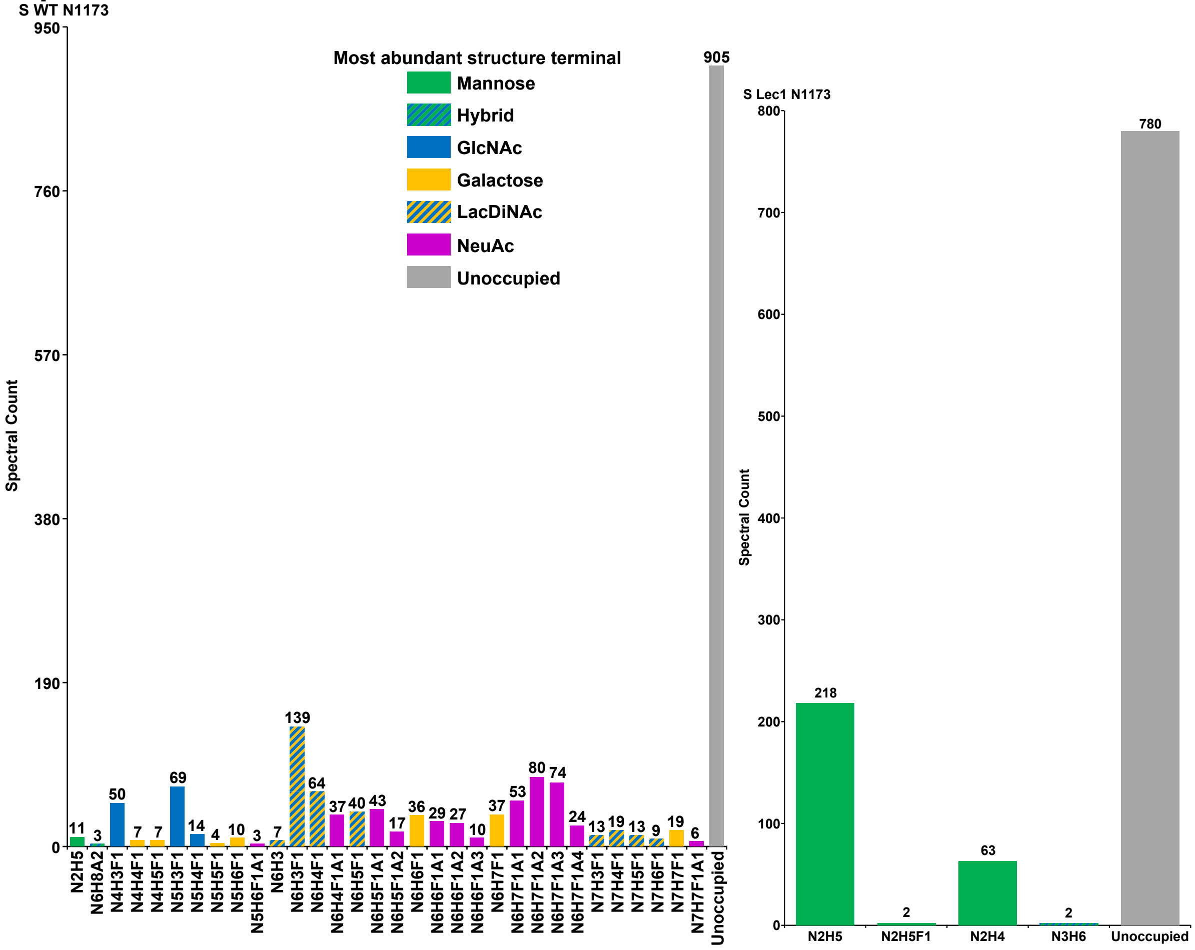

#### S42: Site occupancy of SARS-CoV-2 spike glycoprotein sequon 22 (N1194) expressed in HEK293F cells

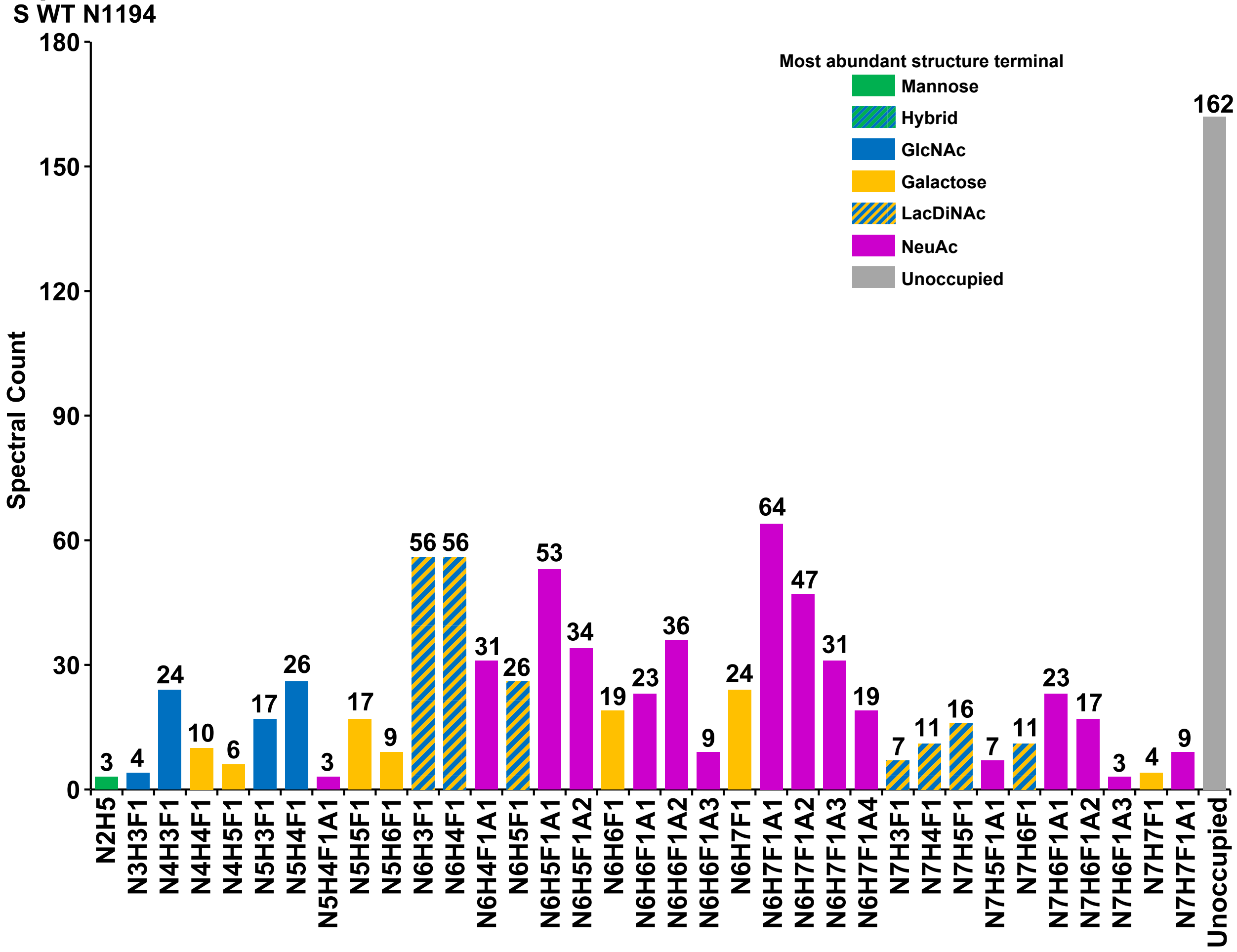


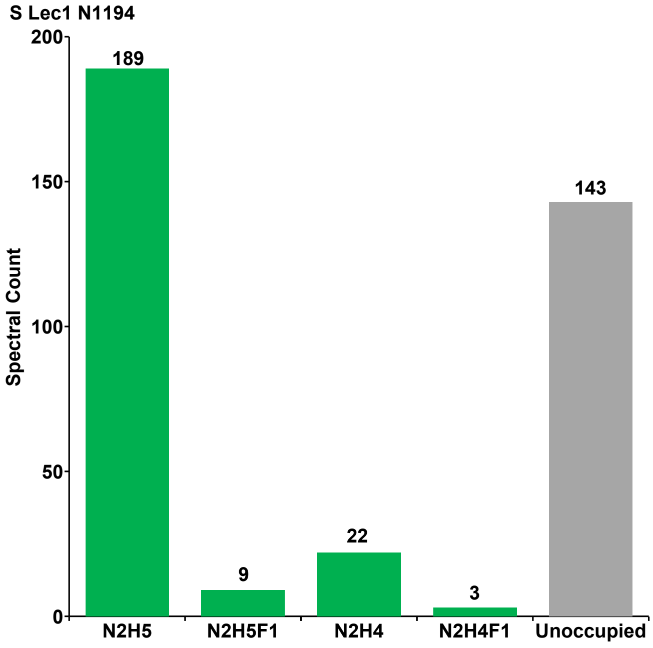
